# Efficient accumulation of new irregular monoterpene malonyl glucosides in *Nicotiana benthamiana* achieved by co-expression of isoprenyl diphosphate synthases and substrate-producing enzymes

**DOI:** 10.1101/2025.08.06.668877

**Authors:** Iryna Gerasymenko, Yuriy V. Sheludko, Volker Schmidts, Heribert Warzecha

## Abstract

Irregular monoterpenes have limited natural sources but possess unique activities applicable in medicine and agriculture. To enable sustainable plant-based production of these compounds, we established a transient expression procedure to enhance the biosynthetic flux in *Nicotiana benthamiana* toward dimethylallyl diphosphate (DMAPP), a substrate for isopentenyl diphosphate synthases (IDSs) that generate irregular monoterpene skeletons. Considering the benefits of glycosylation for accumulating and storing monoterpenes in extractable form, we focused on developing a platform for production of non-volatile glycosylated irregular monoterpenes using three IDS that form branched and cyclic structures. The analysis of methanolic leaf extracts from transiently transformed *N. benthamiana* plants revealed six major new components, 6*-O-*malonyl*-β-*D-glucopyranoside and 6*-O-*malonyl*-β-*D-glucopyranosyl-(1→2)-*β*-D-glucopyranoside derivatives of chrysanthemol, lavandulol and cyclolavandulol, five of which are novel compounds. Alleviating two bottlenecks in the DMAPP formation in plastids by co-expressing 1-deoxyxylulose 5-phosphate synthase and isopentenyl diphosphate isomerases increased the yield of chrysantemyl and lavandulyl glucosides produced by plant-derived IDS to 1.7 ± 0.4 μmol g^-1^ FW and 1.4 ± 0.3 μmol g^-1^ FW, respectively. A bacterial cyclolavandulyl diphosphate synthase operated efficiently in chloroplasts and cytoplasm. The highest irregular monoterpene concentrations were achieved in cytoplasm by co-expression of hydroxymethylglutaryl-CoA reductase, the bottleneck enzyme of the mevalonate pathway for DMAPP biosynthesis. The mean level of cyclolavandulyl glucosides reached 3.9 ± 1.5 μmol g^-1^ FW; the top-performing plants contained 6.6 μmol g^-1^ FW. This yield represents the highest amount of irregular monoterpene glycosides produced in plant systems.

## 1. Introduction

Terpenoids form the largest group of natural products, with over 80 000 different structures identified (Christianson, 2017). Approximately three-quarters of all known terpenes have been isolated from plants (Li et al., 2024), where they serve various functions, including defensive, communicative, and stress-protective roles. The terpene scaffold is a common structural feature found in many natural and synthetic compounds that have pharmaceutical, cosmetic, and agrochemical applications, often playing a crucial role in their biological activity. In pharmacology, these molecules are substantial contributors to six major classes of drugs: steroids, tocopherols, taxanes, artemisinins, ingenanes, and cannabinoids (Jansen and Shenvi, 2014;Yang et al., 2020).

Biosynthesis of terpenes proceeds through the formation of C5 isoprenyl diphosphate isomers, isopentenyl diphosphate (IPP) and dimethylallyl diphosphate (DMAPP), which undergo condensation to yield various groups of terpenes. This reaction, catalyzed by isoprenyl diphosphate synthases (IDS), typically starts with the head-to-tail (1’-4) coupling of IPP and DMAPP, resulting in the linear monoterpene scaffolds with *trans*- or *cis*-configuration. The chain can be further elongated through either additional head-to-tail or head-to-head (1’-1) linkages. The initial linear skeletons can be cyclized and further modified by terpene synthases and other enzymes, e.g. cytochromes P450 (Boutanaev et al., 2015). Most terpenes are formed using this mechanism and are classified as regular terpenes. In contrast, a limited number of so called irregular terpenes are produced by the head-to-middle linkage of two DMAPP molecules, which results in the direct formation of branched or cyclized structures with specific biological activities (Fig. 1).

**Fig. 1.**
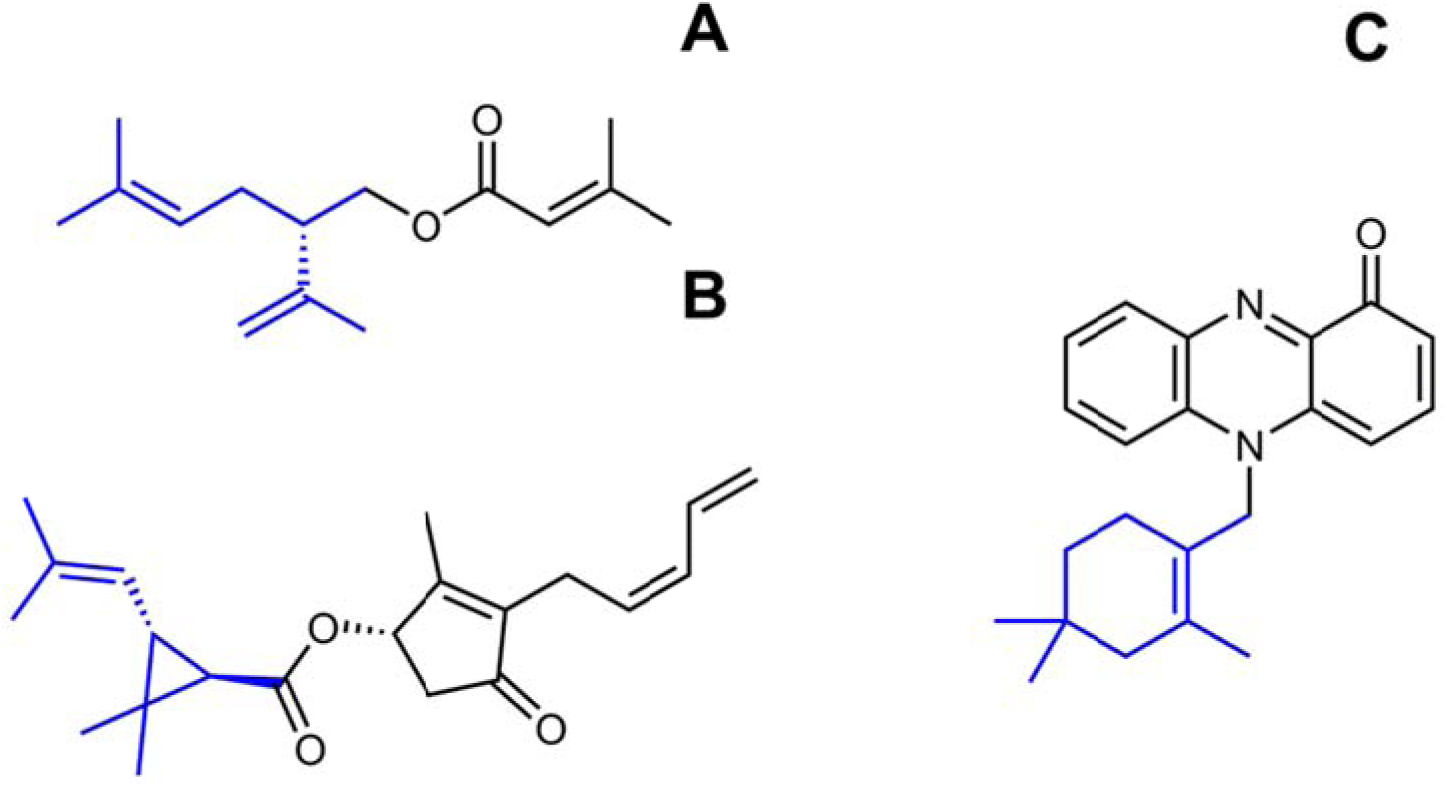
Examples of irregular monoterpene motifs in the structures of biologically active small molecules. A, (S)-(+)-Lavandulyl senecioate, the sex pheromone of vine mealybug, *Planococcus ficus*. B, Pyrethrin I, a natural irregular monoterpene known for its potent insecticidal activity. C, Lavanducyanin, a phenazine-derived meroterpenoid exhibiting cytotoxic and antibiotic properties.

For example, compounds with branched lavandulyl or various cyclic structures often function as insect pheromones and are promising candidates for developing sustainable pest control strategies (Zou and Millar, 2015;Rizvi et al., 2021). Natural pyrethrins, derivatives of the irregular monoterpene chrysanthemyl diphosphate with a cyclopropane skeleton, along with chemically produced pyrethroids serve as highly effective insecticides. They are typically harmless to animals and are known for their rapid biodegradation (Lybrand et **a**l., 2020). Cyclolavandulyl moiety is a part of several types of pharmaceutically valuable compounds, e.g. phenazine-derived meroterpenoids, such as lavanducyanin, exhibiting cytotoxic and antibiotic activiti**e**s, and neuroprotective carbazole alk loid lavanduquinocin (Shin-Ya et al., 1995;Baumgartner and McKinnie, 2024).

Although significant progress has been made in the chemi**c**al synthesis of various terpenes (Kanwal et al., 2022), the producti n of irregular mono**t**erpenes often remains inefficient and may involve hazardous reagents. For example, synthesizing achiral β-cyclolavandulol from a ommercial precursor through a 5-step chemical route yielded 33-50% (Kinoshita et al., 1995; Knölker and Fröhner, 1998). In a stereodivergent total synthesis of lavandulol, the process involved 11-16 reaction steps and resulted in an overall yield of 6-26% (Bhosale and Waghmode, 2017). As an environmentally friendly alternative, biocatalytic methods are developed, e.g. preparing enantiomerically pure (*R*)-lavandulol by the enzyme-catalyzed resolution of racemic mixtures. (Zada and Dunkelblum, 2006).

Multiple attempts have been made to produce regular terpenes using microbial biofactories. Extensive metabolic engineering aimed at increasing the precursor supply and inhibiting competing pathways combined with the optimization of fermentation conditions resulted in impressive sesquiterpene titers reaching up to 140 g/l farnesene (Meadows et al., 2016) and more than 40 g/l amorphadiene in yeast (Westfall et al., 2012). For monoterpenes, the accumulation levels are significantly lower (Yang et al., 2025) due to their volatility and toxicity to the host cells. An effective way of alleviating these problems is in situ product extraction (Brennan et al., 2012) or conversion into less volatile and toxic molecules. For geraniol, esterification to geranyl acetate increased the product accumulation in *E. coli* systems to 10.4 and 19 g/l (Wang et al., 2022b;Shukal et al., 2024), while the highest reported level of free alcohol for this chassis is 2.1 g/l (Wang et al., 2021). Further development of this strategy allowed for recovery of geraniol at concentration of 13.2 g/l by hydrolysis of geranyl acetate (Wang et al., 2022a).

The microbial production of irregular monoterpenes has been rarely reported. Recently, engineered *Saccharomyces cerevisiae* achieved a yield of 309 mg/l of lavandulol through fed-batch fermentation in a 5 liter bioreactor (Nie et al., 2025).

More research on biotechnological production of irregular monoterpenes has been carried out using plants as hosts. Plant cells offer access to precursor flux through either the mevalonic acid (MVA) pathway in the cytoplasm or the methylerythritol phosphate (MEP) pathway in the plastids, with the ability to localize enzymes in different cell compartments. Efforts to engineer terpenoid metabolism in plants to enhance the accumulation of native irregular monoterpenes or to produce heterologous compounds have involved stable genetic transformation of tobacco (Yang et al., 2014;Matáeos-Fernández et al., 2023), as well as host species that naturally produce high levels of terpenoids, such as *Chrysanthemum morifolium* (Hu et al., 2018), *Lavandula×intermedia* (Munoz-Bertomeu et al., 2006) and *Solanum lycopersicum* (Xu et al., 2018a).

A convenient platform for the rapid production of recombinant proteins and manipulating plant metabolism is transient expression (Gerasymenko et al., 2019;Nosaki et al., 2021). Advantages of this strategy include a high level of recombinant protein accumulation, which can exceed that of nuclear transformed plants by several times. Additionally, it offers flexibility in testing various gene combinations and allows for the restriction of expression products to certain plant organs. This capability enables temporary accumulation of potentially toxic products at concentrations that would be harmful to the entire plant.

There is particular interest in using plants for the transient biosynthesis of small molecules with valuable activities, employing combinations of heterologous enzymes directed to specific compartments within the plant cell. The development of such a platform offers a source of inexpensive material for *in planta* or whole-cell biocatalysis as transient expression protocols are easily scalable from analytical to preparative and manufacturing levels. However, it requires adaptation of foreign proteins to the plant’s intracellular environment and optimization of intrinsic metabolic fluxes.

The transient expression of genes encoding irregular IDS enzymes, lavandulyl and chrysanthemyl diphosphate synthases (LDS and CDS), was reported in *N. benthamiana* (Xu et al., 2018b;Matáeos-Fernández et al., 2023). This approach was used for proving the functionality of IDS enzymes and enzymes that oxidizes *trans*-chrysanthemol to chrysanthemic acid, but no experiments on increasing the yield of heterologous irregular monoterpenes by expanding the supply of precursors were attempted.

In this study, we endeavoured to improve the performance of plants as production system for irregular monoterpenes by enlarging the pool of DMAPP, the substrate for IDS performing the irregular coupling. To enhance the metabolic flux through the MEP and MVA pathways, the corresponding bottleneck enzymes were overexpressed. In addition, the enzymes producing DMAPP from dimethylallyl phosphate and IPP were introduced (Fig. 2). Previous research has shown that the major part of heterologous terpenoids in transgenic plants underwent glucosylation (Lucker et al., 2001;Hu et al., 2018;Xu et al., 2018a;Xu et al., 2018b), a common modification allowing for sequestration and storage of hydrophobic and often toxic monoterpenes in plant cells (Dudareva and Pichersky, 2008). Therefore, we focused on the accumulation of non-volatile forms of irregular monoterpenes that can be easily extracted from the harvested plant material unlike volatile metabolites requiring continuous trapping techniques. We identified six major components of methanolic extracts from *N. benthamiana* plants expressing irregular IDS enzymes, five of them being previously unknown substances (Fig. 2, compounds **1**-**6**), and achieved the yield of the products in the range of 1.4 – 3.9 μmol g^-1^ FW r presenting the highest reported concentrations of **i**rregular monoterpenes produced in a plant syst m.

**Fig. 2.**
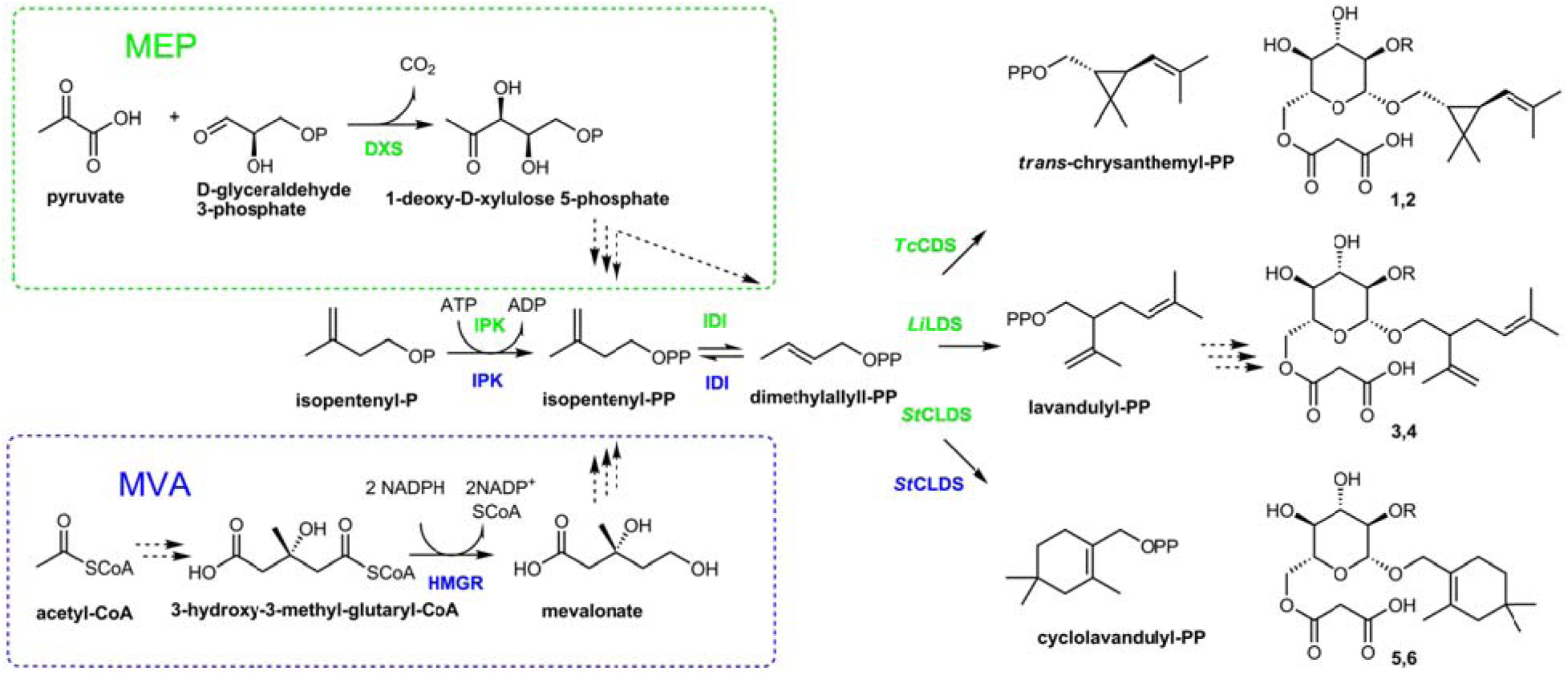
Selected steps of the metabolic pathway for irregular monoterpene malonyl glucoside biosynthesis in *N. benthamiana*. Solid arrows indicate the reactions catalized by the enzymes heterologously expressed in this study. 1-deoxy-D-xylulose 5-phosphate synthase (DXS), chrysanthemyl diphosphate synthase (*Tc*CDS), and lavandulyl diphosphate synthase (*Li*LDS) were localized to chloroplasts (highlighted in green); 3-hydroxy-3-methyl-glutaryl-CoA reductase (HMGR) operated in cytoplasm (highlighted in blue); cyclolava**n**dulyl diphosphate synthase (CLDS), isopentenyl phosphate kinase (IPK), and isopentenyl diphosphate somerases (IDI) were expressed with and without chloroplast targeting peptide. Structures **1**, **3**, **5**, R = H; **2**, **4**, **6**, R = Glu. Abbreviation: Glu, glucopyranose; CoA, coenzyme A; MEP, methylerythritol phosphate pathway; MVA, mevalonate pathway; PP, diphosphate; P, phosphate.

## 2. Materials and methods

### 2.1. Analytical instruments and assays

LC-ESI-MS analysis was performed on a 1260 Infinity HPLC system coupled to a G6120B quadrupole mass spectrometry detector (Agilent, USA). The separation was carried out on a Zorbax Extend-C18 (4.6 x 150 mm, 3.5 μm) column (Agilent, USA), according to the protocol described by Nagel (Nagel et al., 2012). The detection of C10 prenyl glycosides was performed in a negative total ion current and a single ion mode.

An Impact II system (Bruker Daltonik, Germany) was used for recording ESI-HR-MS spectra.

HPLC analysis was performed using a 1260 Infinity system connected to a Zorbax Eclipse Plus C18 column, 4.6 × 250 mm, 5 μm (Agilent, USA), and the mobile phase consisted of 50 mM ammonium acetate (pH 4.5) (A) and 70:30 (v/v) acetonitrile:50 mM ammonium acetate (pH 4.5) (B) (Sun et al., 2010) with the binary gradient elution program (% B): 20–50 within 25 min; 50-100 within 1 min. The column was flushed with 100% B for 3 min and re-equilibrated with 20% B for an additional 3 min. The flow rate was set to 1.0 ml/min and detection was at 256 nm.

^1^H and ^13^C NMR, CLIP-COSY (25 Hz mixing (Koos et al., 2016), NOESY (500 ms mixing), ^1^H -^13^C multiplicity-edited HSQC and ^1^H -^13^C HMBC, TOCSY and DOSY (stimulated echo, bipolar gradient pairs, δ/2 = 1 ms, Δ = 60 ms, 16 gradient steps linearly spaced between 2% and 98%) spectra were recorded in DMSO-*d_6_* using an AVANCE III HD instrument (Bruker, USA) equipped with a 5 mm QCI CryoProbe (^1^H/^19^F- ^31^P/^13^C/^15^N/^2^H with a z-gradient coil with maximum amplitude of 53 G/cm) at 700 MHz (^1^H) and 176 MHz (^13^C). All measurements were carried out at 300 K. Bruker TopSpin 3.5pl7 was used for data acquisition using pulse sequences as provided by the vendor library. Default parameter values were used unless noted otherwise.

### 2.2. Genetic vectors and transient expression

Genes of IDS and auxiliary enzymes used in this study (Table 1) were either synthesized or amplified as described in the supplementary materials (S1.2). Gene *GFP* served as a physiologically neutral control. The cloned or synthetic genes were inserted into genetic vectors using the Golden Braid modular cloning system (Sarrion-'Perdigones et al., 2013).

**Table 1.**
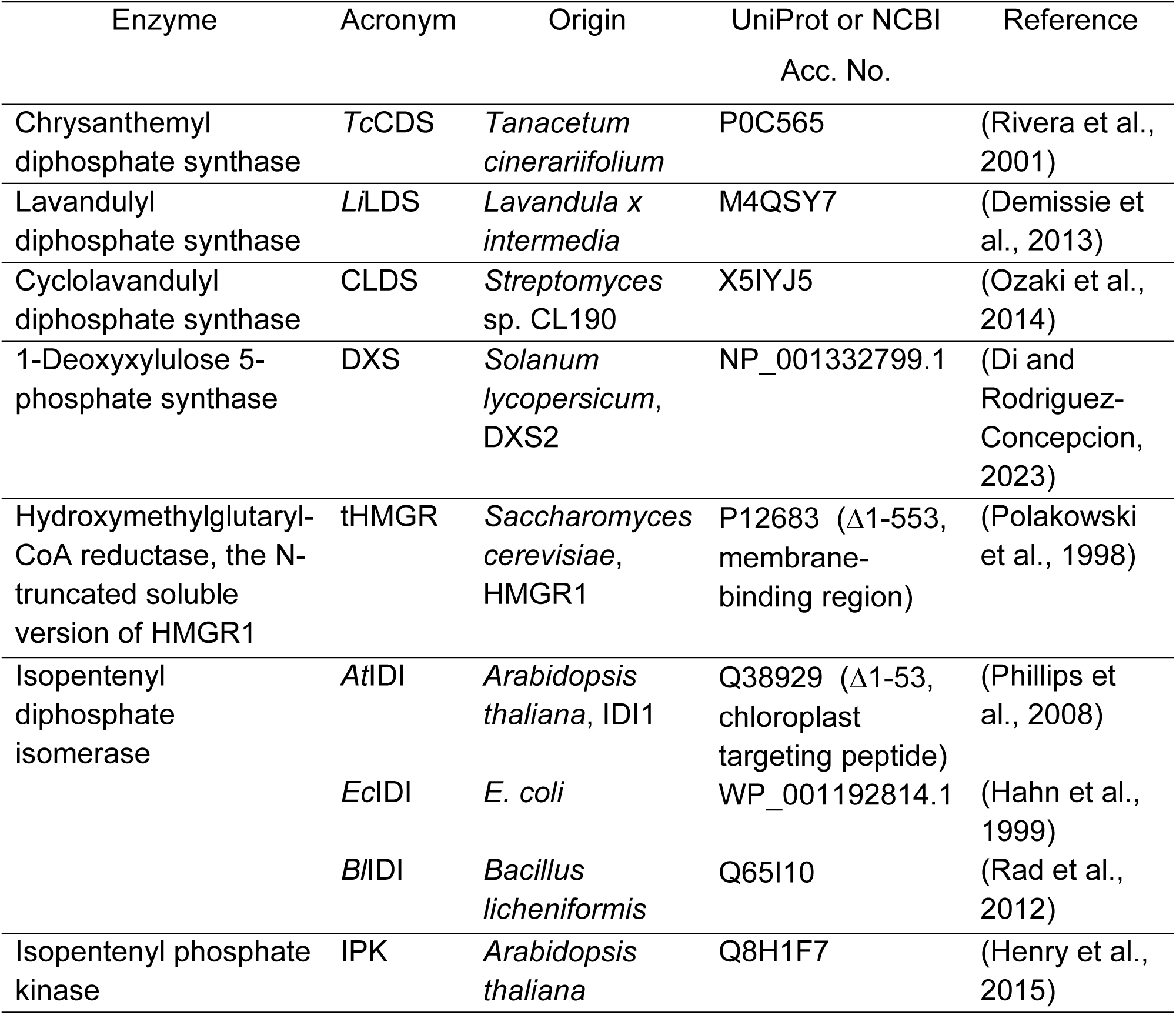
Enzymes used in experiments on the biosynthesis of irregular monoterpene malonyl glucosides in *N. benthamiana*.

The transcriptional units (TUs) for all genes were assembled in alpha-level vectors and included 35S CaMV promoter and nopaline synthase gene terminator from *A. tumefaciens*. For protein transport into chloroplasts, the sequence encoding an artificial chloroplast transit peptide from the pICH13688 plasmid (ICONGenetics, Halle, Germany, GenBank accession number AM888351.1) was added to the TUs for IDIs and IPK. The TUs for IDS genes were combined in omega-level vectors with the TU for p19 suppressor of post-transcriptional gene silencing from the tomato bushy stunt virus. The TUs for auxiliary genes were either expressed from alpha-level vectors or combined in omega-level vectors as shown in Table S1.1. The vectors were introduced into the *A. tumefaciens* EHA105 strain. The transient expression was performed in plants of *N. benthamiana* as described earlier (Gerasymenko et al., 2019). For co-expression, the bacterial suspensions carrying corresponding vectors were mixed in equal volumes before infiltration (Table S1.1). The biomass was collected six days after infiltration.

### 2.3. Quantification of monoterpene malonyl glucosides

The leaf explants (130 ± 10 mg) were frozen in liquid nitrogen, ground and extracted with 200 μl of 80% methanol in an ultrasound bath filled with water/ice slurry during 30 min. After centrifugation (10 min, 17 000 x g), the supernatants were transferred in new Eppendorf tubes and centrifuged once more for 10 min at 17 000 g. The supernatants after the second centrifugations were used for LC-ESI-MS as described in 2.1.

A one-way ANOVA and Tukey’s HSD test were performed using the calculator retrieved from https://www.socscistatistics.com/tests/anova/default2.aspx.

### 2.4. Isolation of irregular monoterpene malonyl glucosides

For the preparative isolation of irregular monoterpene malonyl glucosides (**1**-**6**), the plant material from the various experiments on the expression of the corresponding IDS gene was combined. The infiltrated leaves (20-25 g) were frozen in liquid nitrogen, ground and extracted with 80% (v/v) methanol (40 ml) in an ultrasound bath filled with water/ice slurry during 30 min. After centrifugation (10 min, 12 500 x g), the supernatant was filtered through filter paper, dried on a rotary evaporator and taken up in 20 ml methanol. After the second centrifugation (10 min, 12 500 x g), the supernatant was filtered through filter paper, dried on the rotary evaporator and dissolved in 2 ml methanol. HPLC-MS analysis displayed that the extract contained the corresponding monoterpene malonyl mono- and diglucosides in different proportions. For normal phase chromatography, the column (18 mm id) was packed with silica gel (40–60 μm, 230–400 mesh, 10 g) equilibrated in ethyl acetate. After extract application, the column was washed with 50 ml ethyl acetate, and the fractions of 10 ml were collected during elution with 150 ml ethyl acetate:methanol 4:1 mixture followed by 150 ml ethyl acetate:methanol 35:15 mixture and 50 ml methanol. The fractions were analysed by HPLC-MS and those containing monoterpene malonyl mono- and diglucosides were pooled (typically fractions 4-13 eluted with acetate:methanol 4:1 mixture for monoglucosides and fractions 14-28 eluted with acetate:methanol 35:15 mixture for diglucosides). After drying on a rotary evaporator, the combined fractions were dissolved in 1 ml 5 mM ammonium bicarbonate (ABC) and subjected to reverse phase chromatography on a Sep-Pak C18 1 cc Vac Cartridge (Waters, USA) activated with acetonitrile and equilibrated with ABC. The cartridge was washed with 3 ml ABC, and elution was performed with an acetonitrile:ABC 95:5 mixture (10-12 ml) collecting 1 ml fractions. After LC-MS analysis, the fractions containing monoterpene malonyl mono- or diglucosides were combined, dried on a rotary evaporator, dissolved in methanol, and separated on ALUGRAM® Xtra SIL G/UV254 TLC plates (0.2 mm silica gel layer, Macherey-Nagel (Germany)) using ethyl acetate:methanol 1:1 mixture as a mobile phase. After the plates were resolved and dried, a 1.5 cm strip was cut from one side and developed using a potassium permanganate stain. The silica gel from the area corresponding to the position of the main compound in the stained section was extracted with methanol which yielded 2-5 mg of material. The obtained samples were dissolved in DMSO-d_6_ (99.8 atom%D, Roth (Germany)) for NMR analysis.

Chrysanthemyl-1*-O-*(6*-O-*malonyl)*-β-*D-glucopyranoside (**1**): amorphous solid; ESIMS m/z 357.2 [M - COOH] ^−^; HRESIMS m/z 803.3721 [2M - H] ^−^ (calcd for C_38_H_59_O_18_, 803.3707; Δmass 1.7 ppm); m/z 401.1833 [M - H] ^−^ (calcd for C_19_H_29_O_9_, 401.1817; Δmass 4.0 ppm); m/z 357.1937 [M - COOH] ^−^ (calcd for C_18_H_29_O_7_, 357.1919; Δmass 5.0 ppm); ^1^H-NMR (700 MHz, DMSO-d_6_) δ (ppm): 4.86 (d, *J* = 8.2 Hz, 1H), 4.16 – 4.05 (m, 2H), 4.13 (d, *J* = 7.9 Hz, 1H), 3.82 (dd, 1H, *J* = 11, 8.7 Hz), 3.43 (dd, 1H, *J* = 11, 5.7 Hz), 3.31 – 3.27 (m, 1H), 3.14 (t, *J* = 9.4, 8.9 Hz, 1H), 3.09 (t, *J* = 9.3, 8.9 Hz, 1H), 2.96 (t, *J* = 9.4, 7.9 Hz, 1H), 1.65 (s, 3H), 1.62 (s, 3H), 1.10 – 1.07 (m, 1H), 1.07 (s, 3H), 0.99 (s, 3H), 0.75 (dt, *J* = 8.5, 5.6 Hz, 1H); ^13^C-NMR (176 MHz, DMSO-d_6_) δ (ppm): 169.19, 168.43, 131.71, 123.91, 102.27, 76.41, 73.69, 73.36, 70.10, 68.56, 63.51, 45.61, 31.8, 27.56, 25.43, 21.79, 22.37, 21.22, 18.12; (Supplementary material, Table S1.3, Figure S2.1-S2.8).

Chrysanthemyl-1*-O-*(6*-O-*malonyl)*-β-*D-glucopyranosyl-(1→2)*-β-*D-glucopyranoside (**2**): amorphous solid; ESIMS m/z 519.3 [M - COOH] ^−^; HRESIMS m/z 563.2366 [M - H] ^−^ (calcd for C_25_H_39_O_14_, 563.2345; Δmass 3.7 ppm); m/z 519.2452 [M - COOH] ^−^ (calcd for C_18_H_29_O_7_, 519.2447; Δmass 1.0 ppm); ^1^H-NMR (700 MHz, DMSO-d_6_) δ (ppm): 4.86 (d, *J* = 8.2 Hz, 1H), 4.40 (d, *J* = 7.9 Hz, 1H), 4.31 (d, *J* = 7.7 Hz, 1H), 4.14 – 4.10 (m, 1H); 4.07 (d, *J* = 11.6 Hz, 1H), 3.79 (t, *J* = 10.7, 8.1 Hz, 1H), 3.62 (d, *J* = 11.9 Hz, 1H), 3.51 – 3.47 (m, 2H), 3.38 (t, *a* = 9.1 Hz, 1H), 3.36 – 3.33 (m, 1H), 3.30 – 3.26 (m, 1H), 3.16 – 3.13 (m, 1H), 3.10 – 3.16 (m, 2H), 3.06 – 3.03 (m, 1H), 2.99 (t, *J* = 8.3 Hz, 2H), 2.92 – 2.88 (m, 2H), 1.65 (s, 3H), 1.63 (s, 3H), 1.11 – 1.08 (m, 1H), 1.07 (s, 3H), 0.98 (s, 3H), 0.81 – 0.77 (m, 1H); ^13^C-NMR (176 MHz, DMSO-d_6_) δ (ppm): 169.72, 167.19, 131.49, 123.94, 103.9, 101.02, 81.68, 77.02, 76.06, 75.51, 74.92, 73.71, 69.76, 69.72, 68.83, 63.29, 60.72, 45.98, 31.64, 27.62, 25.42, 22.36, 21.53, 21.14, 18.14; (Supplementary material, Table S1.3, Figure S2.9-S2.16). The sample used for NMR analyses contained impurities that hindered the direct identification of carbon resonance positions in the ^13^C NMR spectrum. The corresponding signal values were retrieved from HSQC and HMBC spectra.

Lavandulyl-1*-O-*(6*-O-*malonyl)*-β-*D-glucopyranoside (**3**): amorphous solid; ESIMS m/z 357.2 [M - COOH] ^−^; HRESIMS m/z 803.3716 [2M - H] ^−^ (calcd for C_38_H_59_O_18_, 803.3707; Δmass 1.1 ppm); m/z 401.1837 [M - H] ^−^ (calcd for C_19_H_29_O_9_, 401.1817; Δmass 5.0 ppm); m/z 357.1937 [M - COOH] ^−^ (calcd for C_18_H_29_O_7_, 357.1919; Δmass 5.0 ppm); ^1^H-NMR (700 MHz, DMSO-d_6_) δ (ppm): 5.02 (t, 1H, *J* = 7.0 Hz), 4.73 (s, 1H), 4.69 (s, 1H), 4.14 (d, *J* = 7.8 Hz, 1H), 4.08-4.12 (m, 2H), 3.63 (dd, 1H, *J* = 9.8, 7.1 Hz), 3.44 (dd, 1H, *J* = 9.8, 6.6 Hz), 3.30 (m, 1H), 3.14 (t, *J* = 8.9 Hz, 1H), 3.09 (t, *J* = 10.6, 9.3 Hz, 1H), 2.95 (t, *J* = 9.3, 7.8 Hz, 1H), 2.92 – 2.88 (m, 2H), 2.3 (m, 1H), 2.2 (dt, *J* = 13.5, 6.4 Hz, 1H), 1.98 (m, 1H), 1.64 (s, 3H), 1.63 (s, 3H), 1.56 (s, 3H); ^13^C-NMR (176 MHz, DMSO-d_6_) δ (ppm): 169.7, 167.63, 145.49, 131.36, 122.48, 111.81, 102.92, 76.33, 73.75, 73.4, 70.8, 70, 63.43, 46.45, 45.86, 28.21, 25.63, 20.12, 17.77; (Supplementary material, Table S1.3, Figure S2.17-S2.24).

Lavandulyl-1*-O-*(6*-O-*malonyl)*-β-*D-glucopyranosyl-(1→2)*-β-*D-glucopyranoside (**4**): amorphous solid; ESIMS m/z 519.3 [M - COOH] ^−^; HRESIMS m/z 563.2351 [M - H] ^−^ (calcd for C_25_H_39_O_14_, 563.2345; Δmass 0.9 ppm); m/z 519.2435 [M - COOH] ^−^ (calcd for C_18_H_29_O_7_, 519.2447; Δmass 2.2 ppm); ^1^H-NMR (700 MHz, DMSO-d_6_) δ (ppm): 5.02 (t, 1H, *J* = 7.1 Hz), 4.73 (s, 1H), 4.68 (s, 1H), 4.38 (d, *J* = 7.7 Hz, 1H), 4.31 (d, *J* = 7.6 Hz, 1H), 4.16 – 4.06 (m, 2H), 3.66 (dd, 1H, *J* = 9.4, 7.4 Hz), 3.61 (d, *J* = 11.5 Hz, 1H), 3.50 – 3.48 (m, 1H), 3.44 (dd, 1H, *J* = 9.4, 6.2 Hz), 3.39 (t, *J* = 9.0 Hz, 1H), 3.37 – 3.34 (m, 2H), 3.31 – 3.27 (m, 1H), 3.19 – 3.16 (m, 1H), 3.17 – 3.10 (m, 2H), 3.03 – 3.00 (m, 1H), 2.98 (t, *J* = 8.4 Hz, 1H), 2.95 – 2.90 (m, 2H), 2.29 (m, 1H), 2.2 (dt, *J* = 13.7, 6.4 Hz, 1H), 1.99 (m, 1H), 1.64 (s, 3H), 1.63 (s, 3H), 1.56 (s, 3H); ^13^C-NMR (176 MHz, DMSO-d_6_) δ (ppm): 169.6, 168.15, 145.58, 131.22, 122.57, 111.68, 104.11, 101.36, 81.4, 77.1, 76.01, 75.58, 75.1, 73.53, 70.74, 69.87, 69.77, 63.33, 60.75, 46.54, 45.69, 28.14, 25.62, 20.19, 17.78; (Supplementary material, Table S1.3, Figure S2.25-S2.32).

β-Cyclolavandulyl-1*-O-*(6*-O-*malonyl)*-β-*D-glucopyranoside (**5**): amorphous solid; ESIMS m/z 357.2 [M - COOH] ^−^; HRESIMS m/z 803.3701 [2M - H] ^−^ (calcd for C_38_H_59_O_18_, 803.3707; Δmass 0.8 ppm); m/z 401.1814 [M - H] ^−^ (calcd for C_19_H_29_O_9_, 401.1817; Δmass 0.9 ppm); m/z 357.1917 [M - COOH] ^−^ (calcd for C_18_H_29_O_7_, 357.1919; Δmass 0.5 ppm); ^1^H-NMR (700 MHz, DMSO-d_6_) δ (ppm): 4.23 (d, *J* = 11.4 Hz, 1H), 4.18 (dd, *J* = 11.8, 1.8 Hz, 1H), 4.06 (dd, *J* = 11.8, 6.9 Hz, 1H), 4.03 (d, *J* = 7.9 Hz, 1H), 3.97 (d, *J* = 11.4 Hz, 1H), 3.24 (ddd, *J* = 8.9, 6.9, 1.8 Hz, 1H), 3.11 (t, *J* = 9.4, 8.8 Hz, 1H), 3.07 (t, *J* = 9.4, 8.8 Hz, 1H), 2.97 (t, *J* = 9.1, 8.0 Hz, 1H), 2.96 – 2.94 (m, 2H), 2.14 (d, *J* = 17.4 Hz, 1H); 1.95 (d, *J* = 17.4 Hz, 1H), 1.73 (s, 2H), 1.62 (s, 3H), 1.28 (t, *J* = 6.6 Hz, 2H), 0.86 (s, 3H), 0.86 (s, 3H); ^13^C-NMR (176 MHz, DMSO-d_6_) δ (ppm): 169.31, 168.08, 130.76, 124.58, 100.43, 76.47, 73.89, 73.22, 70.2, 66.73, 63.58, 45.58, 45.32, 35.12, 28.77, 28.12, 28.08, 24.95, 18.82; (Supplementary material, Table S1.3, Figure S2.33-S2.40).

β-Cyclolavandulyl-1*-O-*(6*-O-*malonyl)*-β-*D-glucopyranosyl-(1→2)*-β-*D-glucopyranoside (**6**): amorphous solid; ESIMS m/z 519.3 [M - COOH] ^−^; HRESIMS m/z 563.2342 [M - H] ^−^ (calcd for C_25_H_39_O_14_, 563.2345; Δmass 0.7 ppm); m/z 519.2463 [M - COOH] ^−^ (calcd for C_18_H_29_O_7_, 519.2447; Δmass 3.1 ppm); ^1^H-NMR (700 MHz, DMSO-d_6_) δ (ppm): 4.39 (d, *J* = 7.8 Hz, 1H), 4.24 (d, *J* = 7.7 Hz, 1H), 4.15 (d, *J* = 11.6 Hz, 0H), 4.13 – 4.08 (m, 2H), 4.09 (dd, *J* = 11.6, 6.5 Hz, 1H), 3.60 (d, *J* = 11.6 Hz, 1H), 3.51 – 3.47 (m, 1H), 3.41 – 3.35 (m, 1H), 3.34 – 3.31 (m, 1H), 3.31 – 3.28 (m, 1H), 3.18 – 3.14 (m, 1H), 3.17 – 3.11 (m, 1H), 3.15 – 3.11 (m, 1H), 3.05 – 3.02 (m, 1H), 2.98 (t, *J* = 8.3 Hz, 1H), 2.94 – 2.90 (m, 2H), 2.14 (d, *J* = 17.2 Hz, 1H), 2.01 (d, *J* = 17.2 Hz, 1H), 1.71 (s, 2H), 1.60 (s, 3H), 1.27 (t, *J* = 6.5 Hz, 2H), 0.85 (s, 6H); ^13^C-NMR (176 MHz, DMSO-d_6_) δ (ppm): 169.54, 168.13, 129.85, 124.99, 104.01, 99.69, 81.51, 77.01, 76.01, 75.76, 75.07, 73.63, 69.99, 69.72, 67.35, 63.3, 60.8, 45.63, 45.58, 35.16, 28.74, 28.25, 28.02, 24.68, 18.87; (Supplementary material, Table S1.3, Figure S2.41-S2.48).

### 2.5. Acid hydrolysis and derivatization with 3-methyl-1-phenyl-5-pyrazolone (MPP)

Compounds **1**-**6** (2-5 mg of each) were incubated with 2 ml of 2 M trifluoroacetic acid in locked Eppendorf tubes at 105°C for six hours. After incubation, the mixtures were cooled and centrifuged. The supernatants were then diluted with 10 ml of methanol and evaporated under vacuum. The resulting residues were dissolved in 100-200 μl of deionized water, and 5-10 μl of 3 M NaOH was added to neutralize the residual acid. Reference samples were prepared as 50 mM solutions of monosaccharides.

Derivatization was performed according to the method described by Sun et al. (Sun et al., 2010) with minor modifications. For derivatization, 20 μl of the hydrolyzed glycoside solution or reference sample was mixed with 180 μl of acetonitrile, along with 20 μl of a 0.5 M MPP solution in methanol and 10 μl of a 25% ammonium hydroxide solution. The samples were incubated for 25 minutes at 70°C, then cooled and diluted with 760 μl of acetonitrile before HPLC analysis.

The relative configuration of the sugar moiety in compounds **1**-**6** was confirmed by comparing the retention times of the derivatized carbohydrates to that of a reference sample of D-glucose (t_R_ = 19.6 min).

## 3. Results

### 3.1. Transient expression of heterologous IDS enzymes in *N. benthamiana*

We applied *trans*- and *cis*-IDS enzymes of plant origin performing irregular coupling, chrysanthemyl diphosphate synthase from *Tanacetum cinerariifolium* (*Tc*CDS) and lavandulyl diphosphate synthase from *Lavandula x intermedia* (*Li*LDS), catalysing the formation of a cyclopropane ring (Rivera et al., 2001) and a branched moiety (Demissie et al., 2013), respectively. Additionally, we included a prokaryotic cyclolavandulyl diphosphate synthase (CLDS), a *cis*-IDS derived from *Streptomyces* sp. capable of head-to-trunk 1’-2 condensation which is followed by the creation of a six-carbon ring through 4-3’ bond closure (Ozaki et al., 2014). The *Tc*CDS and *Li*LDS enzymes were expressed with their native chloroplast transit peptides. The bacterial CLDS was examined in its native form lacking identifiable targeting signals and in fusion with an artificial chloroplast targeting peptide.

To monitor terpene glucoside accumulation, we extracted the plant material with 80% methanol on the 6^th^ day post infiltration and analyzed it using LC-MS. Transient expression of each irregular IDS enzyme in *N. benthamiana* resulted in the detection of two major new compounds in the leaf methanolic extracts (Fig. 3). These metabolites were identified as 6*-O-*malonyl*-β-*D-glucopyranoside and 6*-O-*malonyl*-β-*D-glucopyranosyl-(1→2)-*β*-D-glucopyranoside derivatives of chrysanthemol, lavandulol and cyclolavandulol (compounds **1-6**, Fig. 2). Among these substances, only chrysanthemyl-1*-O-*(6*-O-*malonyl)*-β-*D-glucopyranoside (**1**) was characterized previously (Yang et al., 2014;Hu et al., 2018). In this report, we provide a detailed elucidation of the structures of new molecules based on one- and two-dimensional NMR assays. The original NMR spectra, along with the CLIP-COSY, HMBC, and NOESY correlations are available in the supplementary materials (Table S1.3; Figure S2.1 - S2.48).

**Fig. 3.**
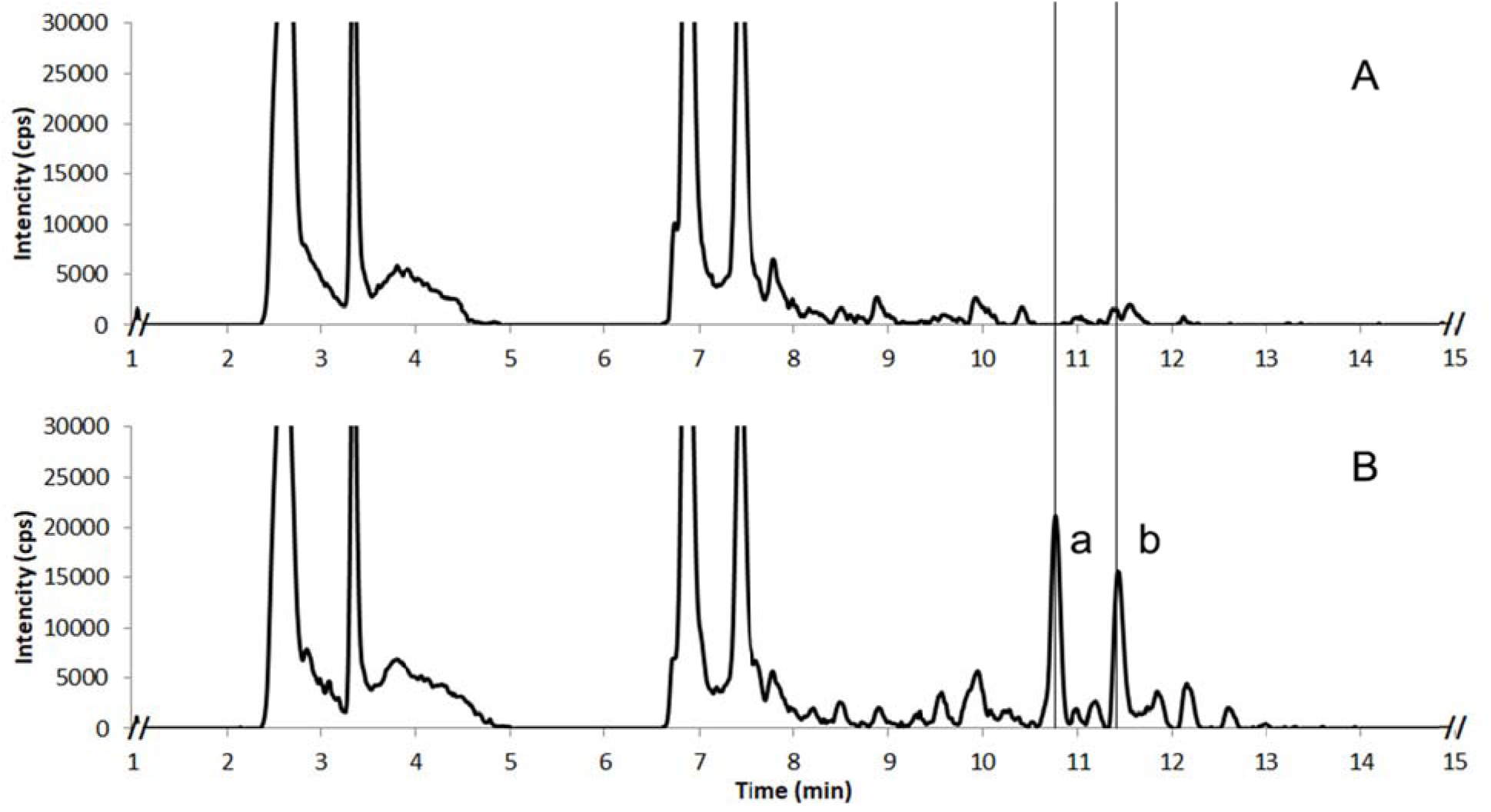
LCMS chromatograms of methanolic extracts from *N. benthamiana* plants expressing GFP as a negative control (A) and an irregular IDS enzyme, chloroplast-targeted CLDS (B). Detection in negative ion mode; two new peaks were identified as 6*- O-*malonyl*-β-*D-glucopyranoside (compound **5,** b) and 6*-O-*malonyl*-β-*D-glucopyranosyl-(1→2)-*β*-D-glucopyranoside (compound **6,** a) derivatives of cyclo**l**avandulyl.

### 3.2. Structure elucidation of irregular monoterpene malonyl glycosides

#### 3.2.1. Monoglucosylated derivatives (6*-O-*malonyl*-β-*D-glucopyranosides) of chrysanthemyl, lavandulyl and cyclolavandulyl (1, 3, and 5)

ESI-HRMS analysis of **1**, **3**, and **5** in negative mode revealed base peak with *m/z* values of 803.3721, *m/z* 803.3716, and *m/z* 803.3701, respec**t**ively. This suggests a molecular formula of C_38_H_59_O_18_ for each compound. Additionally, the spectra displayed ions with *m/z* values of 401.1833, *m/z* 401.1837, nd *m/z* 401.1814, respectively, corresponding to the molecular fo**r**mula C_19_H_29_O_9_ ([M–H]^−^) and indicating that the base peak is likely an adduct [2M–H]^−^.TOCSY spectra of **1**, **3**, and **5** revealed two distinct networks of protons coupled within a spin system. One group of signals was specific to each compound and corresponded to the protons associated with the respective monoterpene moiety. The other group consisted of six protons that exhibited similar NMR shift values across all three molecules, characteristic of a pyranose ring, particularly glucopyranose (Brito-Arias, 2007). The anomeric proton H-1’ appeared as a doublet at δ 4.1 – 4.0 ppm with a coupling constant (*J*) of 7.9-7.8 Hz, indicating an axial-axial interaction with H-2’ and β-glycosidic linkage. The chemical shifts of the C-1’ carbon in compounds **1**, **3** and **5** fell within the range of δ 102.9 – 100.4 ppm, which is characteristic of a glycosidic bond. Three triplets at δ 3.2 – 3.0 ppm, with a coupling of 9-10 Hz, were assigned to the protons H-2’ through H-4’ of D-pyranoses in a ^4^C_1_ conformation. A multiplet (ddd) at δ 3.3 – 3.2 was assigned to H-5’, which was coupled to two H-6’ protons.

The signals for H-6’ in all three compounds were shifted downfield (δ 4.2 – 4.1 ppm), along with the corresponding carbon resonances detected at δ 63.6 – 63.4 ppm. These shifts suggest derivatization of the hydroxyl group at C-6’, which was confirmed by ^1^H -^13^C HMBC analysis. In the HMBC spectra of compounds **1**, **3** and **5**, we observed correlations between the H-6’ protons and a quaternary carbon at δ 169.7 – 169.2. Additionally, two protons in the range of δ 3.0 – 2.9 ppm also interacted with this carbon, as well as with another quaternary carbon having a chemical shift in the range of δ 168.4 – 167.6 ppm. These correlations suggest the presence of two carboxylic groups linked by a methylene carbon, indicating malonylation of the glucopyranose ring at the C-6 position.

In substances **1**, **3**, and **5**, the sugar component was identified as glucose through HPLC analysis following acid hydrolysis and MPP derivatization. The retention time of the derivatized hydrolysis product matched that of the reference sample of D-glucose (Figure S2.49) in the HPLC system designed for separating MPP-derivatized reductive monosaccharides (Sun et al., 2010). While none of the techniques used can definitively determine the absolute configuration of the glucose enantiomer in the terpene derivatives, the absence of L-glucose in higher living organisms leads us to conclude that the compounds investigated are D-glucosides.

The proton and carbon chemical shifts of the monoterpene moiety in substances **1**, **3**, and **5** were consistent with the literature data for the corresponding irregular monoterpene alcohols (Kim et al., 2011;Ozaki et al., 2014;Bergmann et al., 2019), respectively). The exception was the resonance positions at the glycosidic bond, which were shifted downfield. The assignments of these signals were confirmed using 2D NMR experiments. The HMBC spectra for all substances displayed correlations between C-1 of the monoterpene part and the H-1’ proton of the sugar. This data revealed that substances **1**, **3**, and **5** contain chrysanthemyl, lavandulyl, and cyclolavandulyl moieties, respectively, linked to a 6*-O-*malonyl-glucopyranose via a 1*-O-β-*D-glycosidic bond.

Based on the obtained data, we identified the structures of substances **1, 3,** and **5** as follows: chrysanthemyl-1*-O-*(6*-O-*malonyl)*-β-*D-glucopyranoside, lavandulyl-1*-O-*(6*-O-*malonyl)*-β-*D-glucopyranoside, and β-cyclolavandulyl-1*-O-*(6*-O-*malonyl)*-β-*D-glucopyranoside, respectively.

To determine the relative stereochemistry of the chrysanthemyl cyclopropane ring in compounds **1** and **2**, we considered the biosynthesis of chrysanthemol by *Tc*CDS from *T. cinerariifolium,* which produces the *trans*-(1*R*,3*R*)-isomer of the molecule (Yang et al., 2014;Hu et al., 2018;Xu et al., 2018a). The NOESY experiment (Figure S2.8 and S2.16) showed similar intensity signals between H-4 and H-2, H-3, 3H-7, and 3H-9. Analyzing this data with 3D models of energy-optimized conformers of both *cis*- and *trans*-**1** supports the assignment of a *trans* configuration for the molecule. In this configuration, the distances between all interacting protons are within 3 Å, with the distance between protons H-4 and H-2 measuring 2.4 Å. In contrast, the calculated distance between H-4 and H-2 protons in the *cis* configuration increases to 3.6 Å, which would significantly weaken the correlation. The high conformational flexibility of the alkyl chain in compound **3** and **4** prevents a similar NOE-based assignment of the relative configuration of the lavandulyl derivatives.

Regarding the native stereoselectivity of the enzyme *Tc*CDS, data of the NOESY experiment and a single set of signals in the NMR spectra, we can assume a *trans*-(1*R*,3*R*)-configuration for the isolated chrysanthemol glucosides **1** and **2**.

#### 3.2.2. Diglucosylated derivatives (6*-O-*malonyl*-β-*D-glucopyranosyl-(1→2)-*β*-D-glucopyranosides) of chrysanthemyl, lavandulyl and cyclolavandulyl (**2**, **4**, and **6**)

Compared to compounds **1**, **3**, and **5**, the ESI-HRMS data for substances **2**, **4**, and **6** revealed an increase in the [M - H]⁻ peak by 162 mass units with the molecular formula C_25_H_39_O_14_, indicating the addition of a hexose moiety.

NMR analysis of compounds **2**, **4**, and **6** showed the presence of the same monoterpene residues as those in compounds **1**, **3**, and **5**, which are bound to a 6*-O-* malonyl hexopyranose. The data from TOCSY, CLIP-COSY, and HMBC confirmed the presence of an additional isolated group of spin-coupled protons associated with a second hexopyranose ring. The coupling constants of 7.9-7.7 Hz for the anomeric protons in both hexoses suggest that β-glycoside linkages are present.

In compounds **2**, **4**, and **6**, the secondary glycosylation occurred at the 2-O’ position, resulting in the deshielding of the C-2’ by approximately 8 ppm compared to compounds **1**, **3**, and **5**. In all three diglycosides, the C-2’ signal appeared between δ 81.7 – 81.4 ppm, while the anomeric C-1’’ signal, linked at O-2’, was also shifted downfield and observed at δ 104.1 – 103.9 ppm. This behavior is consistent with observations for di- and triterpene*-β-*D-glucopyranosyl-(1→2)*-β-*D-glycosides (Mizutani et al., 1984). No significant shifts in the signals for C-3 and C-4 atoms in any of the sugar moieties were observed. Additionally, the absence of a deshielding effect from the malonyl group resulted in an upfield shift of the C-6’’ resonances (δ 60.8 – 60.7) and a downfield shift of C-5’’ (δ 77.1 – 77.0).

The hydrolysis of compounds **2**, **4**, **6**, followed by MPP derivatization, confirmed that both hexapyranose residues are glucose (Figure S2.49). Consequently, we assigned the following structures: compound **2** is chrysanthemyl-1*-O-*(6*-O-*malonyl)*-β-* D-glucopyranosyl-(1→2)*-β-*D-glucopyranoside; compound **4** is lavandulyl-1*-O-*(6*-O-* malonyl)*-β-*D-glucopyranosyl-(1→2)*-β-*D-glucopyranoside; and compound **6** is β-cyclolavandulyl-1*-O-*(6*-O-*malonyl)*-β-*D-glucopyranosyl-(1→2)*-β-*D-glucopyranoside. To the best of our knowledge, these compounds have not been previously reported and represent a new subgroup of natural irregular monoterpene glycosides.

### 3.3. Accumulation of non-volatile forms of irregular monoterpenes after transient expression in *N. benthamiana*

For all three irregular monoterpene skeletons, the ratios of malonyl mono- to diglucosides varied considerably among samples. To facilitate a clear comparison across experiments, the product yields are represented as a sum of the molar amounts of both glucosylated derivatives for each irregular monoterpene. The expression of *Tc*CDS and *Li*LDS with the native chloroplast targeting peptides resulted in comparable amounts of irregular terpene glucosides **1**-**4**. Chrysanthemyl glucosides accumulated at the concentration of 0.9 ± 0.3 μmol g^-1^ FW (288 ± 102 μg g^-1^ FW of **1** and 77 ± 44 μg g^-1^ FW of **2**); the content of lavandulyl glucosides was determined a**s** 0.6 ± 0.3 μmol g^-1^ FW (161 ± 74 μg g^-1^ FW of **3** and 122 ± 100 μg g^-1^ FW of **4**). Cyt**o**plasmic CLDS formed higher amounts of irregular structures, 1.1 ± 0.3 μmol g^-1^ FW (299 ± 89 μg g^-1^ FW of **5** and 195 ± 98 μg g^-1^ FW of **6**). When CLDS was translocated into chloroplasts, the yield of these products significantly incr**e**ased (p < 0.01), reaching 1.9 ± 0.6 μmol g^-1^ FW (527 ± 176 μg g^-1^ FW of **5** and 330 ± 111 μg g^-1^ FW of **6)** (Fig. 4).

**Fig. 4.**
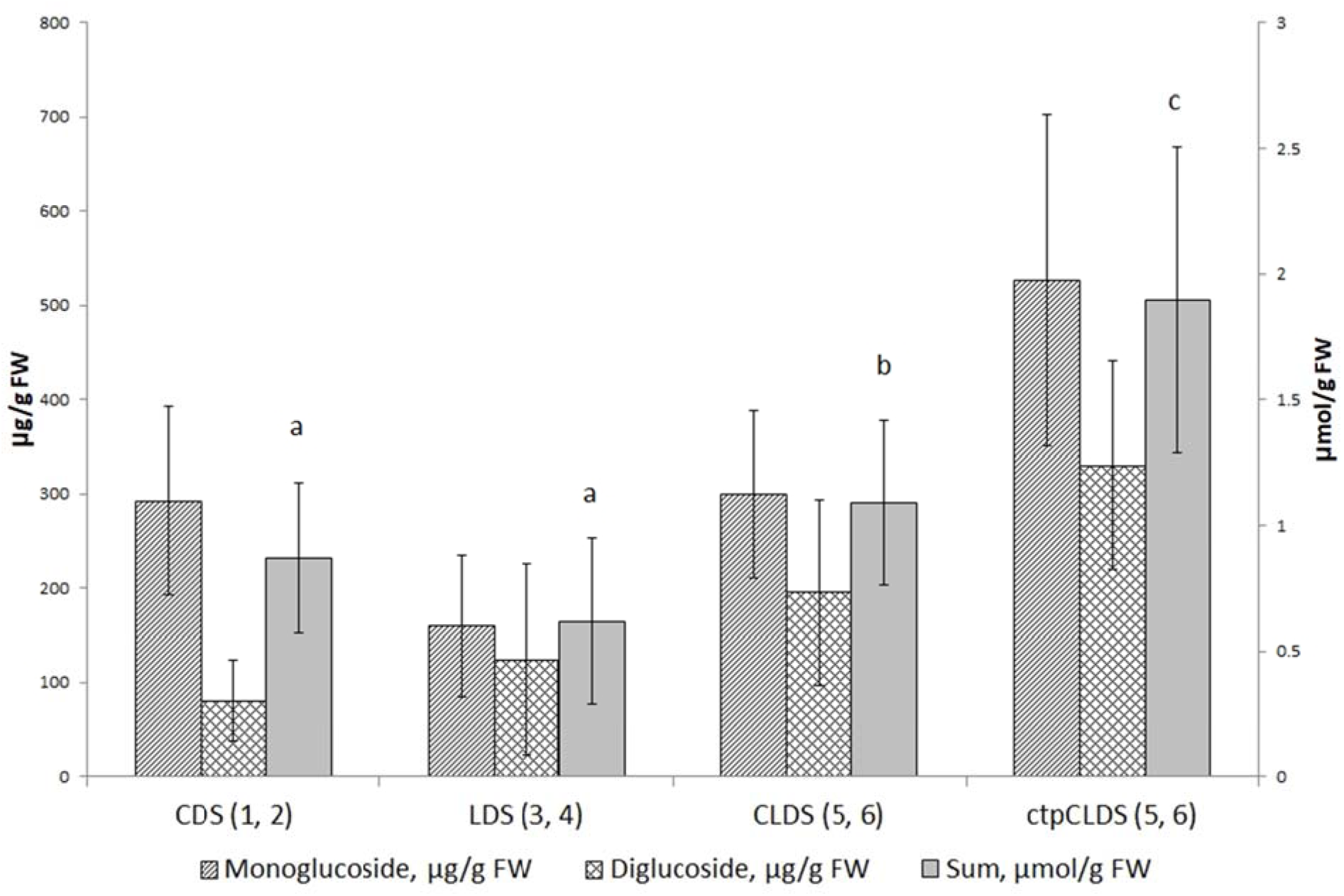
The concentration of monoterpene malonyl-glucosides **1-6** in leaf biomass after transient expression of heterologous IDS in *N. benthamiana*. Different letters indicate significant differences at p < 0.01; *n* = 20-30 repeats.

### 3.4. Supporting irregular monoterpene production in chloroplasts

The common approach to enhancing terpene accumulation in plants involves the overexpression of the enzymes that act as bottlenecks in the biosynthesis pathways for C5 precursors. For the MEP pathway in plastids, metabolic control analysis indicated that the major controlling enzyme is 1-deoxyxylulose 5-phosphate synthase (DXS) (Wright et al., 2014). DXS was repeatedly shown to be the most effective co-expression partner for increasing the biosynthesis of heterologous regular terpenoids in chloroplasts of transiently transformed *N. benthamiana*, e.g. for formation of mono-(Park et al., 2022), sesqui- (Li et al., 2019), and diterpenes (Forestier et al., 2021). Overexpression of DXS was successfully employed also for enhancing the production of native components of essential oil in transgenic lavender (Munoz-Bertomeu et al., 2006). However, when the DXS enzyme was co-expressed with the chloroplast-localized *Tc*CDS, we did not observe a statistically significant increase in glucoside accumulation (p > 0.05), although the mean value was slightly higher.

The next limiting factor in biosynthesis of irregular terpenes may be the unfavourable ratio of DMAPP to IPP. While the MEP pathway produces both isoprenyl diphosphate isomers, IPP is the predominant product, and the MVA pathway ends in IPP only. The overexpression of isopentenyl diphosphate isomerases (IDI), which interconvert C5 precursors, was beneficial even for the formation of regular terpenes that result from the joining of IPP and DMAPP (Chen et al., 2012). We hypothesized that this reaction may be particularly essential for enhancing the irregular coupling involving two DMAPP units. Indeed, a significant improvement in product formation was achieved by simultaneously alleviating two bottlenecks in DMAPP formation, the DXS reaction and shifting the ratio between IPP and DMAPP by an IDI activity. In our experiments, we utilized three IDIs. Two of these enzymes, *Ec*IDI1 from *E. coli* and *At*IDI1 from *A. thaliana*, belong to the widespread type I of the IDI family. The third enzyme, *Bl*IDI2 from *B. licheniformis*, is classified as type II, which is primarily found in thermophilic bacterial and archaeal species and differs significantly in structure from the type I (Berthelot et al., 2012). For all three applied IDI enzymes, we found that co-expressing a chloroplast-targeted isomerase with DXS and *Tc*CDS resulted in a significant increase (p < 0.01) in chrysanthemyl glucoside accumulation when compared to *Tc*CDS expressed alone or in combination with DXS. Co-expressing *Tc*CDS with IDIs alone did not produce any change in product levels (p > 0.05). Additionally, we observed no effect on product **a**ccumulation when *Tc*CDS was co-expressed with HMGR, a bottleneck enzyme in the cytoplasmic mevalonate pathway of precursor biosynthesis, whether or not it was combined with IDI (Fig. 5).

**Fig. 5.**
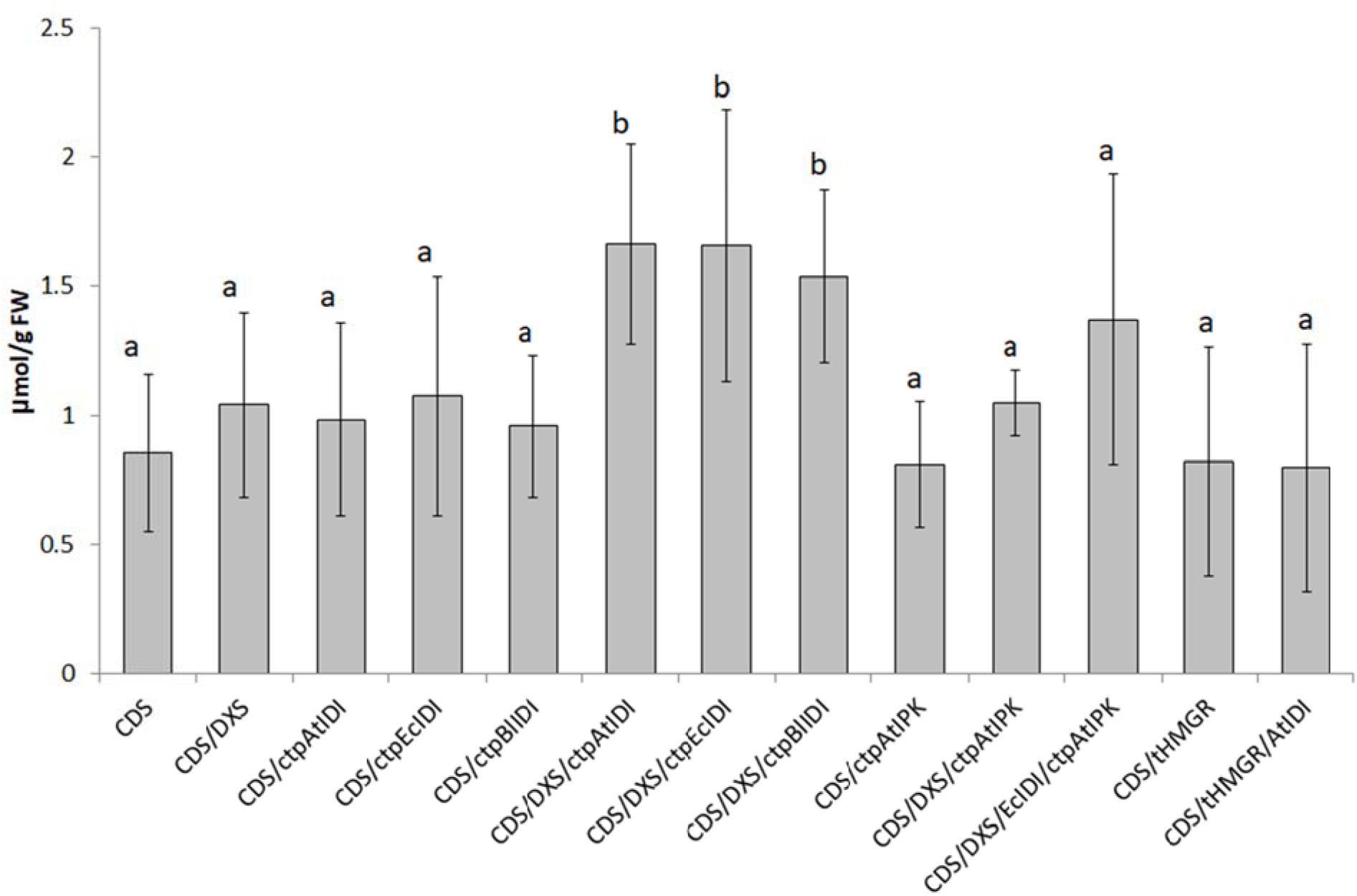
The total concentration of chrysanthemyl-malonyl-glucosides (**1**, **2)** in leaf biomass following transient expression of *Tc*CDS and auxiliary genes in *N. benthamiana*. Different letters indicate significant dif erences **a**t p < 0.01; *n* = 6-27 repeats.

A possibility of increasing the DMAPP pool by recycling monophosphate DMAP by isopentenyl phosphate kinase (IPK) was considered. This activity has been reported in *Arabidopsis*, and overexpressing *At*IPK has led to increased levels of native and heterologous terpenes (Henry et al., 2015;Gutensohn et al., 2021). However, the co-expressed chloroplast-targeted *At*IPK did not significantly enhance *Tc*CDS product accumulation, whether used alone or in combination with DXS or both DXS and IDI (Fig. 5).

The co-expression of DXS and IDI resulted in a statistically significant increase (p < 0.01) in the accumulation of lavandulyl glucosides, too. However, it did not lead to a significant improvement (p > 0.05) in the levels of cyclolavandulyl glucosides produced by the chloroplast-targeted CLDS (Fig. 6). This outcome may be attributed o the initially higher levels of irregular terpene glucoside accumulation following the expression of ctpCLDS.

**Fig. 6.**
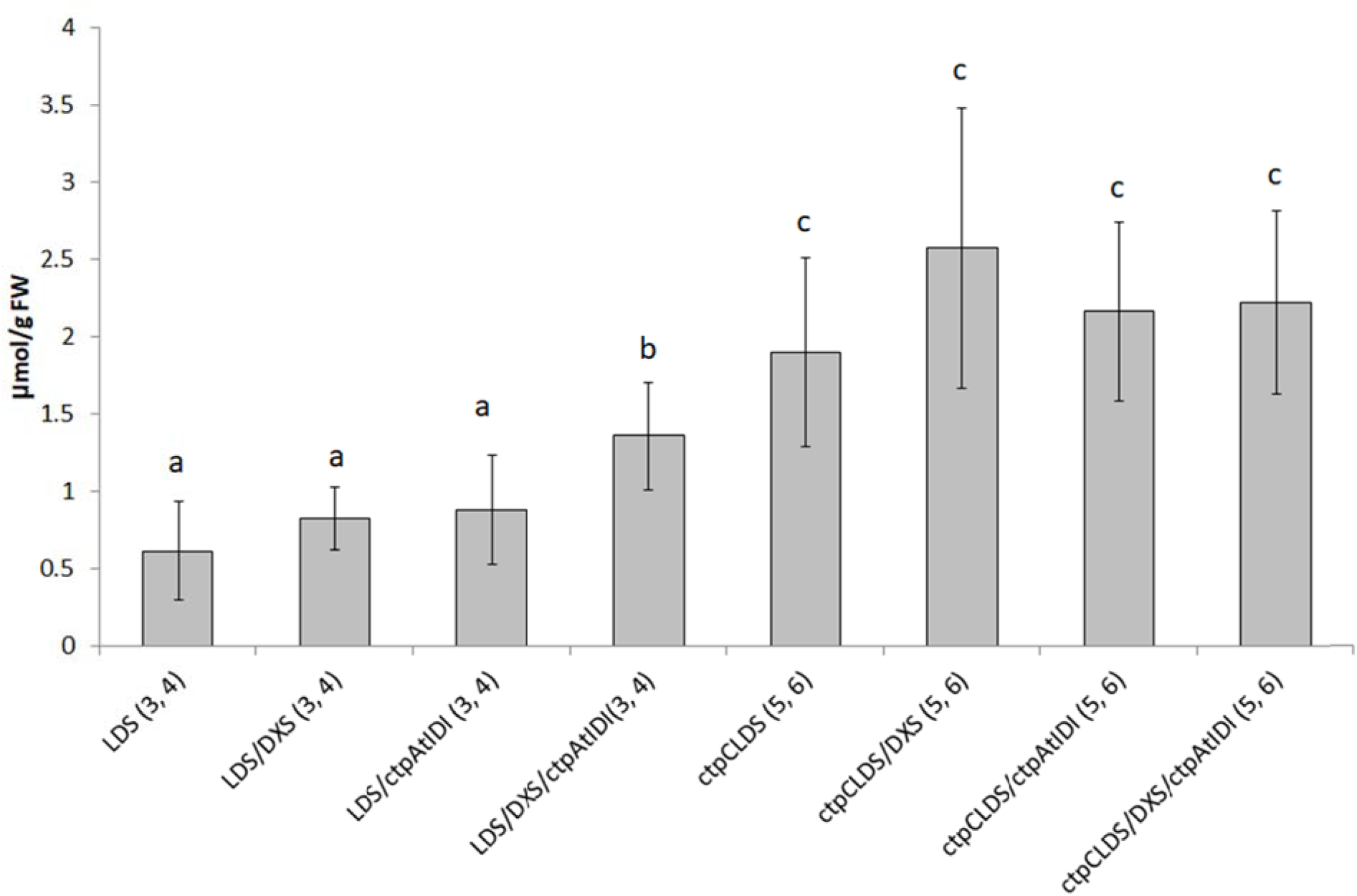
The total concentration of lavandulyl-malonyl-glucosides (**3**, **4)** and cyclolavandulyl-malonyl-glucoside**s** (**5**, **6)** in leaf biomass following transient expression of *Li*LDS or chloroplast-targeted CLDS and auxiliary genes in *N. benthamiana*. Different letters indicate significant differences at p < 0.01; *n* = 6-23 repeats.

### 3.5. Improving cyclolavandulyl skeleton formation in cytoplasm

The accumulation of MVA-pathway derived hete ologous regular sesqui-(Park et al., 2022) and triterpenes (Reed et al., 2017) by transient expression of cytoplasmic enzymes was most effectively supported by co-expression of hydroxymethylglutaryl-CoA reductas (HMGR), particularly the N-truncated s luble version of HMGR1 enzyme from *S. cerevisiae* (tHMGR) (Polakowski et al., 1998). We utilized tHMGR to improve the productivity of CLDS in cytoplasm. Unlike DXS co-expression, inclusion of tHMGR alone resulted in a s atistically significant increase (p < 0.01) in the levels of cyclolavandulyl malonyl-glucosides (**5**, **6**). The addition of IDI or IPK activities did not further enhance the product accumulation. The activity of CLDS in the cytoplasm was not significantly improved (p > 0.05) by the overexpression of bottleneck enzymes of MEP pathway in chloroplasts, such as DXS alone or in combination with ctp*At*IDI (Fig. 7).

**Fig. 7.**
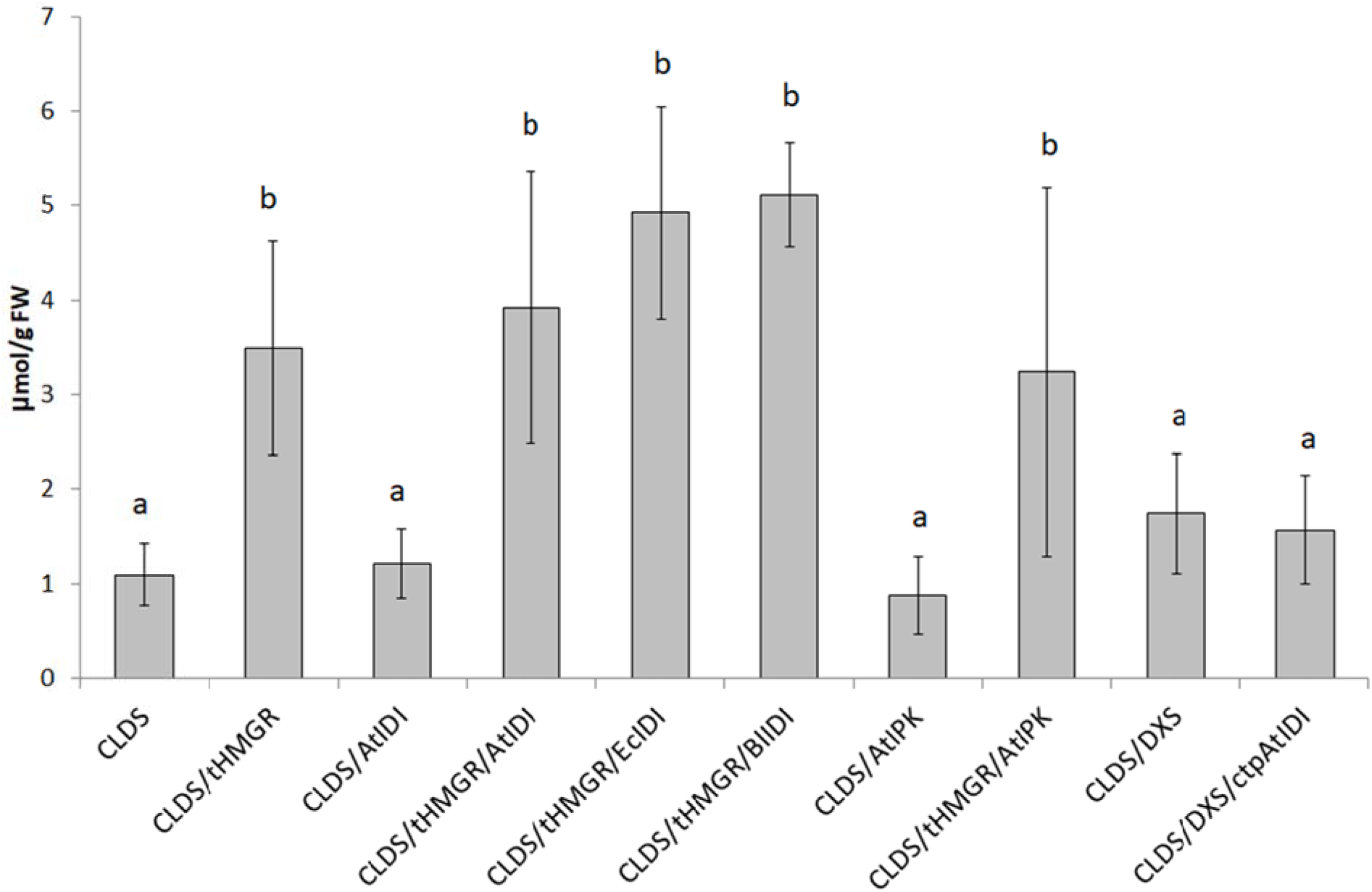
The total concentration of cyclolavandulyl-malonyl-glucosides (**5**, **6)** in leaf biomass following transient expression of CLDS and auxiliary genes in *N. enthamiana*. Different letters indicate significant differences at p < 0.01; *n* = 5-30 repeats.

It is important to note that co**-**expressing CLDS and tHMGR often caused necrosis in the leaf tissue (Fig. S2.50). This phenomenon occurred only i**n** experiments involving tHMGR and cytoplasmic CLDS, with or without additional enzymes. Approximately half of the leaves expressing CLDS in conjunction with tHMGR developed severe necrosis and showed negligible levels of cyclolavandulyl glucosides. These samples were excluded from the calculations of monoterpene glucoside levels presented in Fig. 7.

## 4. Discussion

Considering the benefits of monoterpene glycosylation for easier accumulation and storage in extractable forms within plant cells, we focused our study on developing a platform for producing non-volatile glycosylated irregular monoterpenes in the leaves of *N. benthamiana* after the transient expression of the corresponding IDS genes. Glycosylation is the enzymatic process in which a sugar residue from an activated nucleotide sugar is attached to an acceptor molecule. This process is widely found throughout the plant kingdom. Numerous enzymes, glycosyl transferases and glycosidases, have been identified to facilitate the synthesis and hydrolysis of glycosides. Glycosylation plays several important metabolic roles, such as providing structural support, enabling transport, offering protection, and serving storage functions. For secondary metabolites, many of which are toxic, glucosylation facilitates their transport and storage in vacuoles in an inactive form. Later, through enzymatic hydrolysis, an active aglycone can be released (Kytidou et al., 2020).

Plants produce a wide variety of glycosylated secondary metabolites. Within the monoterpene group, nearly 400 glycosides have been isolated and identified from various plant species over the past few decades (Dembitsky, 2006;Soni et al., 2025). Notably, only 1-2% of these glycosides feature a glucose moiety decorated with a malonyl group. The aglycone portions of these malonyl glycosides consist of regular cyclic monoterpene structures (Yamada et al., 2010;Selenge et al., 2014). These compounds have been isolated from *Dracocephalum foetidum* and *Monarda punctata*, both belonging to the Lamiaceae family, along with various other groups of glycosylated metabolites (Yamada et al., 2010;Selenge et al., 2014). To date, no native malonylated glycosides have been reported in tobacco. Systematic screening of glycosides in tobacco leaf extracts using HPLC-MS has revealed 64 glycosylated structures, which include 39 monosaccharide-linked glycosides, 18 diglycosides, and 7 triglycosides. The aglycone components include flavonoids, coumarins, ionone-related molecules, and sesquiterpene structures (Ding et al., 2015).

The investigation of transgenic tobacco expressing heterologous regular and irregular IDS and terpene synthase genes resulted in a significant increase in the total levels of volatile monoterpenes emitted from the flowers and leaves (Lucker et al., 2004;Yang et al., 2014;Matáeos-Fernández et al., 2023). In one of the early studies, Lucker and colleagues (2004) found no evidence of heterologous monoterpene glycosylation in tobacco cells. However, a later study by Yang and collaborators (2014) demonstrated the presence of chrysanthemyl-malonyl-glucoside derivatives of chrysanthemol and chrysanthemic acid in *N. tabacum* (Yang et al., 2014, Xu et al., 2018b). When the pathway for synthesis of geranic acid was introduced in maize, the most dominant new compound was identified as geranoyl-6*-O-*malonyl-*β-*D-glucopyranoside. It is noteworthy, that this metabolite was not detected in control plants which contained other geraniol derivatives, although at relatively low levels (Yang et al., 2011).

For many monoterpenes, the accumulation of glycosylated forms predominated over that of aglycones. In *Petunia hybrida* plants transformed with linalool synthase, as well as in transgenic *Chrysanthemum morifolium* expressing the *Tc*CDS gene, the synthesized monoterpenes were almost entirely converted and stored in glucosylated form (Lucker et al., 2001;Hu et al., 2018). In transgenic *S. lycopersicum* or *N. benthamiana* that transiently expressed *Tc*CDS along with two oxidoreductases, approximately two-thirds of the produced *trans*-chrysanthemic acid were glycosylated (Xu et al., 2018a;Xu et al., 2018b).

In our experiments, the concentration of terpene monoglucosides prevailed over that of diglucosides, but the ratio varied for different monoterpene moieties, ranging from 3.7 for the compounds **1** and **2** to 1.3 and 1.6 for the compounds **3** and **4** and **5** and **6**, respectively. Notably, the ratio for the compounds **5** and **6** did not differ between cytoplasm and chloroplast-localized IDS, being 1.6 and 1.5, respectively.

The variability in glycoside accumulation among different plant hosts may be influenced by the species-specific activity of glycosylation enzymes. This activity can, in turn, be regulated by the concentration of heterologous monoterpenes and the rate of their biosynthesis. An analysis of the terpenoid database (TPCN) showed a positive correlation between the levels of terpenoids in various plant species and their glycosylation levels (Li et al., 2024). From a plant physiology perspective, glycosylation enhances the water solubility and stability of terpenoids. This process facilitates the transfer and storage of terpenoids in vacuoles and reduces their toxicity to plant cells. During transient expression, the rapid synthesis of heterologous IDS, followed by an intensive formation of products, may stimulate intracellular mechanisms related to metabolite transport and detoxification, further promoting glycosylation.

The reported concentrations of heterologous non-volatile monoterpenes in the tissues of transgenic plants range from 0.07 to 0.7 μmol g^-1^ FW (10 to 121 μg g^-1^ FW) (Lucker et al., 2001;Yang et al., 2011;Xu et al., 2018a). Transient expression of *Tc*CDS, in combination with two oxidoreductases, led to the accumulation of *trans*-chrysanthemic acid in esterified forms, including malonylated glucosides, at a level of 1.1 μmol g^-1^ FW (190 μg g^-1^ FW) which constituted 58% of the total monoterpene content (Xu et al., 2018b). In our experiments, transient co-expression of plant-derived naturally plastid-targeted *Li*LDS and *Tc*CDS with auxiliary substrate-producing enzymes DXS and IDI resulted in accumulation of lavandulyl and chrysanthemyl glycosides at levels of 1.4 ± 0.3 and 1.6 ± 0.4 μmol g^-1^ FW, respectively. Prokaryotic CLPS fused with an artificial chloroplast-targeting peptide allowed for the accumulation of 2.2 ±0.7 μmol g^-1^ FW of cyclolavandulol in the form of non-volatile mono- and diglucosides. Initially higher level of the latter compound was not statistically significantly increased further by enhancing the metabolic flux through the MEP pathway and intensifying the isomerization of IPP to DMAPP by overexpression of DXS and IDI. Subsequent experiments on cytoplasmic formation of irregular monoterpenes have indicated that the observed product concentrations approach the tolerance threshold for plant cells regarding these compounds.

Co-expression of CLDS and tHMGR in the cytoplasm led to the highest concentration of cyclolavandulyl glucosides in the tissue. Significantly better performance of the MVA pathway for terpene production has been repeatedly demonstrated in microbial hosts. Not only in eukaryotic chassis, e.g. *S. cerevisiae*, the highest titers were achieved by engineering the native MVA pathway, but the most effectively producing bacterial strains were constructed by introducing the heterologous enzymes of the MVA route (Yang et al., 2025). In plants, the overexpression of the enzymes of both pathways increases the terpene production (Park et al., 2022). Investigation of substrate availability in plastids and cytoplasm of tomato fruit cell revealed that MVA-derived sesquiterpenes were formed in considerably less amounts than monoterpenes originated from MEP. However, the overexpression of HMGR elevated the product level more efficiently than DXS (Gutensohn et al., 2021). In our experiments, the co-expression of HMGR with CLDS often caused leaf necrosis which was not observed for each of these enzymes alone or for any other protein combination. We assume that either the amount of produced monoterpenes surpasses the capacities of the glycosylation system leading to the accumulation of toxic free alcohols or the malonyl glucosides are also toxic at high concentrations. The mean level of cyclolavandulyl glucosides after co-expression of CLDS and HMGR reached 3.9 ± 1.5 μmol g^-1^ FW; the best samples contained 6.6 μmol g^-1^ FW of these compounds. These values represent the highest reported concentrations of irregular monoterpenes isolated from a plant system.

## 5. Conclusions

To promote sustainable plant-based production of irregular monoterpenes, we established a transient procedure to enhance the biosynthetic flux in *Nicotiana benthamiana* toward DMAPP by coexpressing IDS genes and the genes of substrate-producing enzymes of MEP or MVA pathways. That allowed the accumulation of substantial amounts of non-volatile glucosylated derivatives of irregular monoterpenes. We identified five new metabolites of lavandulol, cyclolavandulol and chrysanthemol in addition to the previously described chrysanthemyl-6*-O-*malonyl*-β-*D-glucopyranoside. Alleviating the bottlenecks in the biosynthesis of the substrate for the key enzymes resulted in the highest yields of irregular monoterpenes reported for the plant systems.

## Acknowledgements

The authors are grateful to the mass spectrometry core facility team of the Chemistry Department at the Technical University of Darmstadt for measurements of the ESI/APCI spectra. The authors thank Prof. Christina M. Thiele, Technische Universität Darmstadt, for measurement time on the 700 MHz NMR spectrometer. The research was supported by the European Research Area Cofund Action ‘ERACoBioTech’ under Horizon 2020, the project SUSPHIRE (Sustainable Production of Pheromones for Insect Pest Control in Agriculture). The financial support from the German Federal Ministry of Education and Research (BMBF), grant number 031B0605, and from the German Research Foundation (DFG), grant number INST 163/444-1 FUGG (QTOF MS) is highly appreciated. YVS acknowledges the support by DFG, grant number 516587177.

