## Supplementary material for "Efficient accumulation of new irregular monoterpene malonyl glucosides in *Nicotiana benthamiana* achieved by co-expression of isoprenyl diphosphate synthases and substrate-producing enzymes"

### Supporting Information

Figure S2.21 $^{1}H$ -$^{1}H$ CLIP-COSY spectrum of 3 in DMSO-$d_6$. 32

Figure S2.22 $^{1}H$ -$^{13}C$ HMBC spectrum of 3 in DMSO-$d_6$. 33

Figure S2.23 $^{1}H$ -$^{1}H$ TOCSY spectrum of 3 in DMSO-$d_6$. 34

Figure S2.24 1H -1H NOESY spectrum of 3 in DMSO-d6. 35

Figure S2.25 COSY (bold lines) and key HMBC (arrows) correlations (A), and NOESY correlations (B) of 4 in DMSO-$d_6$. 36

Figure S2.26 $^{1}H$ NMR spectrum of 4 in DMSO-$d_6$. 37

Figure S2.27 $^{13}C$ NMR spectrum of 4 in DMSO-$d_6$. 38

Figure S2.28 $^{1}H$ -$^{13}C$ HSQC spectrum of 4 in DMSO-$d_6$. 39

Figure S2.29 $^{1}H$ -$^{1}H$ CLIP-COSY spectrum of 4 in DMSO-$d_6$. 40

Figure S2.30 $^{1}H$ -$^{13}C$ HMBC spectrum of 4 in DMSO-$d_6$. 41

Figure S2.31 $^{1}H$ -$^{1}H$ TOCSY spectrum of 4 in DMSO-$d_6$. 42

Figure S2.32 $^{1}H$ -$^{1}H$ NOESY spectrum of 4 in DMSO-$d_6$. 43

Figure S2.33 COSY (bold lines) and key HMBC (arrows) correlations (A), and NOESY correlations (B) of 5 in DMSO-$d_6$. 44

Figure S2.34 $^{1}H$ NMR spectrum of 5 in DMSO-$d_6$. 45

Figure S2.35 $^{13}C$ NMR spectrum of 5 in DMSO-$d_6$. 46

Figure S2.36 $^{1}H$ -$^{13}C$ HSQC spectrum of 5 in DMSO-$d_6$. 47

Figure S2.37 $^{1}H$ -$^{1}H$ CLIP-COSY spectrum of 5 in DMSO-$d_6$. 48

Figure S2.38 $^{1}H$ -$^{13}C$ HMBC spectrum of 5 in DMSO-$d_6$. 49

Figure S2.39 $^{1}H$ -$^{1}H$ TOCSY spectrum of 5 in DMSO-$d_6$. 50

Figure S2.40 $^{1}H$ -$^{1}H$ NOESY spectrum of 5 in DMSO-$d_6$. 51

Figure S2.41 COSY (bold lines) and key HMBC (arrows) correlations (A), and NOESY correlations (B) of 6 in DMSO-$d_6$. 52

Figure S2.42 $^{1}H$ NMR spectrum of 6 in DMSO-$d_6$. 53

Figure S2.43 $^{13}C$ NMR spectrum of 6 in DMSO-$d_6$. 54

Figure S2.44 $^{1}H$ -$^{13}C$ HSQC spectrum of 6 in DMSO-$d_6$. 55

Figure S2.45 $^{1}H$ -$^{1}H$ CLIP-COSY spectrum of 6 in DMSO-$d_6$. 56

Figure S2.46 $^{1}H$ -$^{13}C$ HMBC spectrum of 6 in DMSO-$d_6$. 57

Figure S2.47 $^{1}H$ -$^{1}H$ TOCSY spectrum of 6 in DMSO-$d_6$. 58

Figure S2.48 $^{1}H$ -$^{1}H$ NOESY spectrum of 6 in DMSO-$d_6$. 59

Figure S2.49 Exemplary HPLC separation of the reaction mixture after MPP-derivatization. 60

Figure S2.50 *N. benthamiana* plants expressing StCLDS alone and in combination with tHMGR. 61

### S1 Experimental section

#### S1.1 Chemicals and equipment

Commercial laboratory chemicals, reagents and components of nutrition medium were purchased from the companies Merck (Germany), Roth (Germany), AppliChem (Germany) etc. 3-Methyl-1-phenyl-2-pyrazolin-5-on was ordered from Merck (Germany). Solvents for liquid chromatography were obtained from VWR (Germany) and Roth (Germany).

#### S1.2 Coding sequences

The sequence encoding LDS from *Lavandula x intermedia* was synthesized according to the GenBank entry JX985358.1. The coding sequence of CDS from *Tanacetum cinerariifolium* was designed based on the GenBank entry JX913536.1, with modifications on the N-terminus of the chloroplast transit peptide (submitted to the GenBank, waiting for Acc. No.). The coding sequence of CLDS from *Streptomyces* sp. CL190 was codon-optimized for *Nicotiana*:

gcattcctgcagttacctacagtttctccaataggaagaattaattctaagcttcttattccctctttctcgtcactccgaacc
tgcaccagcactgcaggaaaacgaatcggagagaggccgaagcttgaacgcttctcccgttcgattcctaagtgct
gtttagcaggtgcagaaaccgaaattgacgaggtgacacccaatcacgtcgcaattatcatagacggacacagaa
agtgggcaaagagtagaggggttacagttcaagagggtcatcaaaccggtgttaacaattggaagcatatcatttcc
cgggcttctcaactcggaatcaagcttctcacaatctgggccttatccccgcagaattttaatcgctctaaaatggaagt
tgacttcttgatgaggatttacgaagatttcctacgatccgatgtcaaagaacttgtcaccagccaacaagacattcaat
tttctgcgattggtgacaaatcaagactcccagaatatctacaagacgcaatatcctacgctgaaggactgagccag
gctaacaagggcatgcatttcatactggcggtagcgtacggcggacgtgaagacatcgtggaggcggccagaaa
gatcgcagccaaagtcgaacacggtatcctacgaccagacgacatcgacgaagctacgttcgaacaacatctgat
gaccaacatcacaaaattcccaagcccggatctactgattagggcagccggtgaacagaggctcagcaacttctttc
tatggcagttgcccttcacagaattctactttacgcctaaattgtttccggattttggcgaggcggatcttctcgacgcgctt
gcctcttaccgctgcaggtatagaggcttcggtgaacgaaaaggaattcatgaa

The sequences encoding IDI from *E. coli* and *Bacillus licheniformis* were synthesized according to the GenBank entries AP026104.1 (range 3694692 to 3695234) and CP014781.1 (range 2437718 to 2438764), respectively. The sequences encoding the mature protein IDI1 without the chloroplast targeting peptide and IPK were amplified from the cDNA of *Arabidopsis thaliana* using the primers based on the GenBank sequences NM_121649.6 for IDI1

Fw: 5’-ACTACGTCTCACTCGAGCCGCTTTCTCAGCCGTC-3’;

Rv: 5’-TCTACGTCTCACTCGCTGCGAGCTTGTGAATGG-3’

and AY150412.1 for IPK

Fw: 5’-ACTACGTCTCACTCGAGCCGAGCTGAATATTTCC-3’;

Rv: 5’-TCTACGTCTCACTCGCTGCCTTTGAGAATCTGATG-3’).

All coding sequences were adapted for the GoldenBraid cloning system by deleting the start and stop codons and addition of nucleotides AGCC on 5’-end and GCAG on 3’-end.

**Table S1.1** Combinations of the GoldenBraid vectors used in experiments on the biosynthesis of irregular monoterpene malonyl glucosides in *N. benthamiana*. P35S - 35S CaMV promoter; HisTnos - nopaline synthase gene terminator from *A. tumefaciens* preceded by a sequence encoding 8 histidine residues; ctp – sequence encoding an artificial chloroplast transit peptide

| IDS | Auxiliary genes | Product |
|---|---|---|
| omega1<br>P35S::*Tc*CDS::HisTnos<br>P35S::p19::Tnos | alpha2 | 1, 2 |
| omega1<br>P35S::*Tc*CDS::HisTnos<br>P35S::p19::Tnos | alpha2<br>P35S::*Sl*DXS2::HisTnos | 1, 2 |
| omega1<br>P35S::*Tc*CDS::HisTnos<br>P35S::p19::Tnos | alpha1<br>P35S::ctp::*At*IDI1::HisTnos | 1, 2 |
| omega1<br>P35S::*Tc*CDS::HisTnos<br>P35S::p19::Tnos | alpha1<br>P35S::ctp::*Ec*IDI::HisTnos | 1, 2 |
| omega1<br>P35S::*Tc*CDS::HisTnos<br>P35S::p19::Tnos | alpha1<br>P35S::ctp::*Bl*IDI::HisTnos | 1, 2 |
| omega1<br>P35S::*Tc*CDS::HisTnos<br>P35S::p19::Tnos | alpha1<br>P35S::ctp::*At*IPK::HisTnos | 1, 2 |
| omega1<br>P35S::*Tc*CDS::HisTnos<br>P35S::p19::Tnos | alpha2<br>P35S::tHMGR::HisTnos | 1, 2 |
| omega1<br>P35S::*Tc*CDS::HisTnos<br>P35S::p19::Tnos | omega2<br>P35S::ctp::*At*IDI1::HisTnos<br>P35S::*Sl*DXS2::HisTnos | 1, 2 |

| | | |
|---|---|---|
| omega1<br>P35S::*Tc*CDS::HisTnos<br>P35S::p19::Tnos | omega2<br>P35S::ctp::*Ec*IDI::HisTnos<br>P35S::*Sl*DXS2::HisTnos | 1, 2 |
| omega1<br>P35S::*Tc*CDS::HisTnos<br>P35S::p19::Tnos | omega2<br>P35S::ctp::*Bl*IDI::HisTnos<br>P35S::*Sl*DXS2::HisTnos | 1, 2 |
| omega1<br>P35S::*Tc*CDS::HisTnos<br>P35S::p19::Tnos | omega2<br>P35S::ctp::*At*IPK::HisTnos<br>P35S::*Sl*DXS2::HisTnos | 1, 2 |
| omega1<br>P35S::*Tc*CDS::HisTnos<br>P35S::p19::Tnos | omega2<br>P35S::*At*IDI1::HisTnos<br>P35S::tHMGR::HisTnos | 1, 2 |
| omega1<br>P35S::*Tc*CDS::HisTnos<br>P35S::p19::Tnos | omega2<br>P35S::ctp::*Ec*IDI::HisTnos<br>P35S::*Sl*DXS2::HisTnos<br>alpha1<br>P35S::ctp::*At*IPK::HisTnos | 1, 2 |
| omega1<br>P35S::*Li*LDS::HisTnos<br>P35S::p19::Tnos | alpha2 | 3,4 |
| omega1<br>P35S::*Li*LDS::HisTnos<br>P35S::p19::Tnos | alpha2<br>P35S::*Sl*DXS2::HisTnos | 3,4 |
| omega1<br>P35S::*Li*LDS::HisTnos<br>P35S::p19::Tnos | alpha1<br>P35S::ctp::*At*IDI1::HisTnos | 3,4 |
| omega1<br>P35S::*Li*LDS::HisTnos<br>P35S::p19::Tnos | omega2<br>P35S::ctp::*At*IDI1::HisTnos<br>P35S::*Sl*DXS2::HisTnos | 3,4 |
| omega1<br>P35S::ctp::*St*CLDS::HisTnos | alpha2 | 5,6 |

| | | |
|---|---|---|
| P35S::p19::Tnos | | |
| omega1<br>P35S::ctp::*St*CLDS::HisTnos<br>P35S::p19::Tnos | alpha2<br>P35S::*Sl*DXS2::HisTnos | 5,6 |
| omega1<br>P35S::ctp::*St*CLDS::HisTnos<br>P35S::p19::Tnos | alpha1<br>P35S::ctp::*At*IDI1::HisTnos | 5,6 |
| omega1<br>P35S::ctp::*St*CLDS::HisTnos<br>P35S::p19::Tnos | omega2<br>P35S::ctp::*At*IDI1::HisTnos<br>P35S::*Sl*DXS2::HisTnos | 5,6 |
| omega1<br>P35S::*St*CLDS::HisTnos<br>P35S::p19::Tnos | alpha2 | 5,6 |
| omega1<br>P35S::*St*CLDS::HisTnos<br>P35S::p19::Tnos | alpha2<br>P35S::tHMGR::HisTnos | 5,6 |
| omega1<br>P35S::*St*CLDS::HisTnos<br>P35S::p19::Tnos | alpha1<br>P35S::*At*IDI1::HisTnos | 5,6 |
| omega1<br>P35S::*St*CLDS::HisTnos<br>P35S::p19::Tnos | alpha1<br>P35S::*At*IPK::HisTnos | 5,6 |
| omega1<br>P35S::*St*CLDS::HisTnos<br>P35S::p19::Tnos | omega2<br>P35S::*At*IDI1::HisTnos<br>P35S::tHMGR::HisTnos | 5,6 |
| omega1<br>P35S::*St*CLDS::HisTnos<br>P35S::p19::Tnos | omega2<br>P35S::*Ec*IDI::HisTnos<br>P35S::tHMGR::HisTnos | 5,6 |
| omega1<br>P35S::*St*CLDS::HisTnos<br>P35S::p19::Tnos | omega2<br>P35S::*Bl*IDI::HisTnos<br>P35S::tHMGR::HisTnos | 5,6 |

| | | |
|---|---|---|
| omega1<br>P35S::*St*CLDS::HisTnos<br>P35S::p19::Tnos | omega2<br>P35S::*At*IPK::HisTnos<br>P35S::tHMGR::HisTnos | 5,6 |
| omega1<br>P35S::*St*CLDS::HisTnos<br>P35S::p19::Tnos | alpha2<br>P35S::*Sl*DXS2::HisTnos | 5,6 |
| omega1<br>P35S::*St*CLDS::HisTnos<br>P35S::p19::Tnos | omega2<br>P35S::ctp::*At*IDI1::HisTnos<br>P35S::*Sl*DXS2::HisTnos | 5,6 |

**Table S1.2** $^{1}H$ and $^{13}C$ NMR data of 1-6 in DMSO-$d_5$. Values are in ppm. The multiplicities and coupling constants (*J* in Hz) are in parentheses.

| Position | $^{1}$H NMR | | | | | | $^{13}$C NMR | | | | | |
|---|---|---|---|---|---|---|---|---|---|---|---|---|
| | **1** | **2** | **3** | **4** | **5** | **6** | **1** | **2** | **3** | **4** | **5** | **6** |
| 1 | 3.82 (dd, 1H, *J* = 11, 8.7 Hz); 3.43 (dd, 1H, *J* = 11, 5.7 Hz) | 3.79 (t, *J* = 10.7, 8.1 Hz, 1H); 3.51 – 3.47 (m, 1H) | 3.44 (dd, 1H, *J* = 9.8, 6.6 Hz); 3.63 (dd, 1H, *J* = 9.8, 7.1 Hz) | 3.44 (dd, 1H, *J* = 9.4, 6.2 Hz); 3.66 (dd, 1H, *J* = 9.4, 7.4 Hz) | 4.23 (d, *J* = 11.4 Hz, 1H); 3.97 (d, *J* = 11.4 Hz, 1H) | 4.13 – 4.08 (m, 2H) | 68.56 | 68.83 | 70.80 | 70.74 | 66.73 | 67.35 |
| 2 | 0.75 (dt, *J* = 8.5, 5.6 Hz, 1H) | 0.81 – 0.77 (m, 1H) | 2.30 (m, 1H) | 2.29 (m, 1H) | - | - | 31.8 | 31.64 | 46.45 | 46.54 | 124.58 | 124.99 |
| 3 | 1.10 – 1.07 (m, 1H) | 1.11 – 1.08 (m, 1H) | 1.98 (m, 1H); 2.2 (dt, *J* = 13.5, 6.4 Hz, 1H) | 1.99 (m, 1H); 2.2 (dt, *J* = 13.7, 6.4 Hz, 1H) | 2.14 (d, *J* = 17.4 Hz, 1H); 1.95 (d, *J* = 17.4 Hz, 1H) | 2.14 (d, *J* = 17.2 Hz, 1H); 2.01 (d, *J* = 17.2 Hz, 1H) | 27.56 | 27.62 | 28.21 | 28.14 | 24.95 | 24.68 |
| 4 | 4.86 (d, *J* = 8.2 Hz, 1H) | 4.86 (d, *J* = 8.2 Hz, 1H) | 5.02 (t, 1H, *J* = 7.0 Hz) | 5.02 (t, 1H, *J* = 7.1 Hz) | 1.28 (t, *J* = 6.6 Hz, 2H) | 1.27 (t, *J* = 6.5 Hz, 2H) | 123.91 | 123.94 | 122.48 | 122.57 | 35.12 | 35.16 |
| 5 | - | - | - | - | - | - | 131.71 | 131.49 | 131.36 | 131.22 | 28.77 | 28.74 |
| 6 | 1.62 (s, 3H) | 1.63 (s, 3H) | 1.63 (s, 3H) | 1.63 (s, 3H) | 1.73 (s, 2H) | 1.71 (s, 2H) | 18.12 | 18.14 | 25.63 | 25.62 | 45.58 | 45.58 |
| 7 | 1.65 (s, 3H) | 1.65 (s, 3H) | 1.56 (s, 3H) | 1.56 (s, 3H) | - | - | 25.43 | 25.42 | 17.77 | 17.78 | 130.76 | 129.85 |
| 8 | - | - | - | - | 1.62 (s, 3H) | 1.60 (s, 3H) | 21.79 | 21.53 | 145.49 | 145.58 | 18.82 | 18.87 |
| 9 | 1.07 (s, 3H) | 1.07 (s, 3H) | 4.73 (s, 1H); 4.69 (s, 1H) | 4.73 (s, 1H); 4.68 (s, 1H) | 0.86 (s, 3H) | 0.85 (s, 3H) | 21.22 | 21.14 | 111.81 | 111.68 | 28.08 | 28.02 |
| 10 | 0.99 (s, | 0.98 (s, | 1.64 (s, | 1.64 (s, | 0.86 (s, | 0.85 (s, | 22.37 | 22.36 | 20.12 | 20.19 | 28.12 | 28.25 |

| | | | | | | | | | | | | |
|---|---|---|---|---|---|---|---|---|---|---|---|---|
| | 3H) | 3H) | 3H) | 3H) | 3H) | 3H) | | | | | | |
| Glc-1’ | 4.13 (d, *J* = 7.9 Hz, 1H) | 4.31 (d, *J* = 7.7 Hz, 1H) | 4.14 (d, *J* = 7.8 Hz, 1H) | 4.31 (d, *J* = 7.6 Hz, 1H) | 4.03 (d, *J* = 7.9 Hz, 1H) | 4.24 (d, *J* = 7.7 Hz, 1H) | 102.27 | 101.02 | 102.92 | 101.36 | 100.43 | 99.69 |
| 2’ | 2.96 (t, *J* = 9.4, 7.9 Hz, 1H) | 3.30 – 3.26 (m, 1H) | 2.95 (t, *J* = 9.3, 7.8 Hz, 1H) | 3.31 – 3.27 (m, 1H) | 2.97 (t, *J* = 9.1, 8.0 Hz, 1H) | 3.31 – 3.28 (m, 1H) | 73.36 | 81.68 | 73.4 | 81.4 | 73.22 | 81.51 |
| 3’ | 3.14 (t, *J* = 9.4, 8.9 Hz, 1H) | 3.38 (t, *a* = 9.1 Hz, 1H) | 3.14 (t, *J* = 8.9 Hz, 1H) | 3.39 (t, *J* = 9.0 Hz, 1H) | 3.11 (t, *J* = 9.4, 8.8 Hz, 1H) | 3.41 – 3.35 (m, 1H) | 76.41 | 75.51 | 76.33 | 75.58 | 76.47 | 75.76 |
| 4’ | 3.09 (t, *J* = 9.3, 8.9 Hz, 1H) | 3.16 – 3.13 (m, 1H) | 3.09 (t, *J* = 10.6, 9.3 Hz, 1H) | 3.19 – 3.16 (m, 1H) | 3.07 (t, *J* = 9.4, 8.8 Hz, 1H) | 3.18 – 3.14 (m, 1H) | 70.10 | 69.72 | 70.0 | 69.87 | 70.20 | 69.99 |
| 5’ | 3.31 – 3.27 (m, 1H) | 3.36 – 3.33 (m, 1H) | 3.30 (m, 1H) | 3.37 – 3.34 (m, 2H) | 3.24 (ddd, *J* = 8.9, 6.9, 1.8 Hz, 1H) | 3.34 – 3.31 (m, 1H) | 73.69 | 73.71 | 73.75 | 73.53 | 73.89 | 73.63 |
| 6’ | 4.16 – 4.05 (m, 2H) | 4.14 – 4.10 (m, 1H); 4.07 (d, *J* = 11.6 Hz, 1H) | 4.08-4.12 (m, 2H) | 4.16 – 4.06 (m, 2H) | 4.18 (dd, *J* = 11.8, 1.8 Hz, 1H); 4.06 (dd, *J* = 11.8, 6.9 Hz, 1H) | 4.15 (d, *J* = 11.6 Hz, 0H); 4.09 (dd, *J* = 11.6, 6.5 Hz, 1H) | 63.51 | 63.29 | 63.43 | 63.33 | 63.58 | 63.30 |
| Malonyl $C=O_a$ | - | - | - | - | - | - | 169.19 | 169.72 | 169.7 | 169.60 | 169.31 | 169.54 |
| $CH_2$ | 2.94 – 2.91 (m, 2H) | 2.92 – 2.88 (m, 2H) | 2.92 – 2.88 (m, 2H) | 2.95 – 2.90 (m, 2H) | 2.96 – 2.94 (m, 2H) | 2.94 – 2.90 (m, 2H) | 45.61 | 45.98 | 45.86 | 45.69 | 45.32 | 45.63 |
| $C=O_b$ | - | - | - | - | - | - | 168.43 | 167.19 | 167.63 | 168.15 | 168.08 | 168.13 |
| Glc-1’’ | | 4.40 (d, *J* = 7.9 Hz, 1H) | | 4.38 (d, *J* = 7.7 Hz, 1H) | | 4.39 (d, *J* = 7.8 Hz, 1H) | | 103.9 | | 104.11 | | 104.01 |
| 2’’ | | 2.99 (t, *J* = 8.3 Hz, 2H) | | 2.98 (t, *J* = 8.4 Hz, 1H) | | 2.98 (t, *J* = 8.3 Hz, 1H) | | 74.92 | | 75.10 | | 75.07 |
| 3’’ | | 3.10 – | | 3.17 – | | 3.17 – | | 76.06 | | 76.01 | | 76.01 |

| | | | | | | |
|---|---|---|---|---|---|---|
| | 3.16 (m, 1H) | 3.10 (m, 1H) | 3.11 (m, 1H) | | | |
| 4’’ | 3.10 – 3.16 (m, 1H) | 3.17 – 3.10 (m, 1H) | 3.15 – 3.11 (m, 1H) | 69.76 | 69.77 | 69.72 |
| 5’’ | 3.06 – 3.03 (m, 1H) | 3.03 – 3.00 (m, 1H) | 3.05 – 3.02 (m, 1H) | 77.02 | 77.10 | 77.01 |
| 6’’ | 3.62 (d, *J* = 11.9 Hz, 1H); 3.51 – 3.47 (m, 1H) | 3.61 (d, *J* = 11.5 Hz, 1H); 3.50 – 3.48 (m, 1H) | 3.60 (d, *J* = 11.6 Hz, 1H); 3.51 – 3.47 (m, 1H) | 60.72 | 60.75 | 60.8 |

#### S2 Results

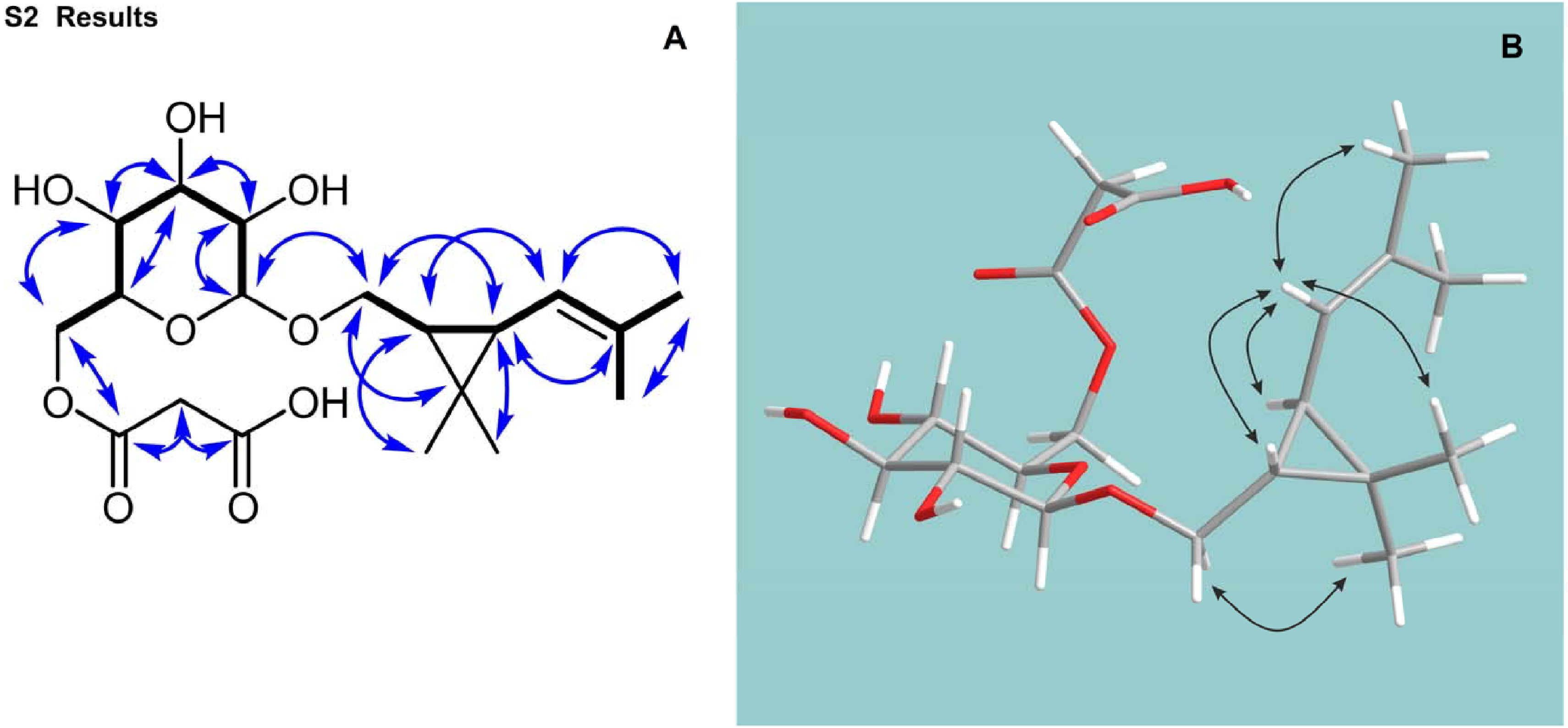

**Figure S2.1** COSY (bold lines) and key HMBC (arrows) correlations (A), and NOESY correlations (B) of **1** in DMSO-$d_6$.

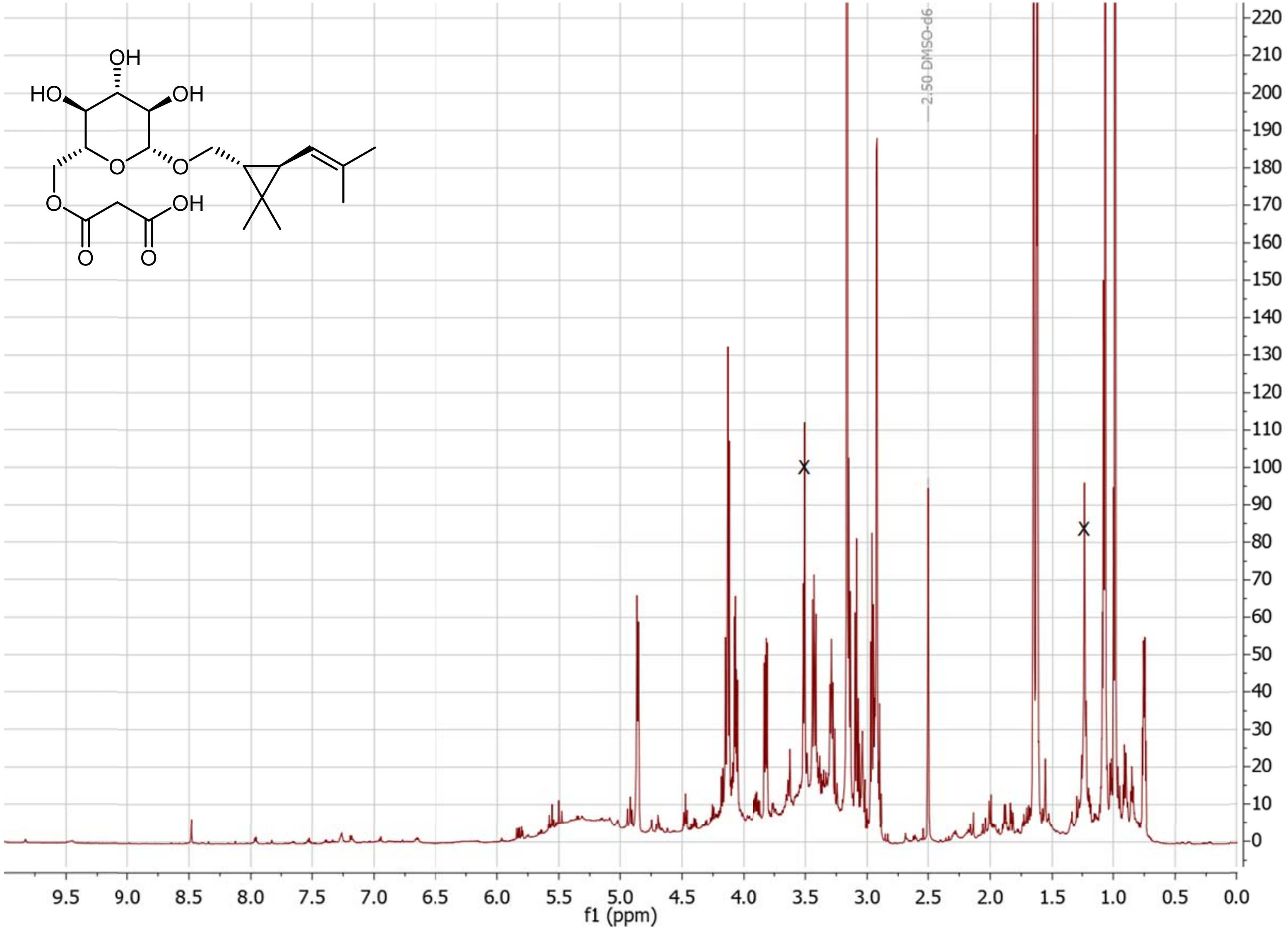

**Figure S2.2** $^{1}$H NMR spectrum of **1** in DMSO-$d_6$. The signals representing residual sample impurities are crossed out.

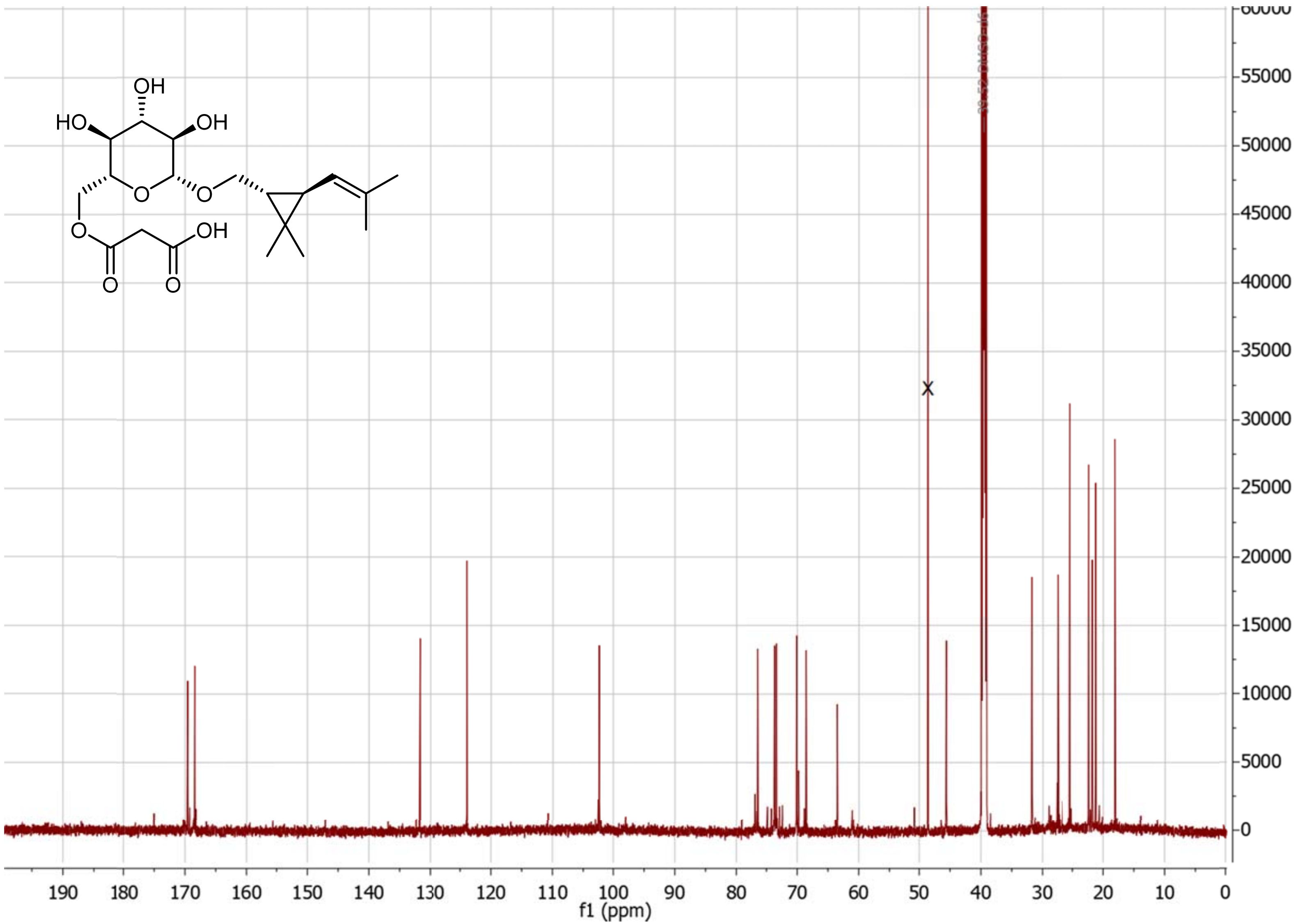

**Figure S2.3** $^{13}$C NMR spectrum of **1** in DMSO-$d_6$. The signals representing residual sample impurities are crossed out.

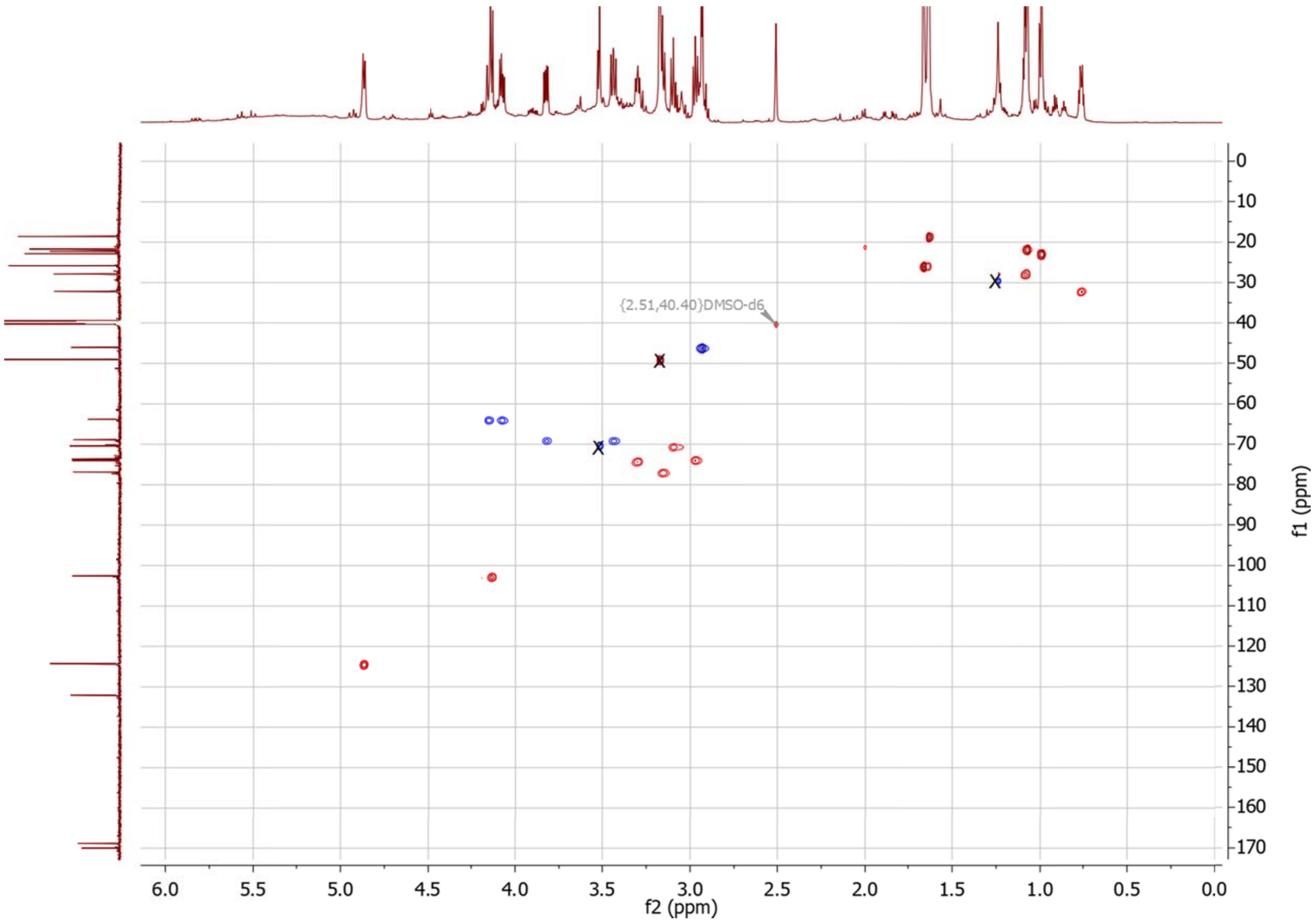

**Figure S2.4** $^{1}H$ -$^{13}C$ HSQC spectrum of **1** in DMSO-$d_6$. The signals representing residual sample impurities are crossed out.

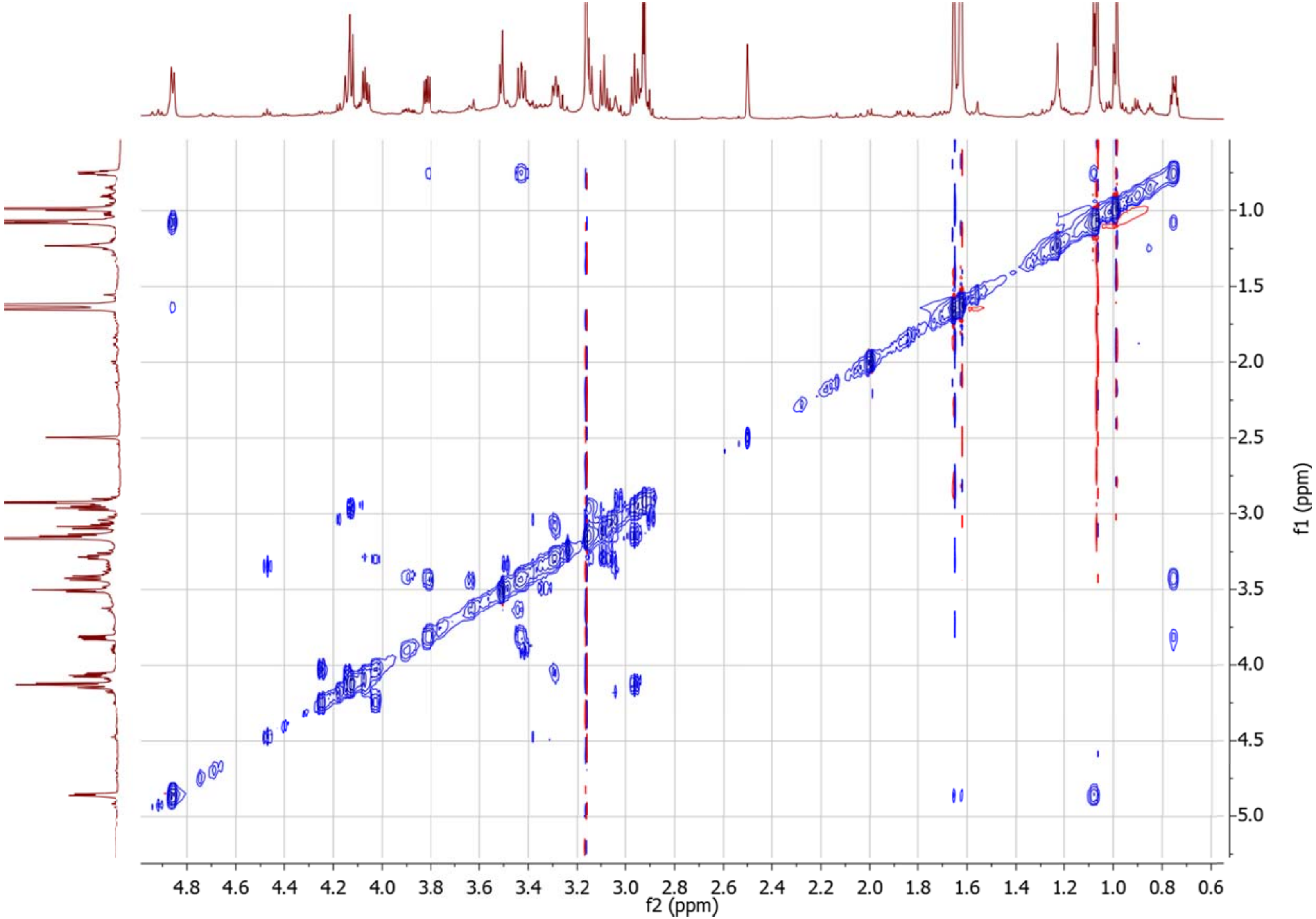

**Figure S2.5** $^1H$ -$^1H$ CLIP-COSY spectrum of **1** in DMSO-$d_6$.

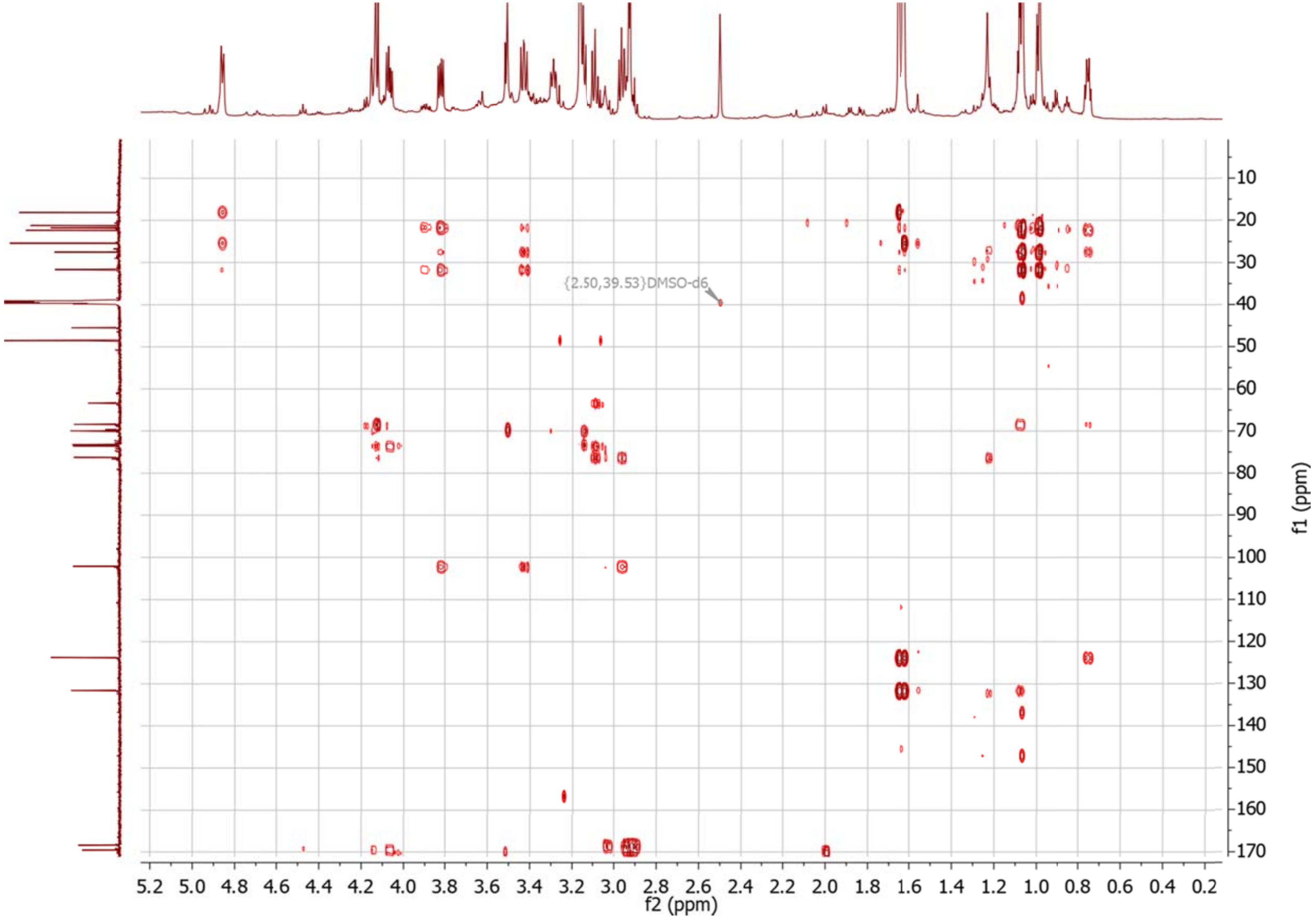

**Figure S2.6** $^{1}H$ -$^{13}C$ HMBC spectrum of **1** in DMSO-$d_6$.

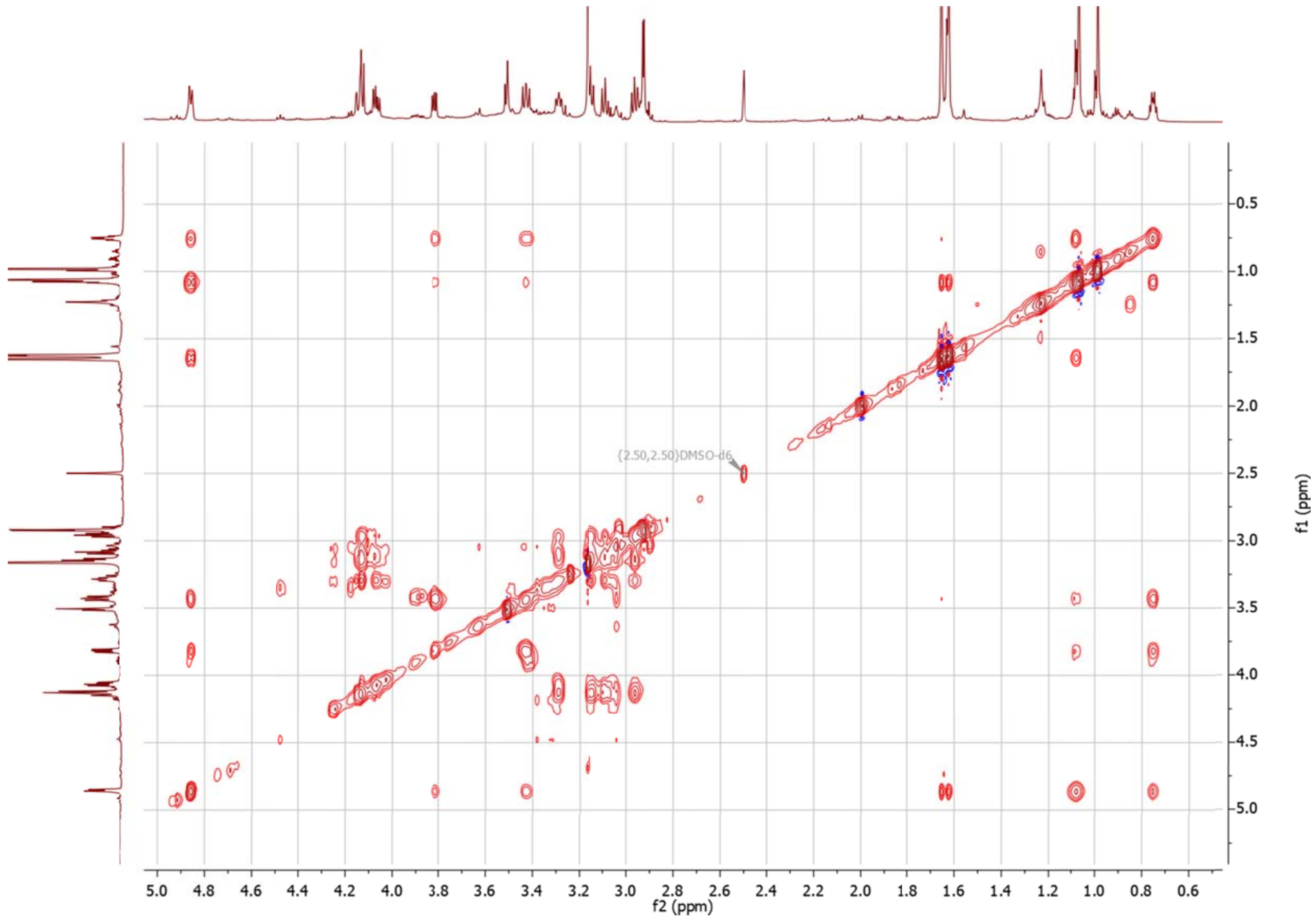

**Figure S2.7** $^1$H -$^1$H TOCSY spectrum of **1** in DMSO-$d_6$.

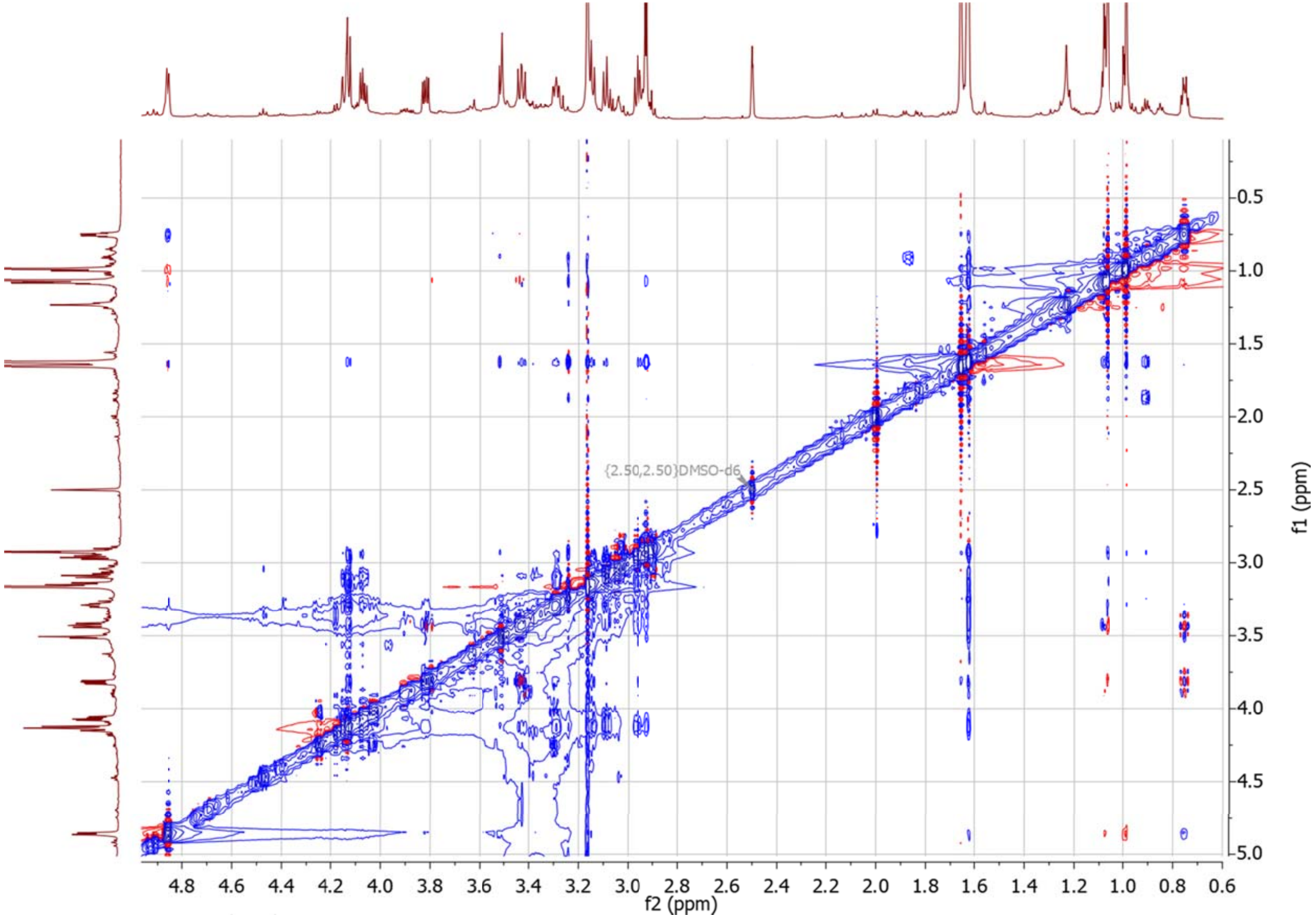

**Figure S2.8** $^{1}H$ -$^{1}H$ NOESY spectrum of **1** in DMSO-$d_6$.

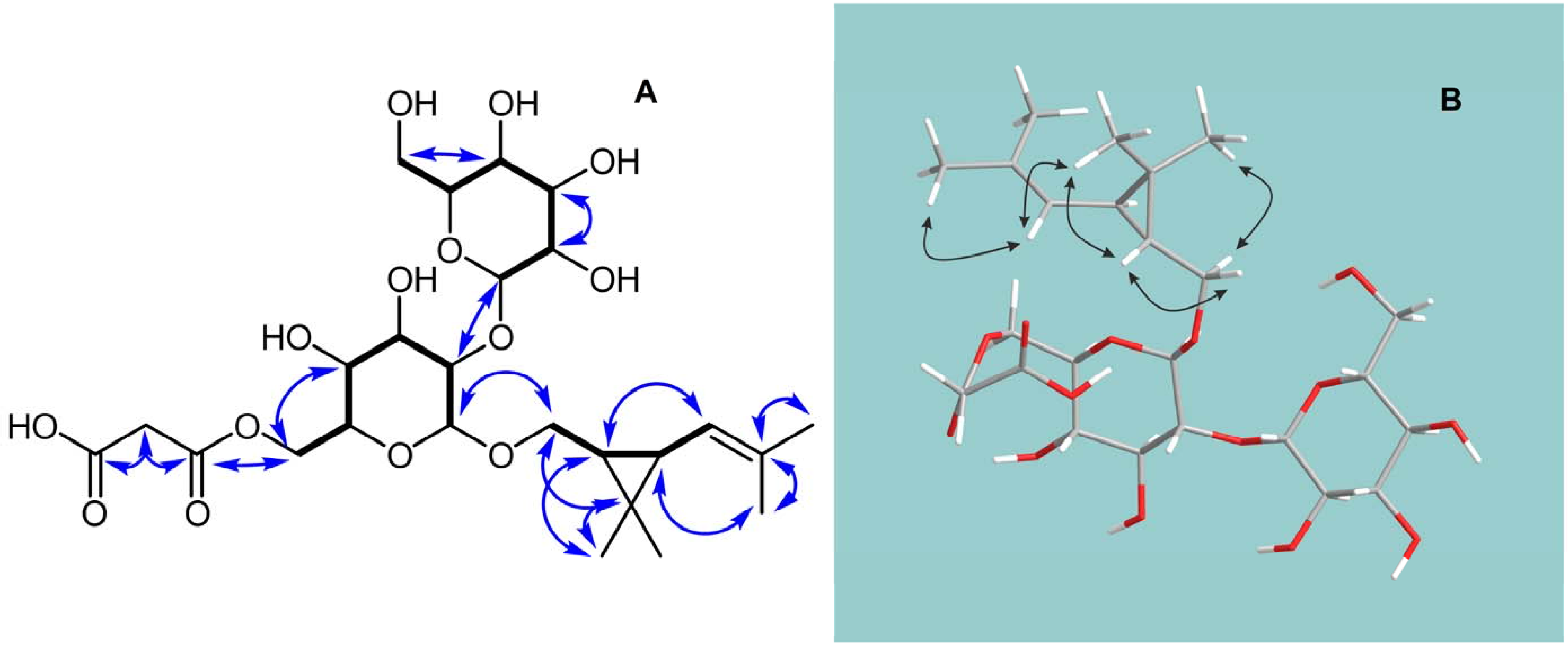

**Figure S2.9** COSY (bold lines) and key HMBC (arrows) correlations (A), and NOESY correlations (B) of **2** in DMSO-$d_6$.

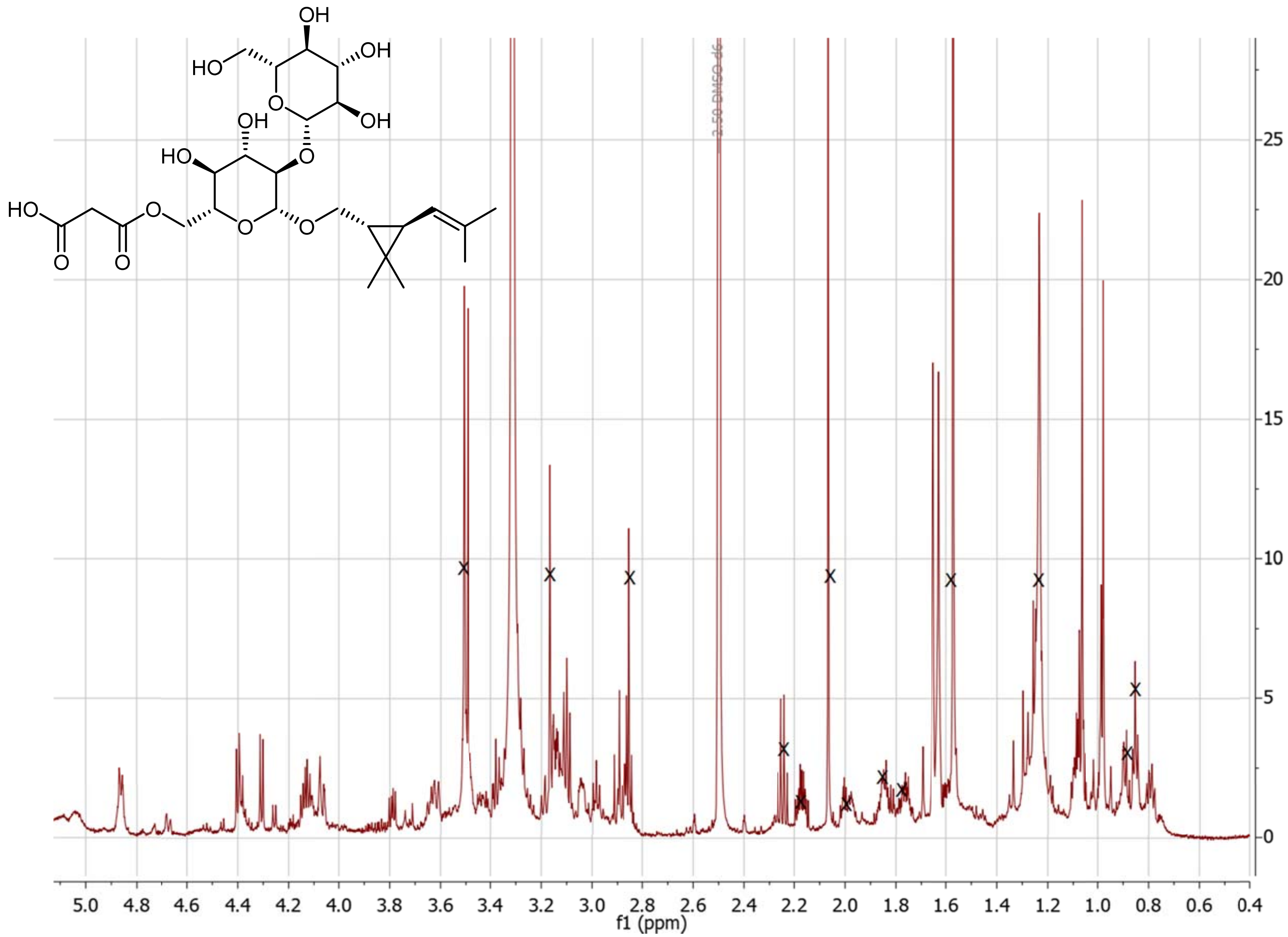

**Figure S2.10** $^{1}$H NMR spectrum of **2** in DMSO-$d_6$. The signals representing residual sample impurities are crossed out.

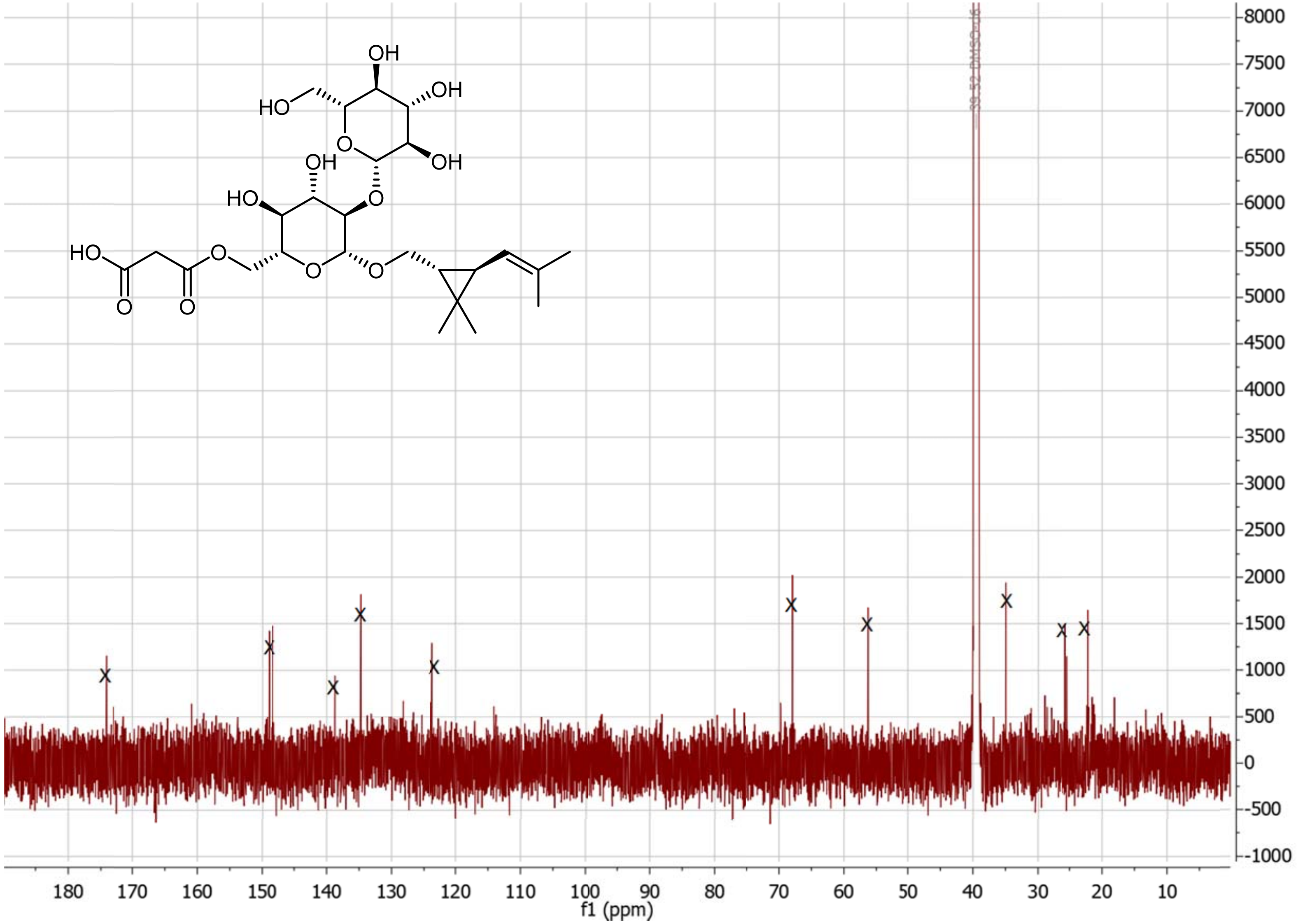

**Figure S2.11** $^{13}$C NMR spectrum of **2** in DMSO-$d_6$. The signals representing residual sample impurities are crossed out.

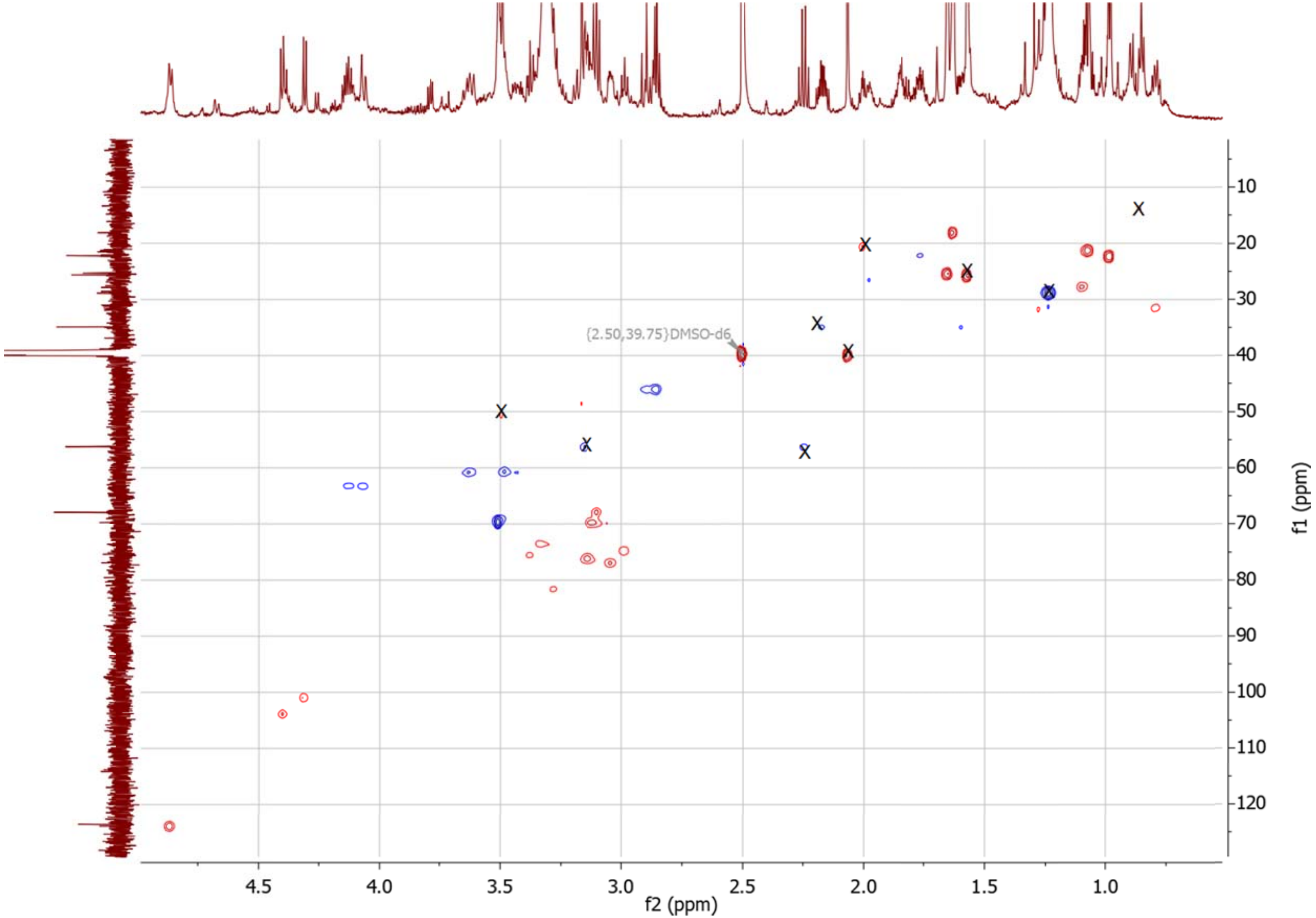

**Figure S2.12** $^{1}$H -$^{13}$C HSQC spectrum of **2** in DMSO-$d_6$. The signals representing residual sample impurities are crossed out.

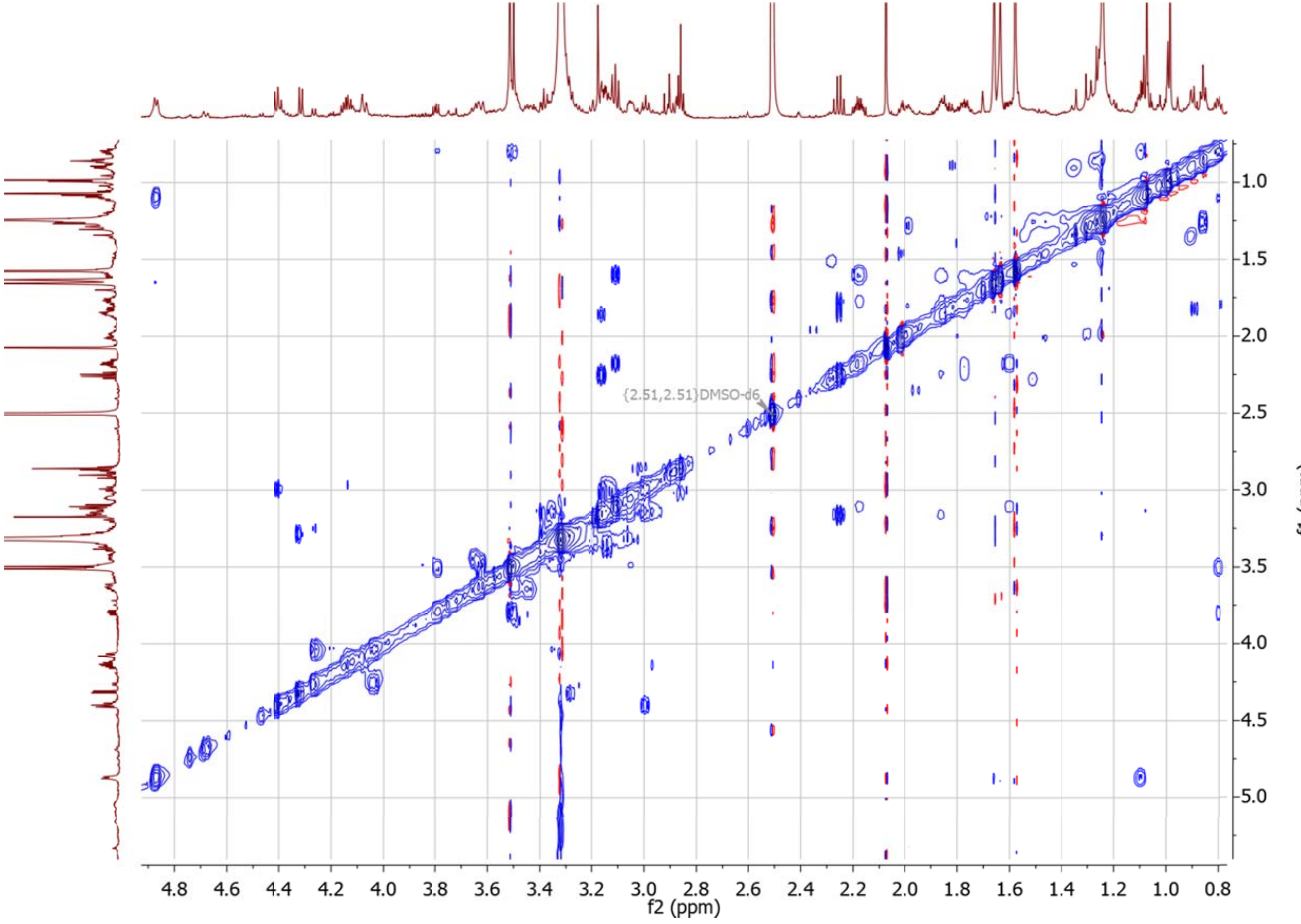

**Figure S2.13** $^{1}$H -$^{1}$H CLIP-COSY spectrum of **2** in DMSO-$d_6$.

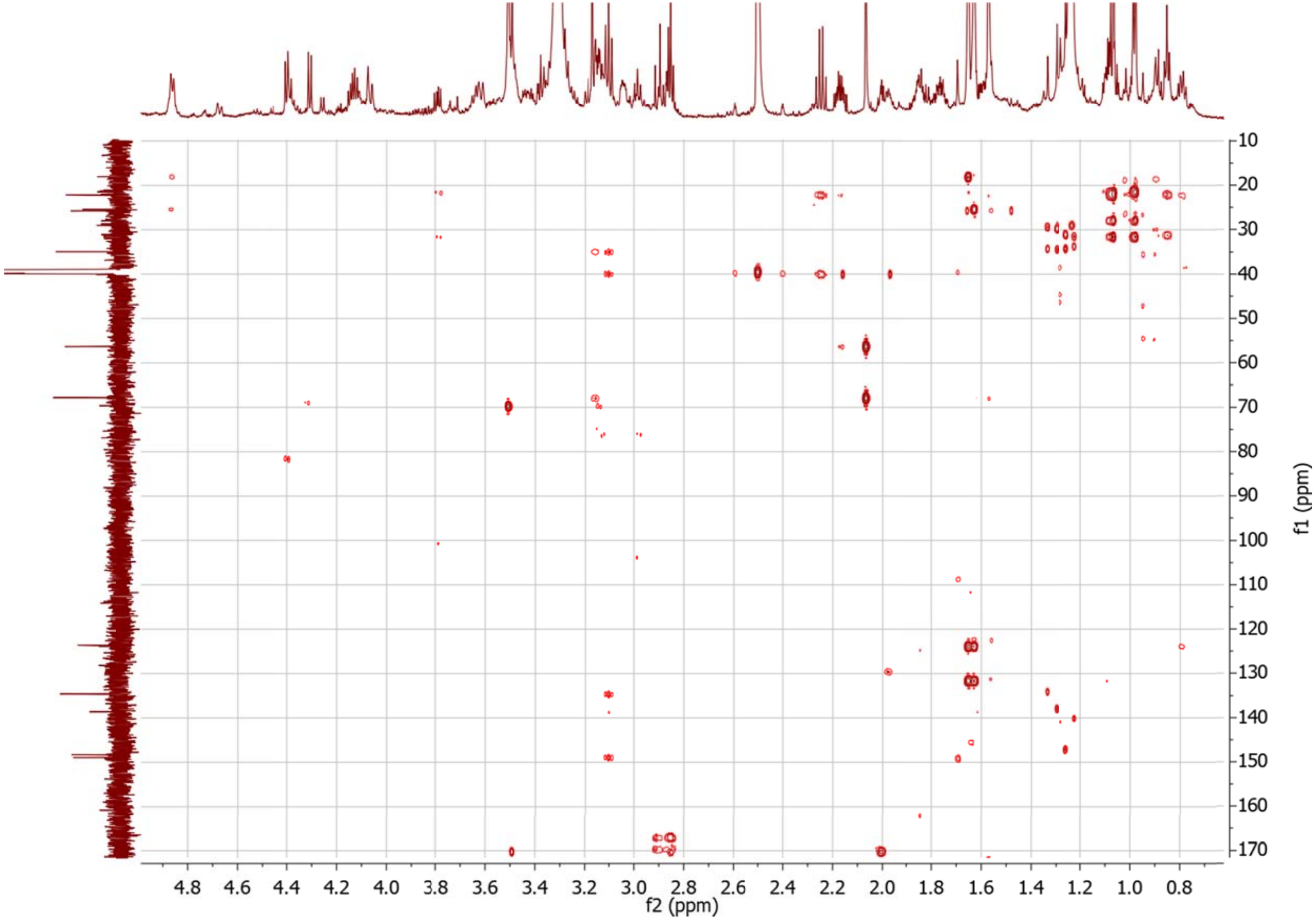

**Figure S2.14** $^{1}$H -$^{13}$C HMBC spectrum of **2** in DMSO-$d_6$.

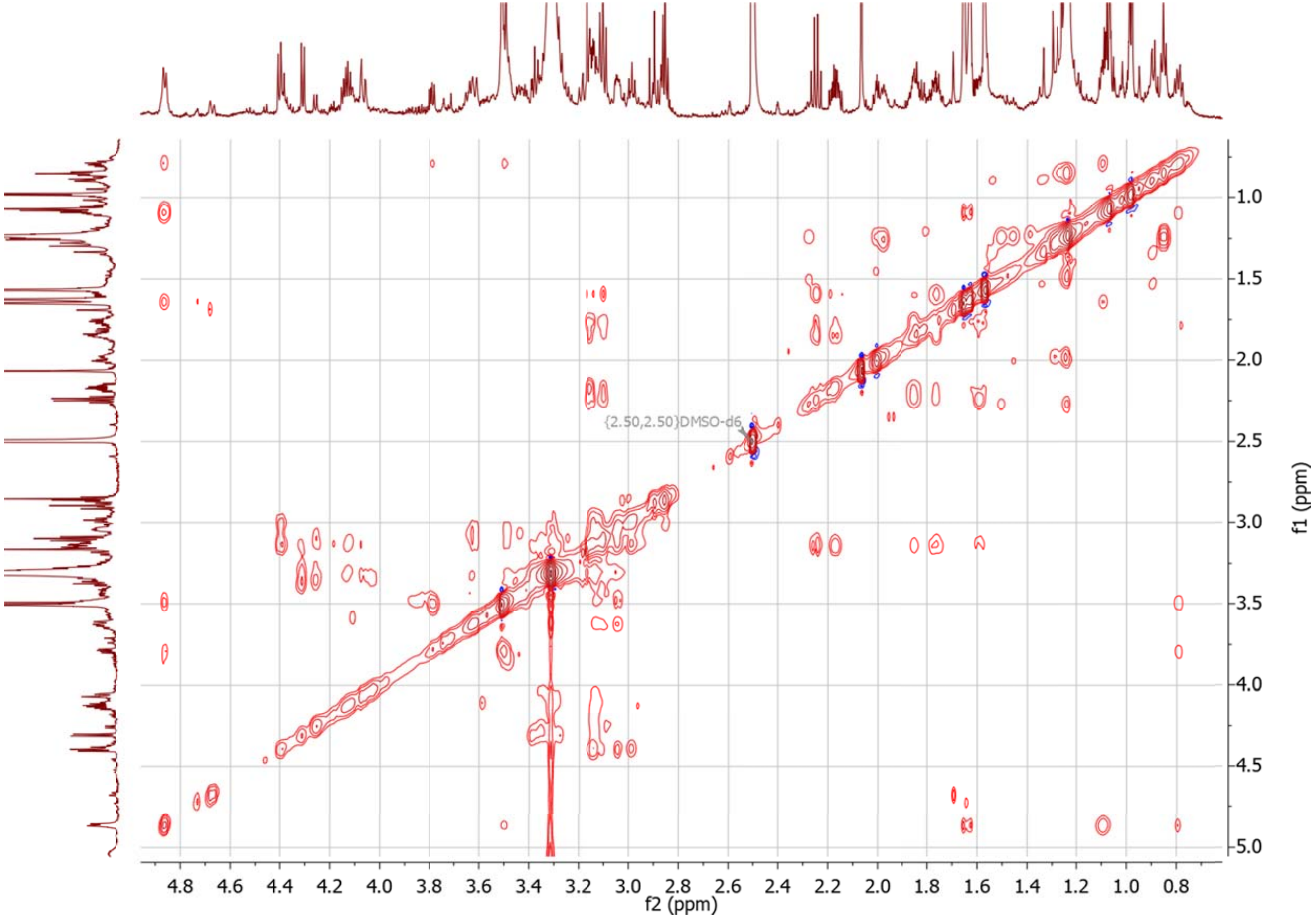

**Figure S2.15** [1]H -[1]H TOCSY spectrum of **2** in DMSO-$d_6$.

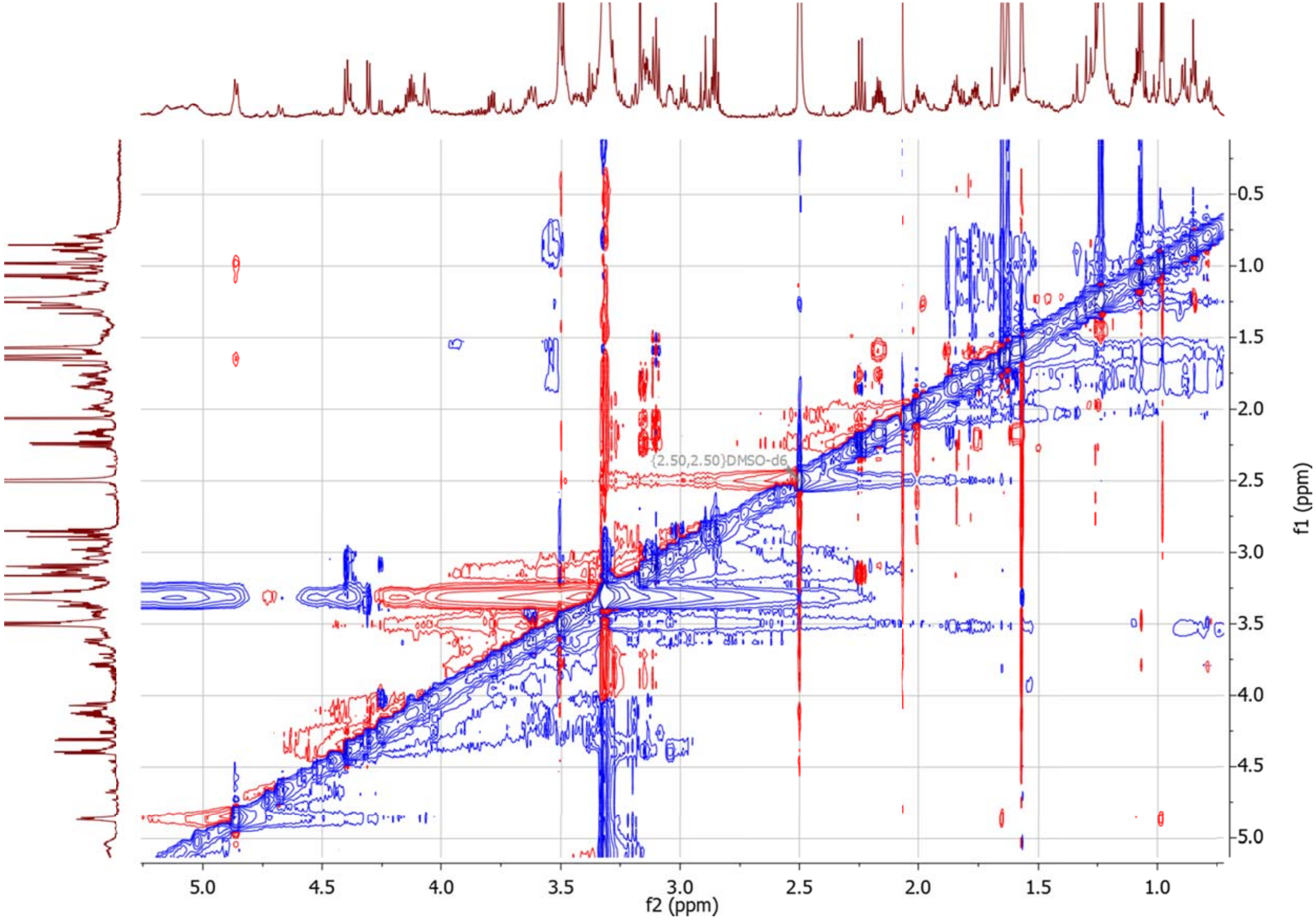

**Figure S2.16** $^{1}$H -$^{1}$H NOESY spectrum of **2** in DMSO-$d_6$.

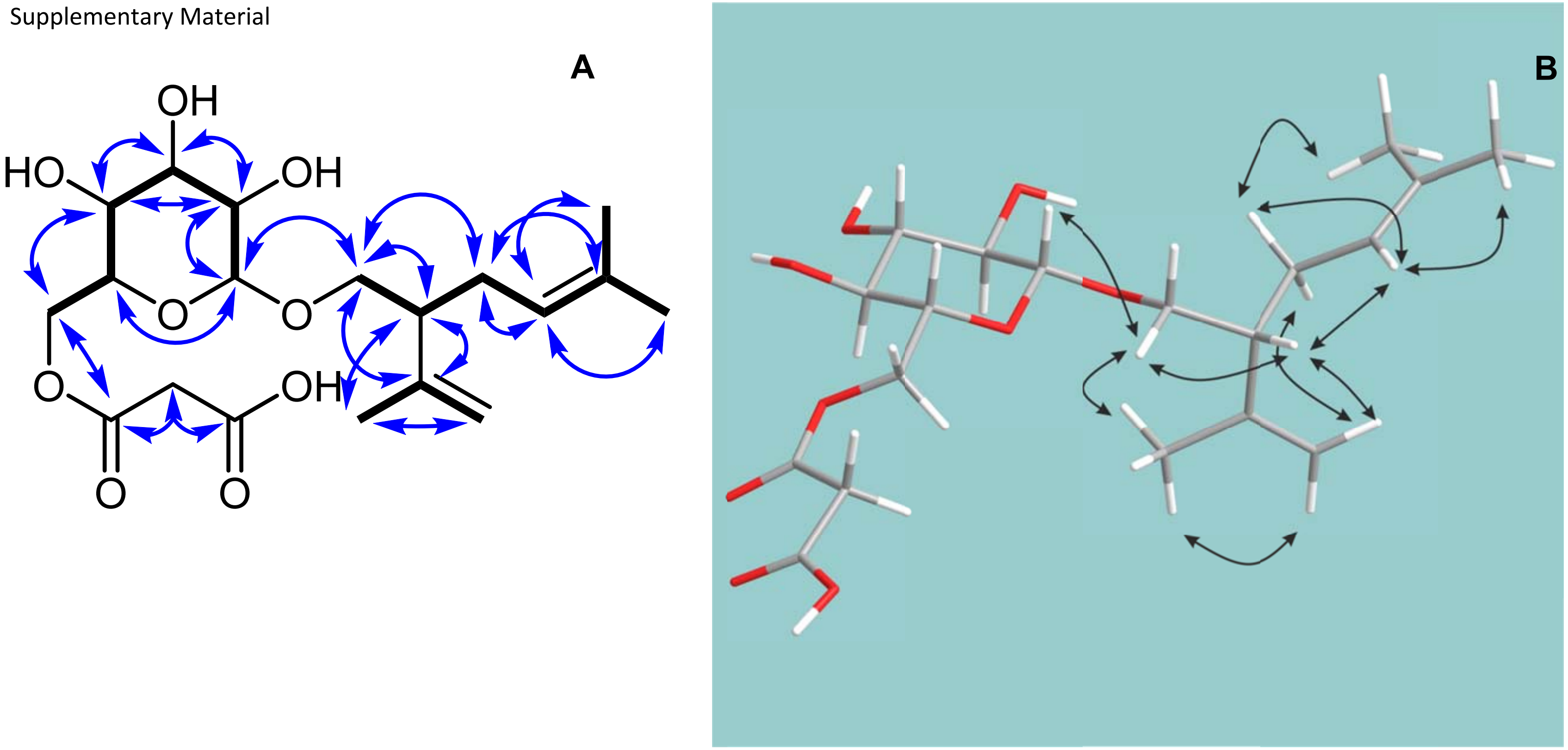

**Figure S2.17** COSY (bold lines) and key HMBC (arrows) correlations (A), and NOESY correlations (B) of **3** in DMSO-$d_6$.

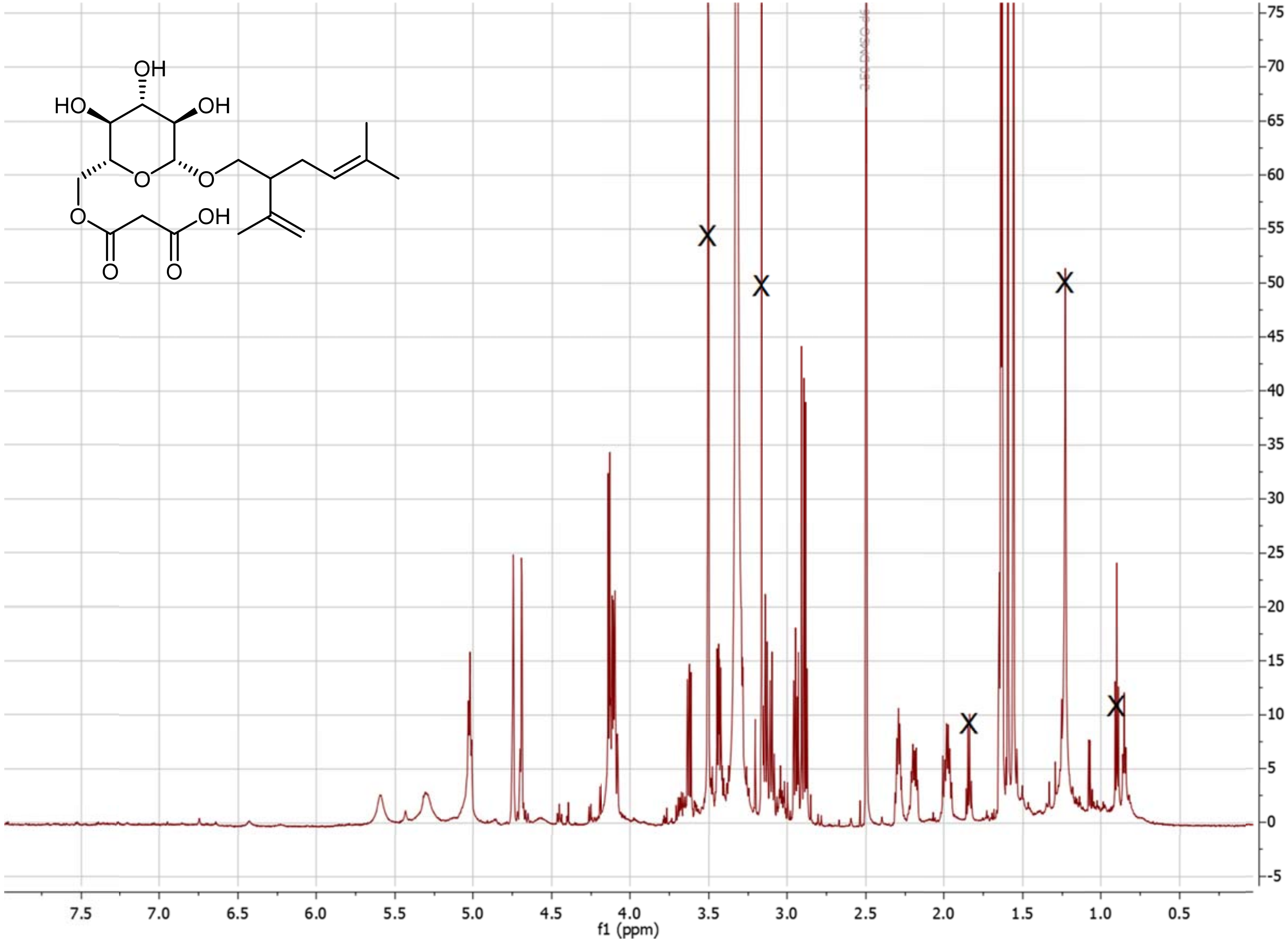

**Figure S2.18** $^{1}$H NMR spectrum of **3** in DMSO-$d_6$. The signals representing residual sample impurities are crossed out.

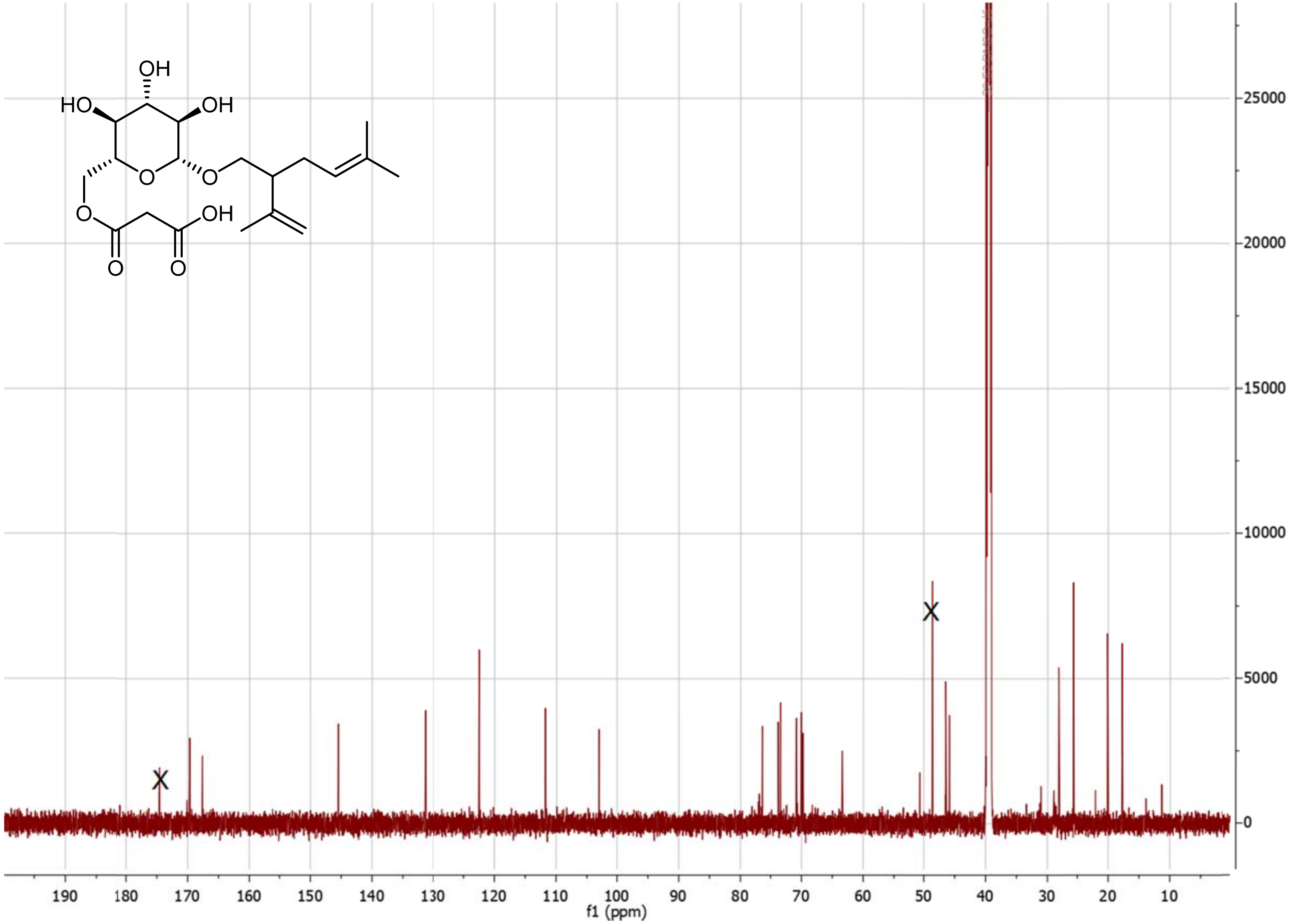

**Figure S2.19** $^{13}C$ NMR spectrum of **3** in DMSO-$d_6$. The signals representing residual sample impurities are crossed out.

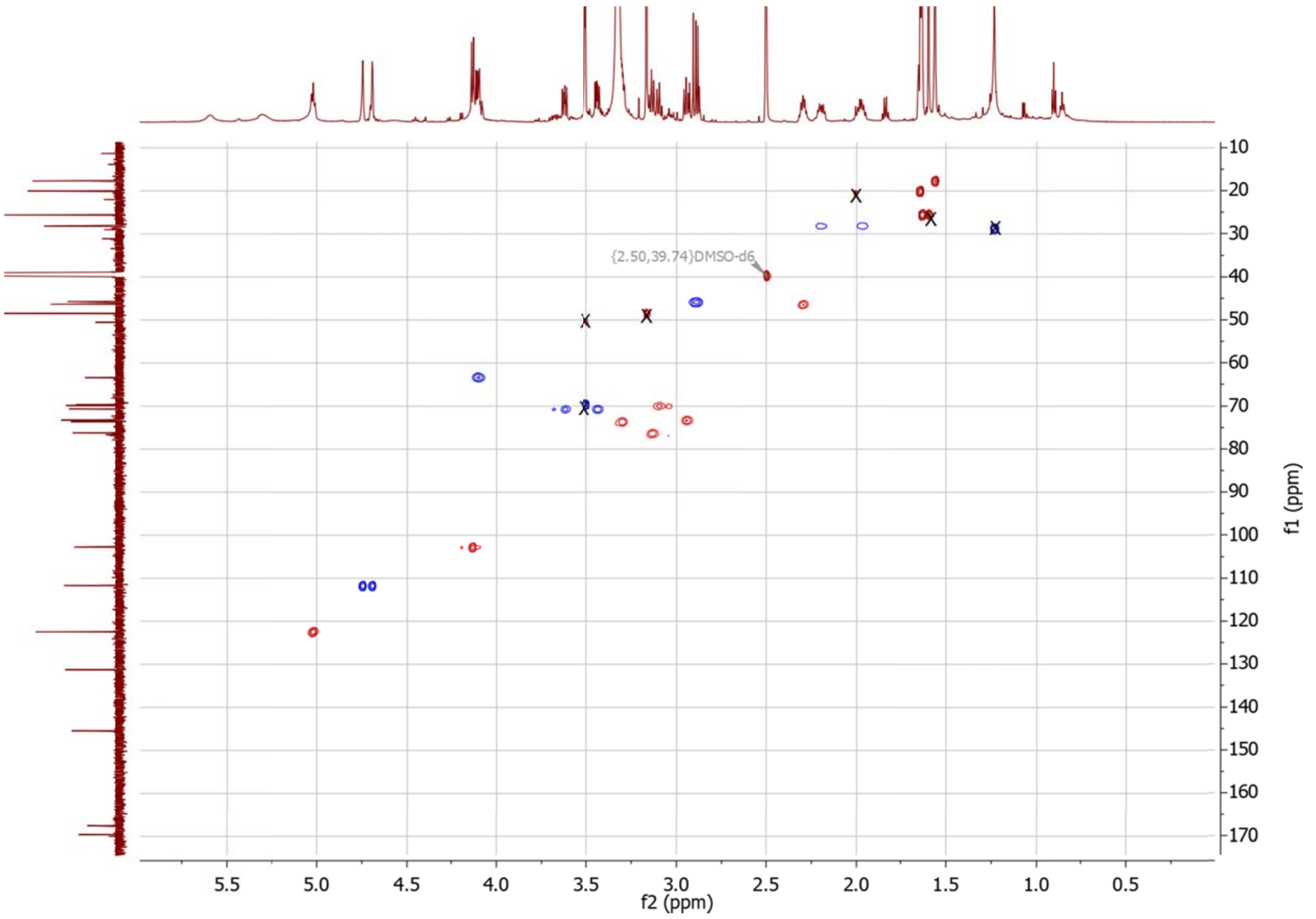

**Figure S2.20** $^{1}H$ -$^{13}C$ HSQC spectrum of **3** in DMSO-$d_6$. The signals representing residual sample impurities are crossed out.

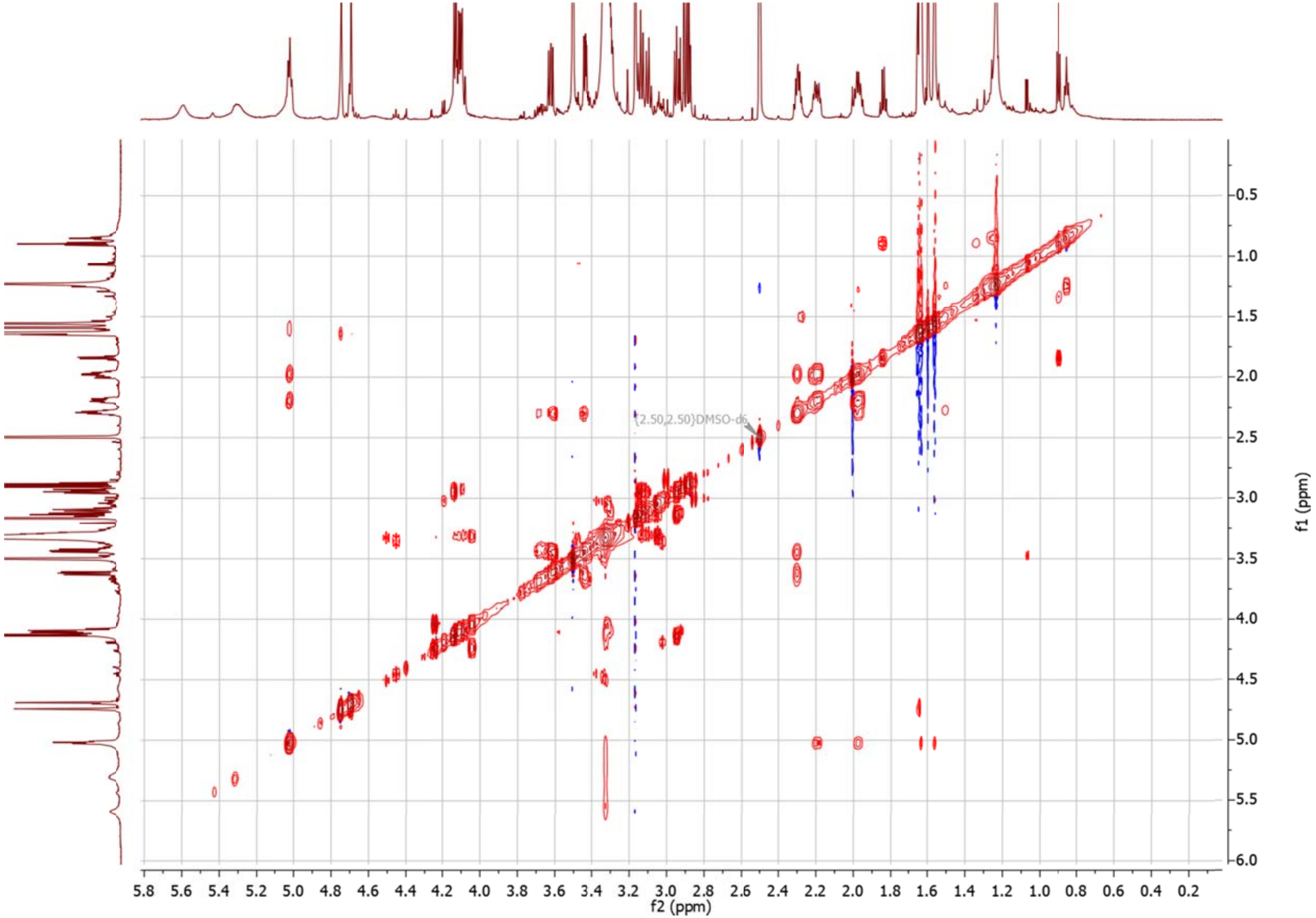

**Figure S2.21** $^{1}H$ -$^{1}H$ CLIP-COSY spectrum of **3** in DMSO-$d_6$.

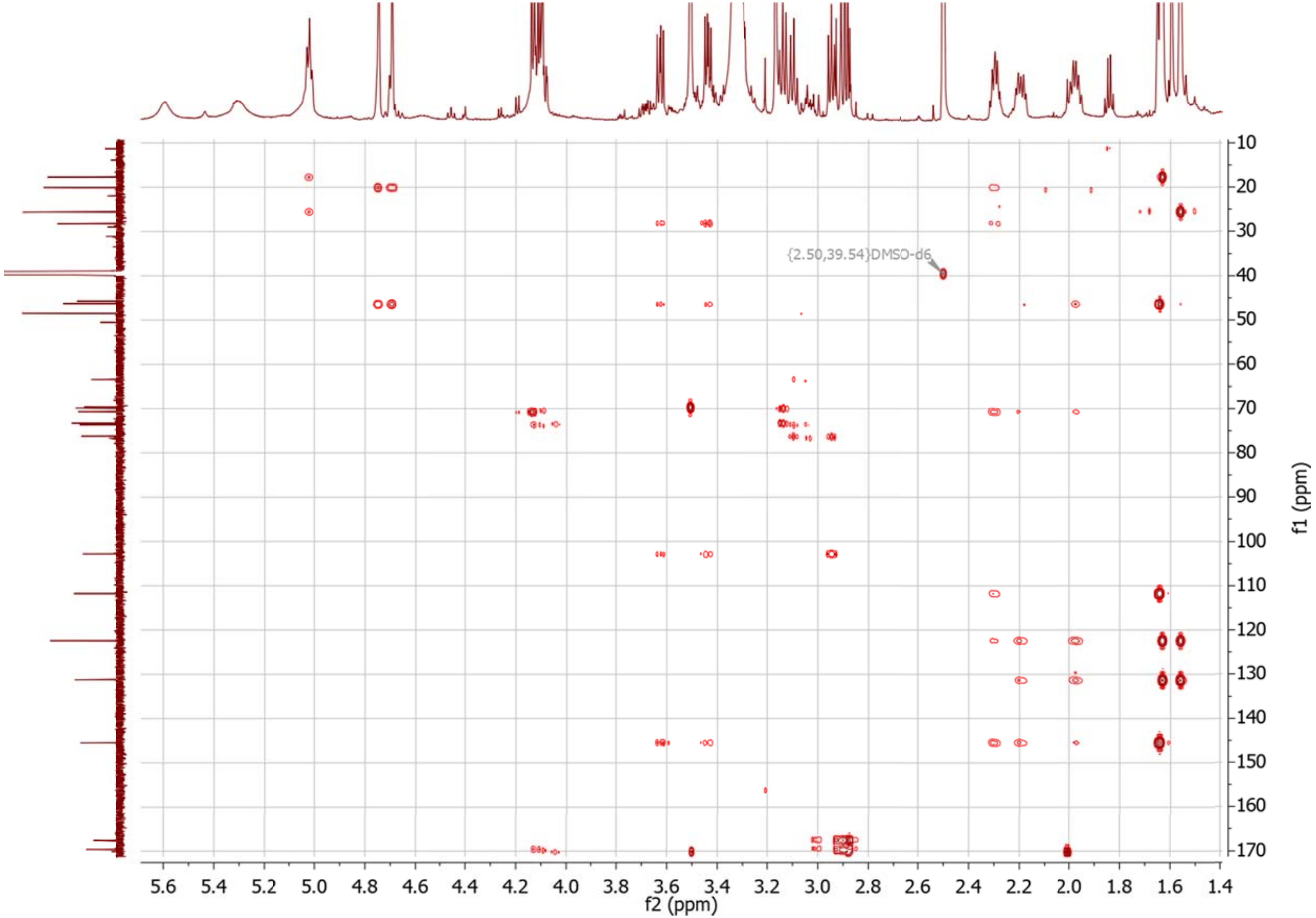

**Figure S2.22** $^{1}H$ -$^{13}C$ HMBC spectrum of **3** in DMSO-$d_6$.

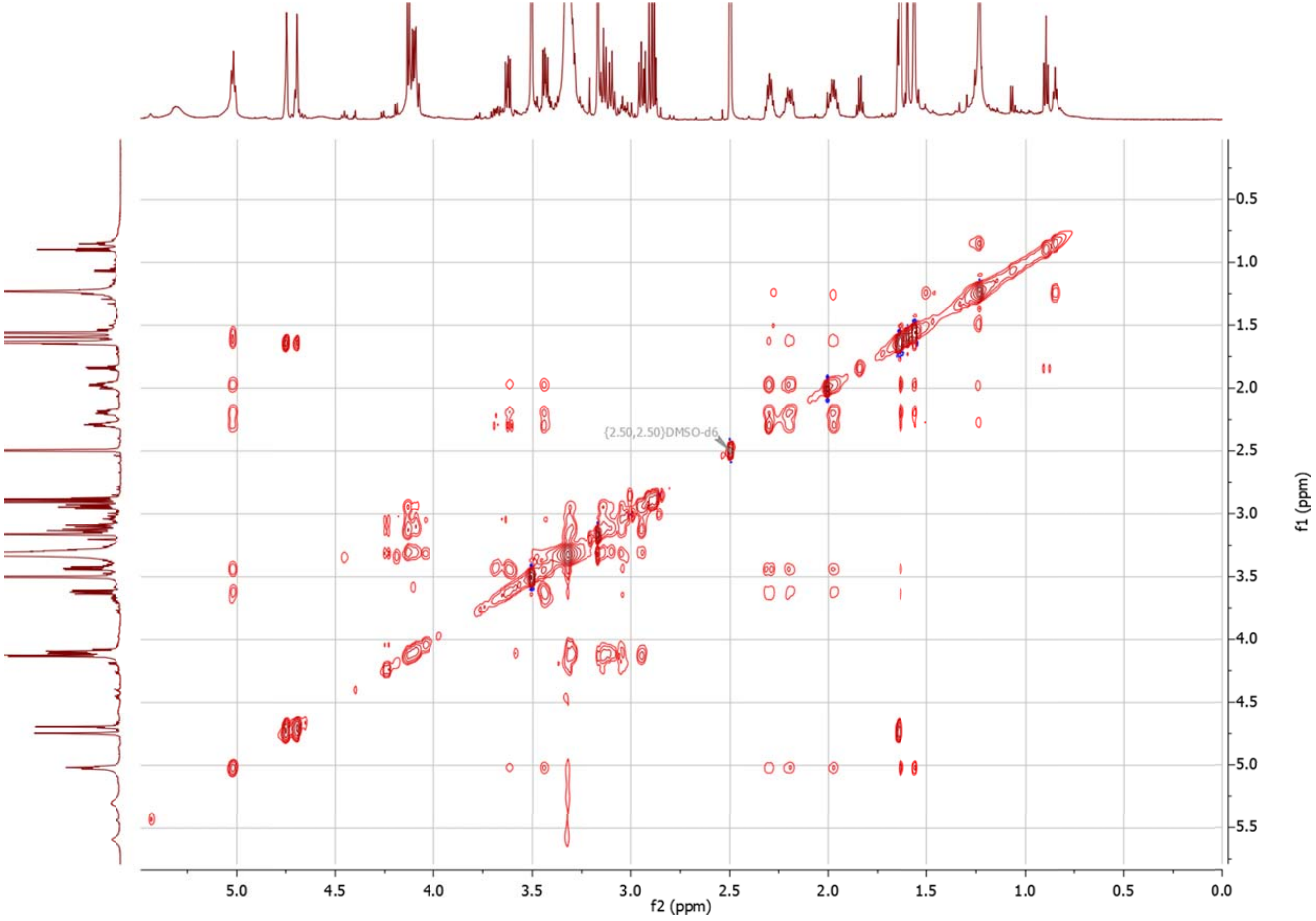

**Figure S2.23** $^{1}H$ -$^{1}H$ TOCSY spectrum of **3** in DMSO-$d_6$.

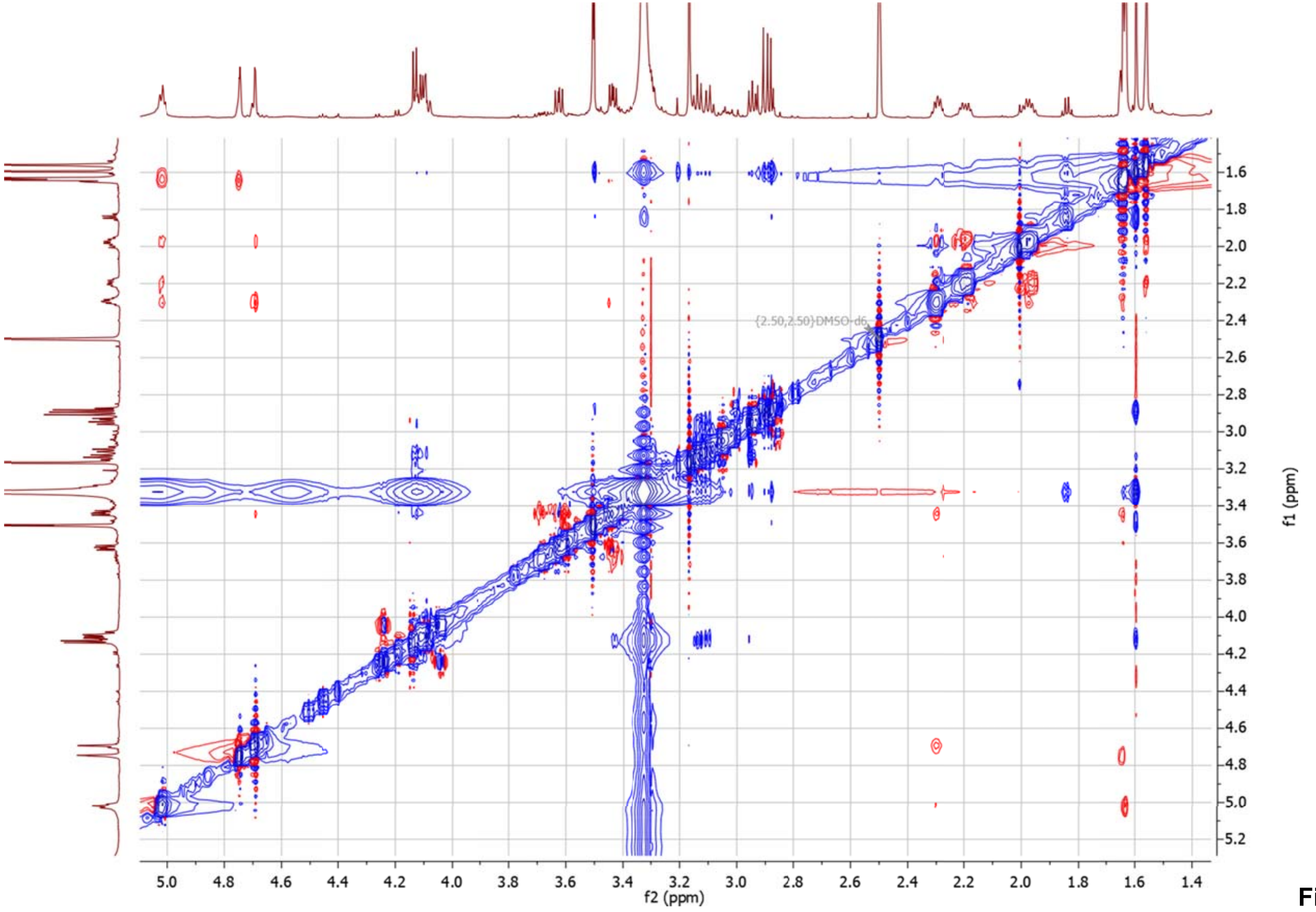

**Figure S2.24** 1H -1H NOESY spectrum of **3** in DMSO-d6.

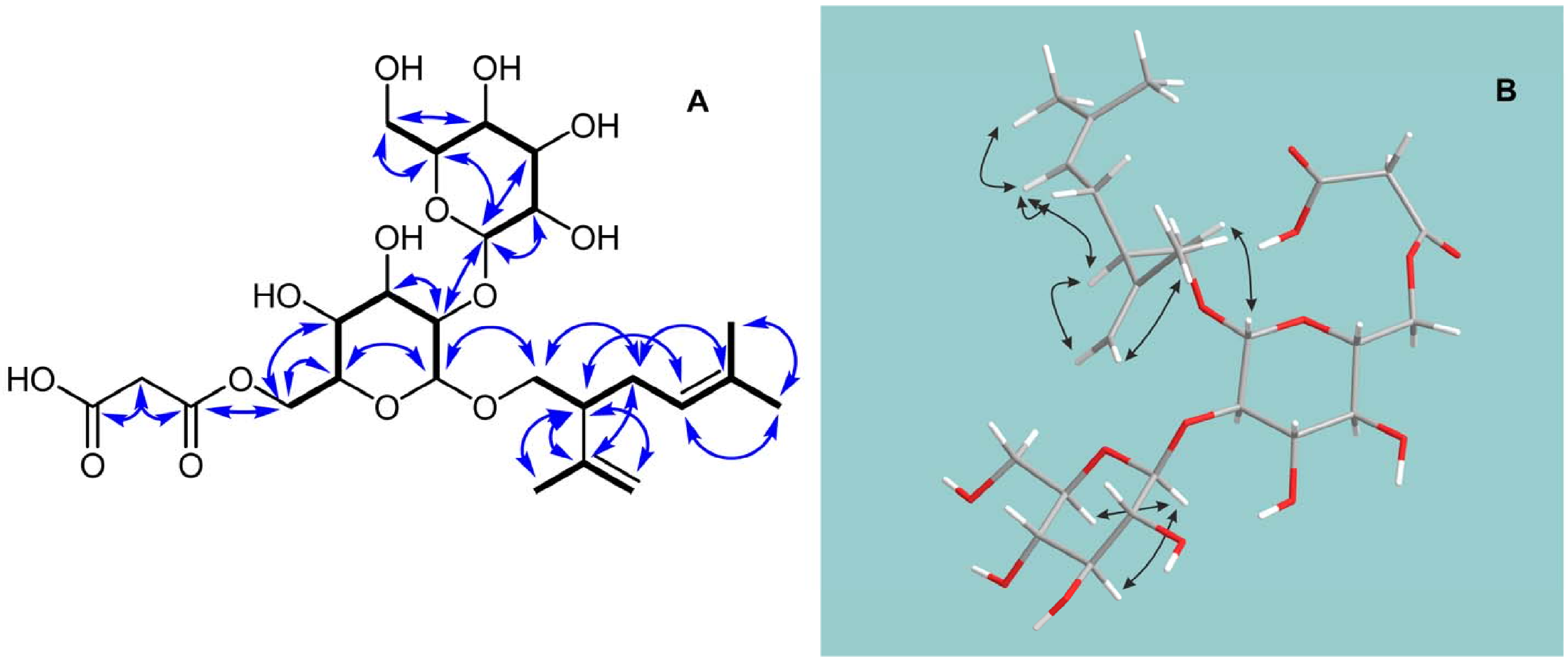

**Figure S2.25** COSY (bold lines) and key HMBC (arrows) correlations (A), and NOESY correlations (B) of **4** in DMSO-$d_6$.

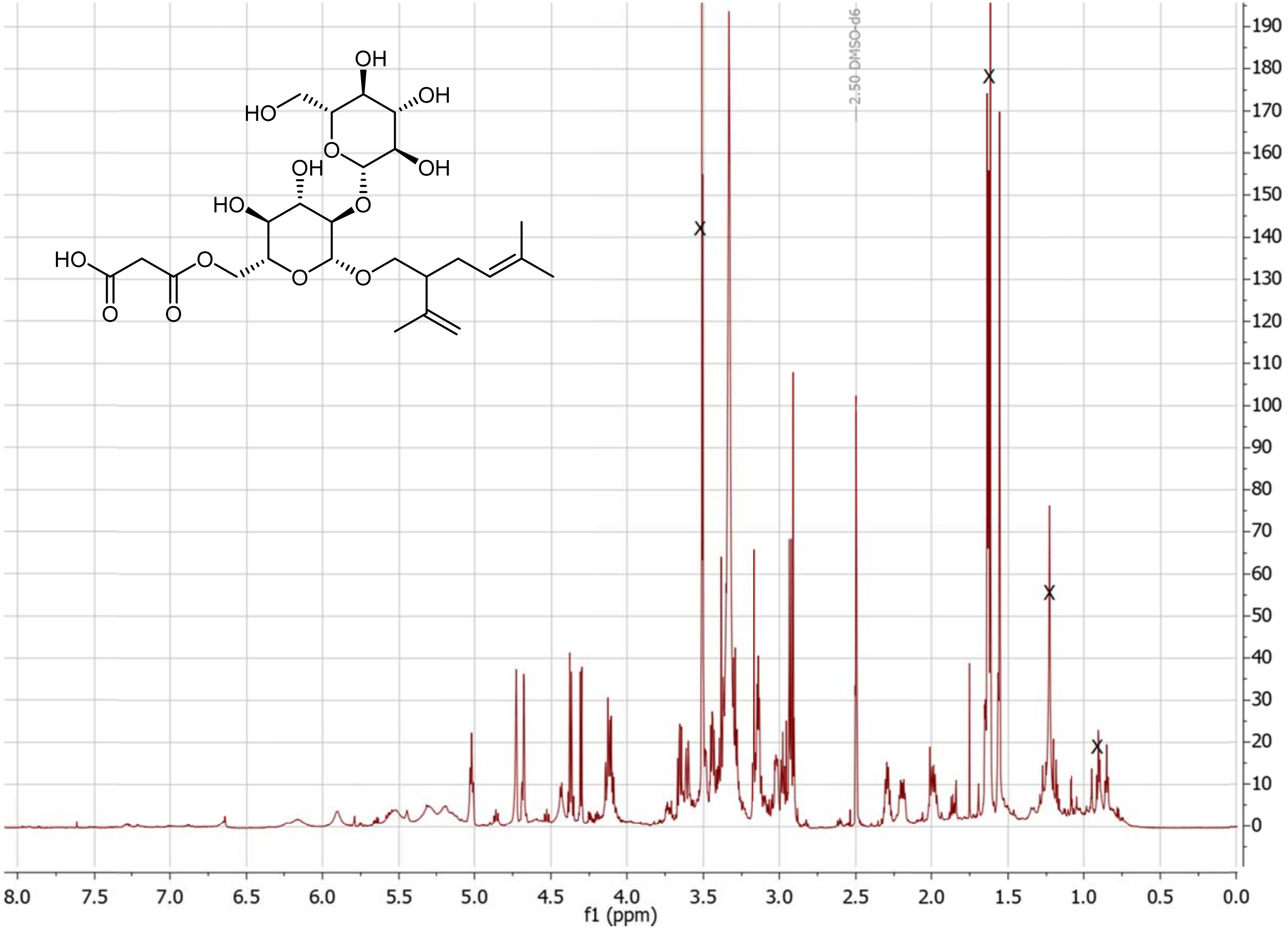

**Figure S2.26** $^1$H NMR spectrum of **4** in DMSO-$d_6$. The signals representing residual sample impurities are crossed out.

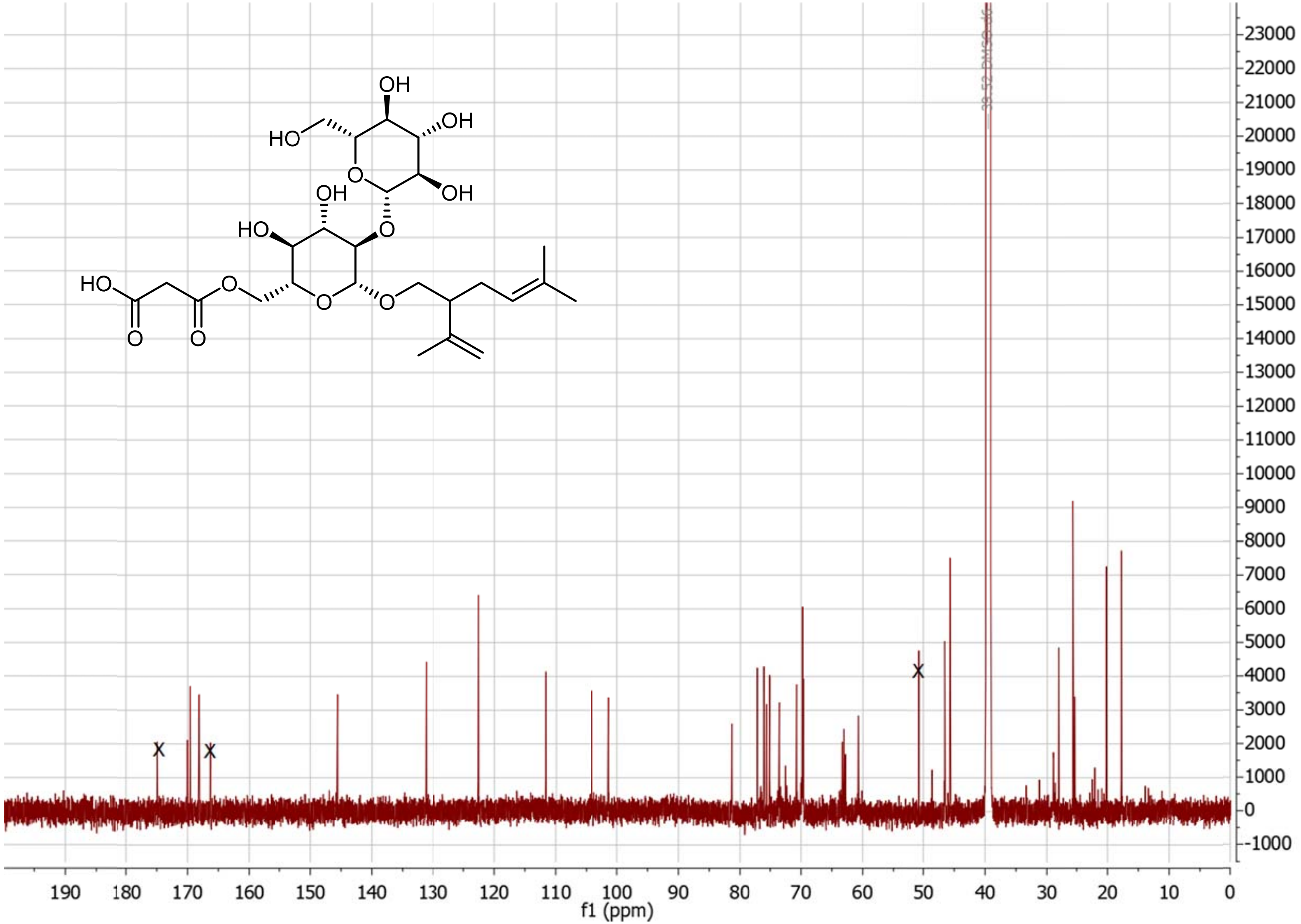

**Figure S2.27** $^{13}C$ NMR spectrum of **4** in DMSO-$d_6$. The signals representing residual sample impurities are crossed out.

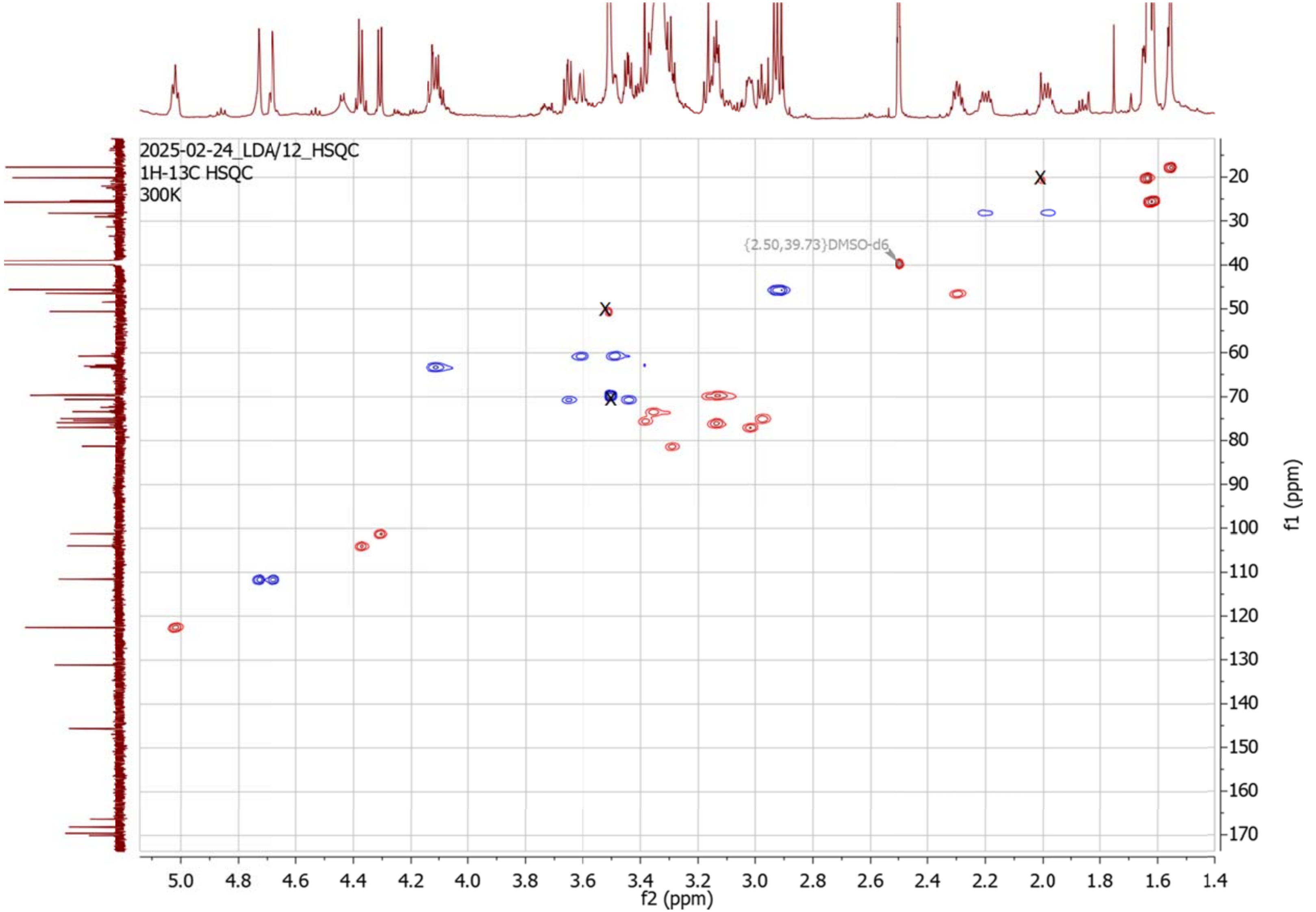

**Figure S2.28** $^{1}H$ -$^{13}C$ HSQC spectrum of **4** in DMSO-$d_6$. The signals representing residual sample impurities are crossed out.

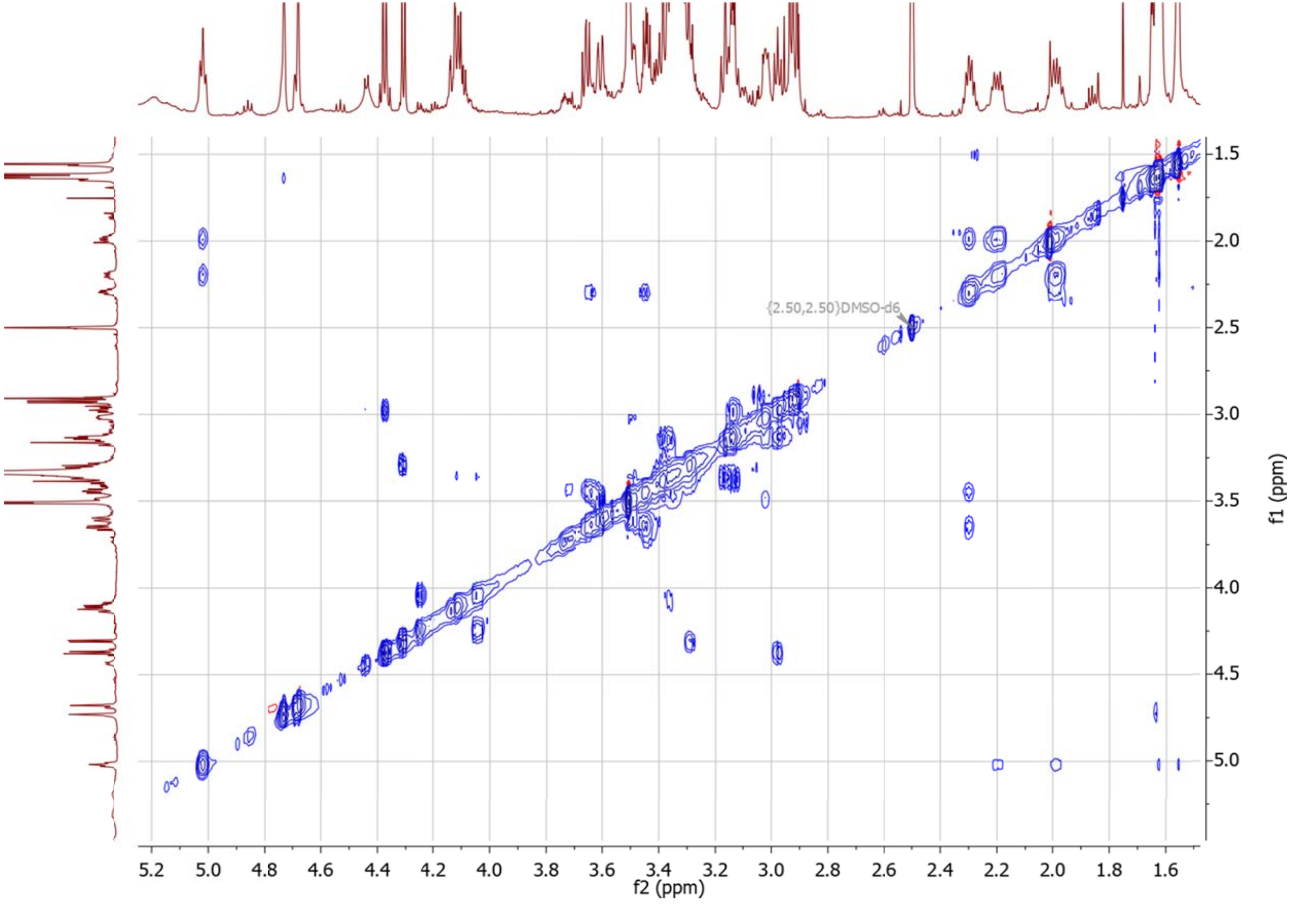

**Figure S2.29** [1]H -[1]H CLIP-COSY spectrum of **4** in DMSO-$d_6$.

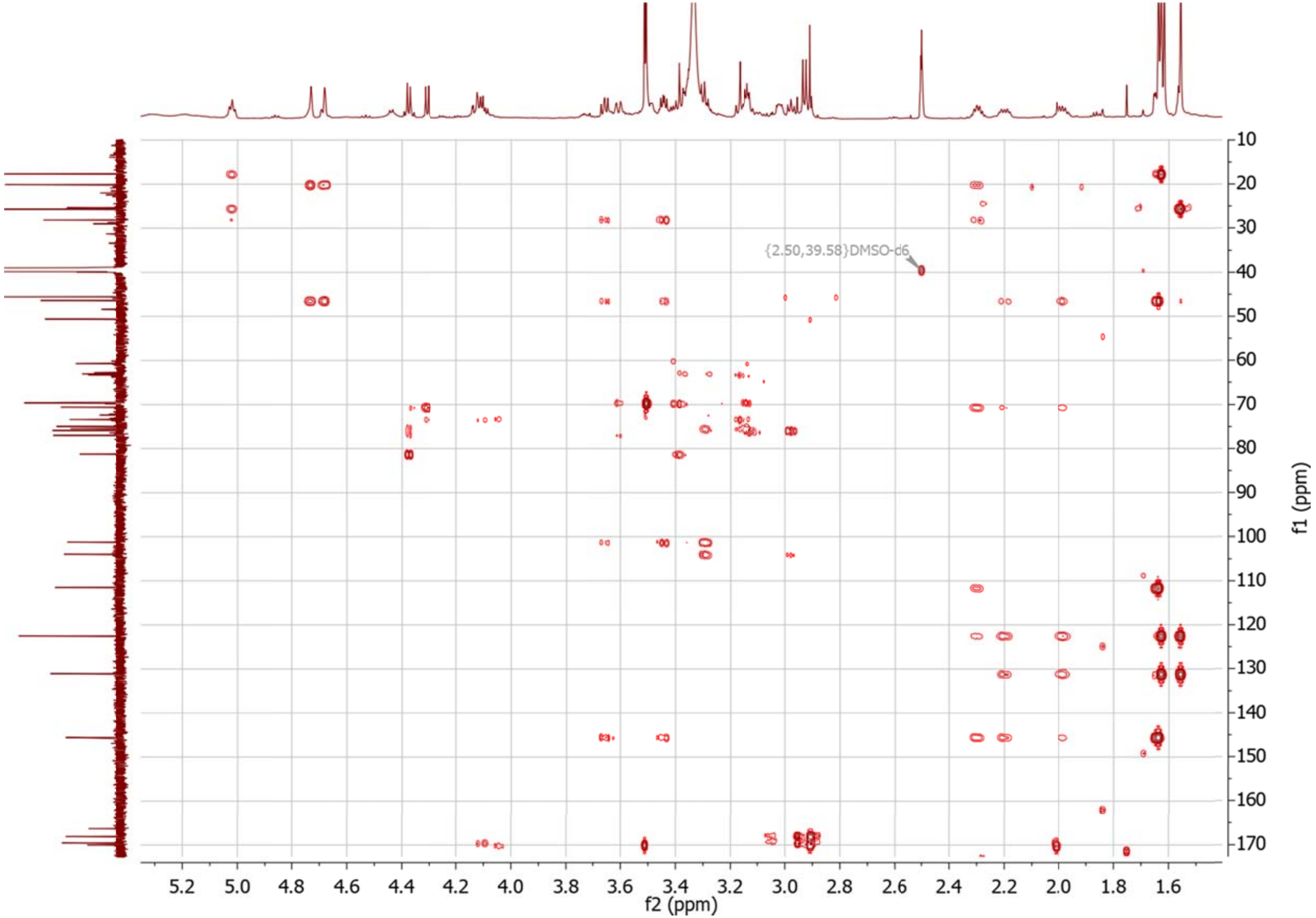

**Figure S2.30** $^{1}H$ -$^{13}C$ HMBC spectrum of **4** in DMSO-$d_6$.

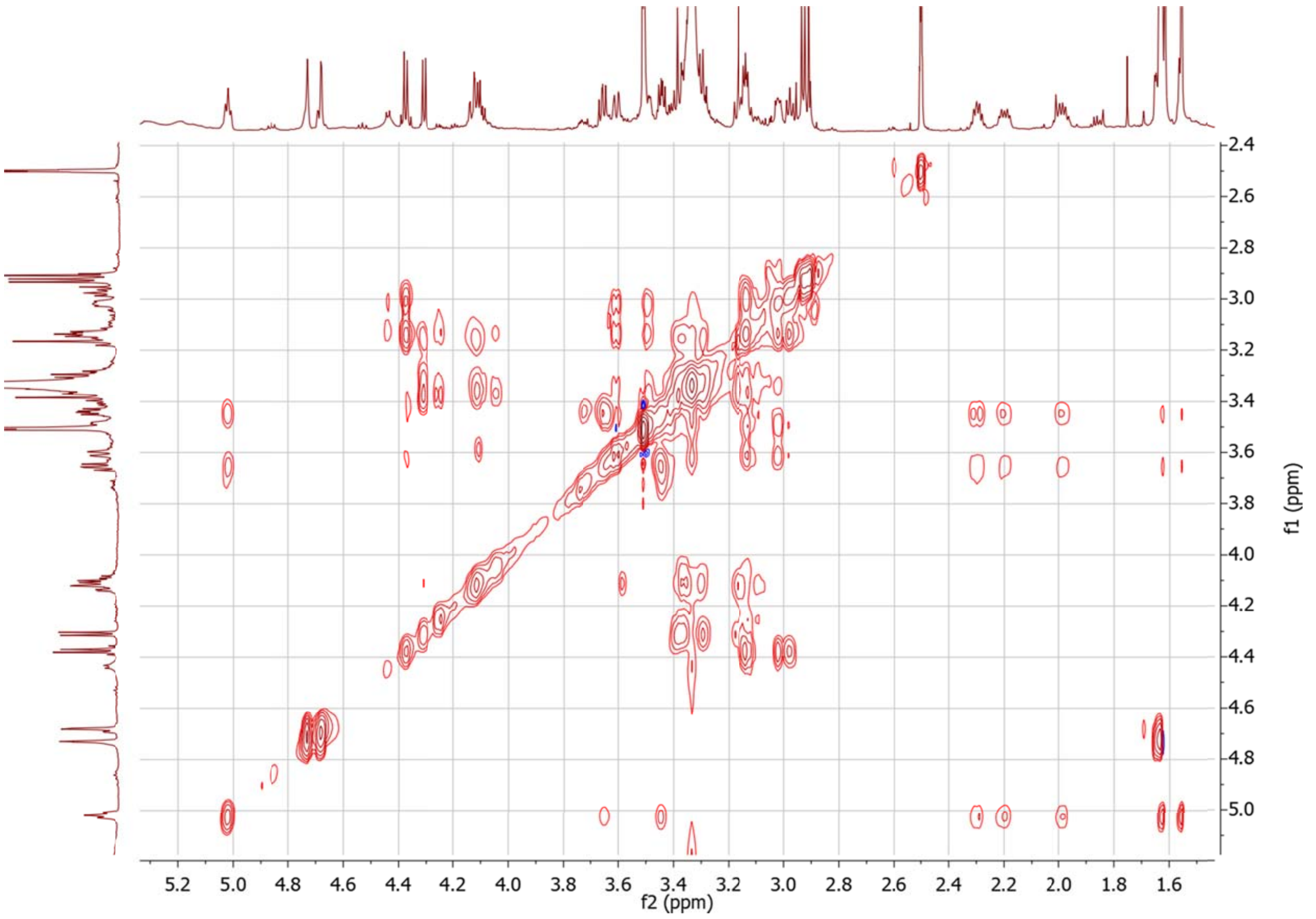

**Figure S2.31** [1]H -[1]H TOCSY spectrum of **4** in DMSO-$d_6$.

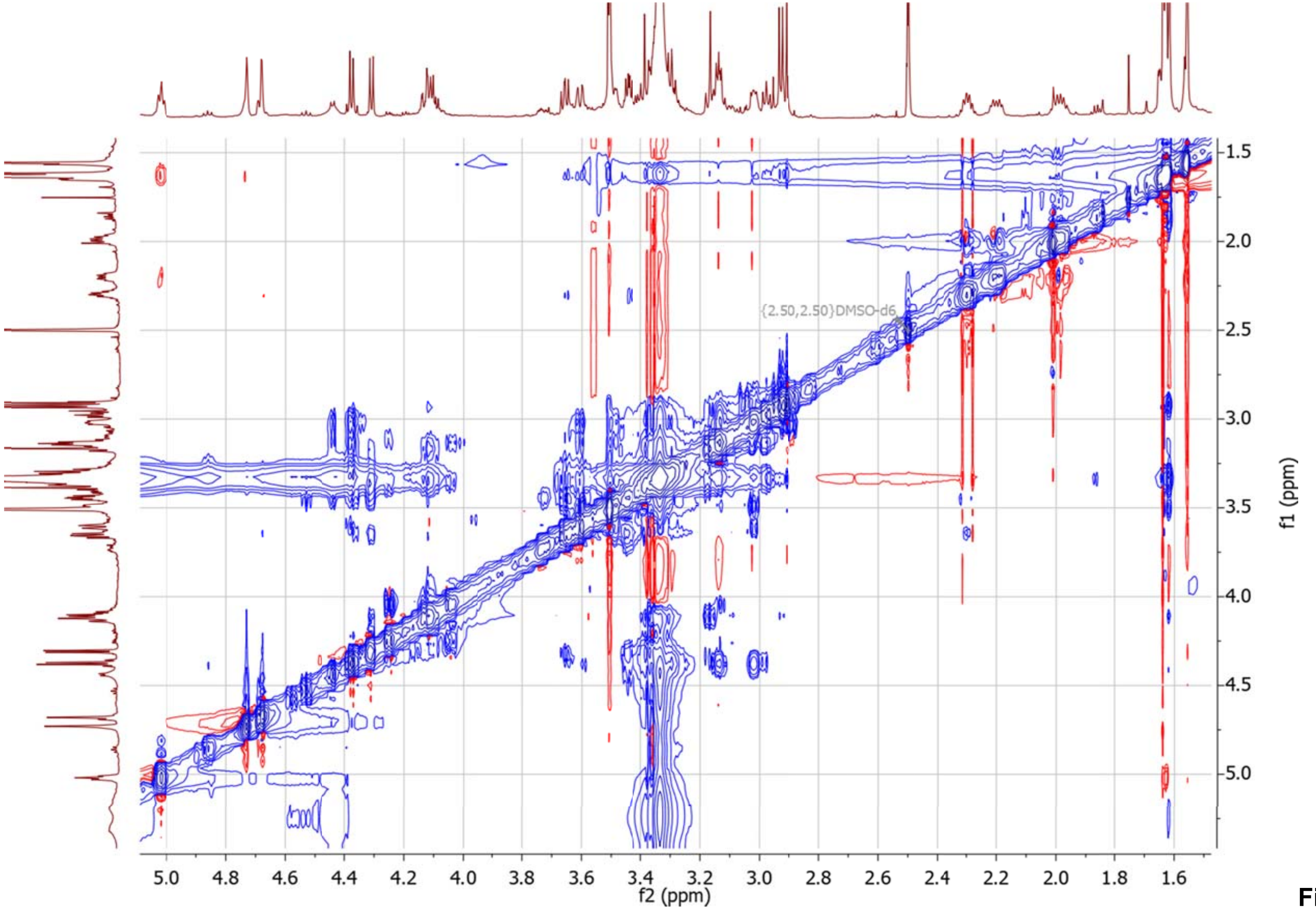

**Figure S2.32** $^1$H -$^1$H NOESY spectrum of **4** in DMSO-$d_6$.

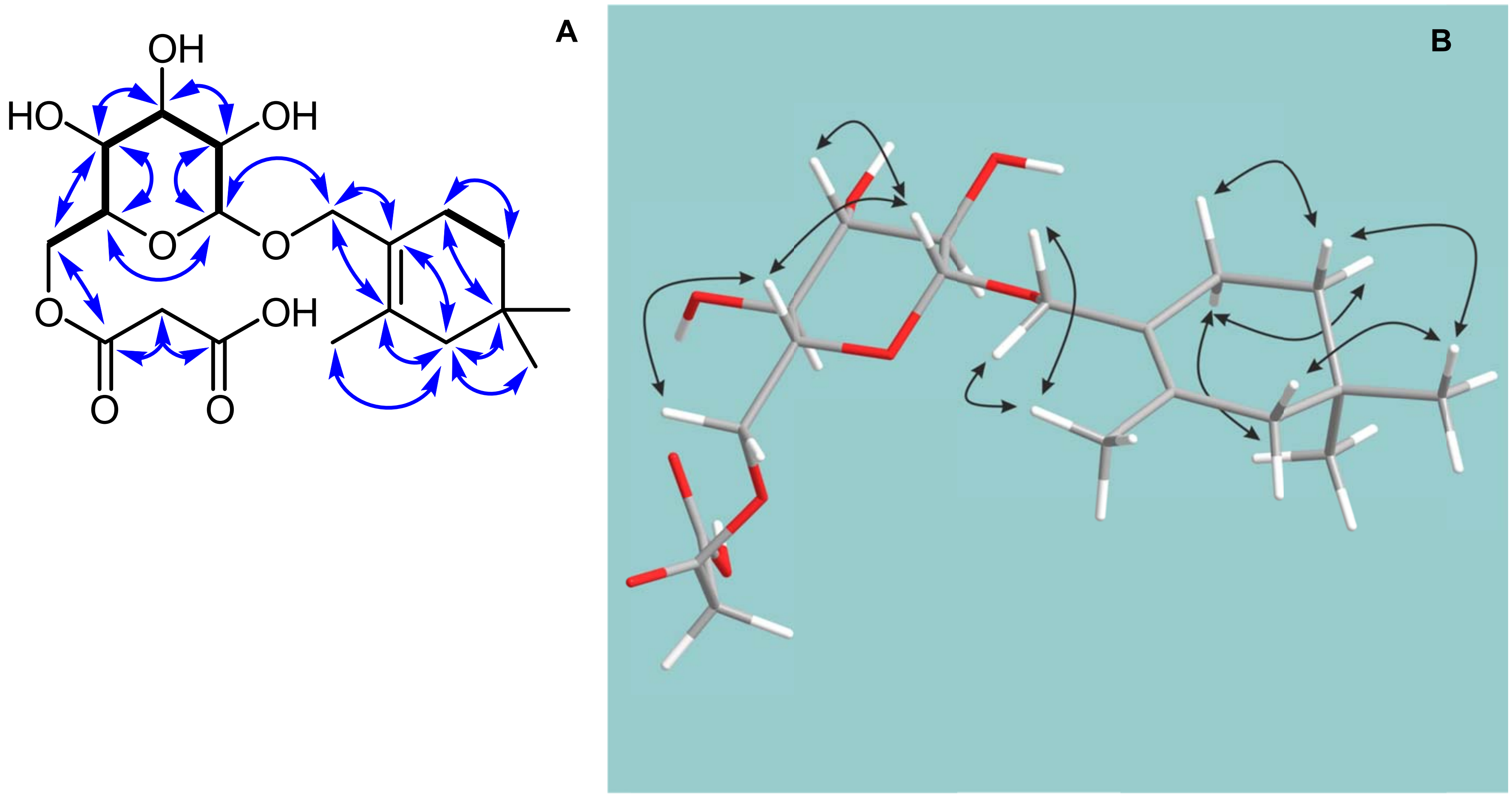

**Figure S2.33** COSY (bold lines) and key HMBC (arrows) correlations (A), and NOESY correlations (B) of **5** in DMSO-$d_6$.

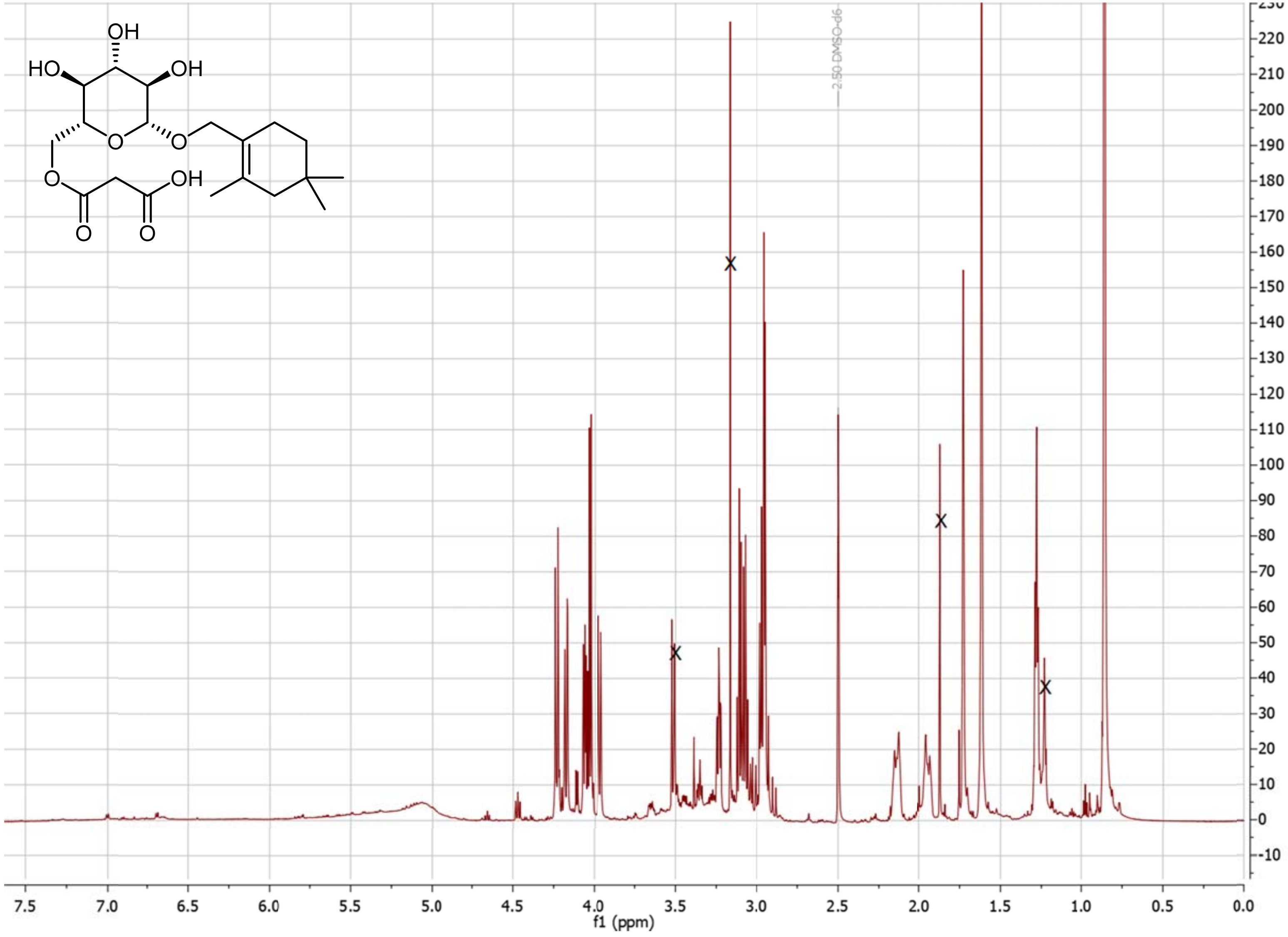

**Figure S2.34** $^{1}$H NMR spectrum of **5** in DMSO-$d_6$. The signals representing residual sample impurities are crossed out.

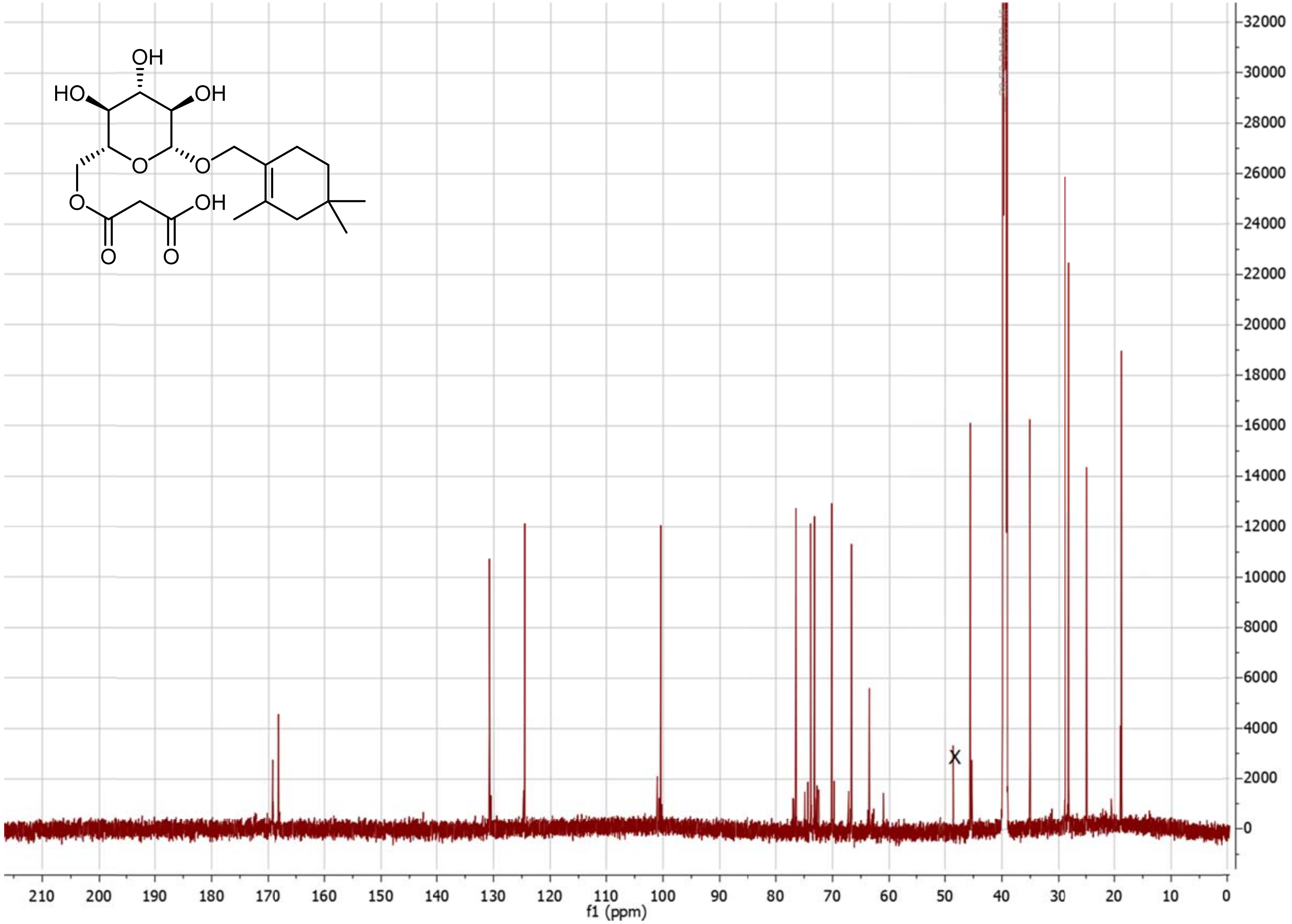

**Figure S2.35** $^{13}$C NMR spectrum of **5** in DMSO-$d_6$. The signals representing residual sample impurities are crossed out.

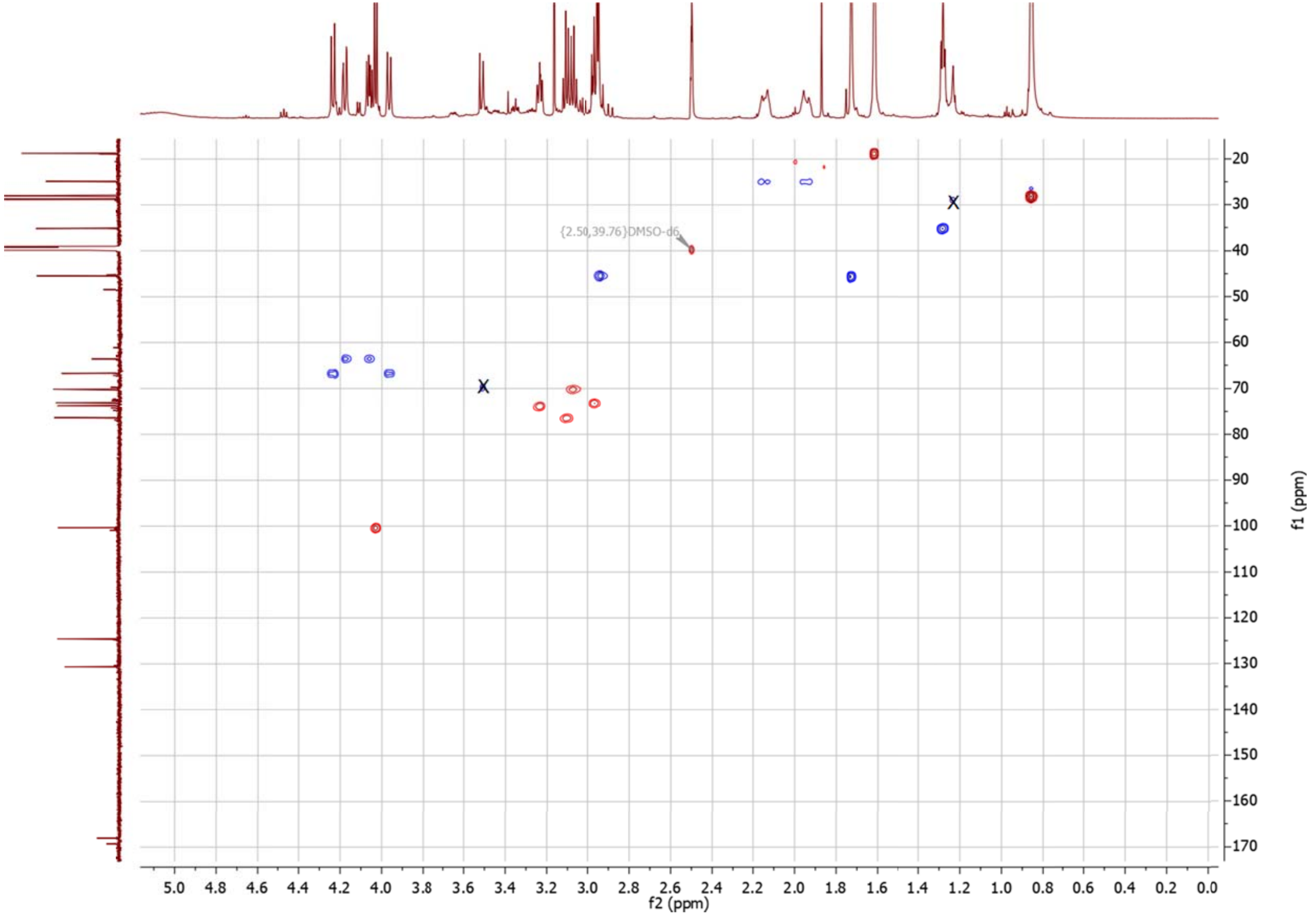

**Figure S2.36** $^{1}H$ -$^{13}C$ HSQC spectrum of **5** in DMSO-$d_6$. The signals representing residual sample impurities are crossed out.

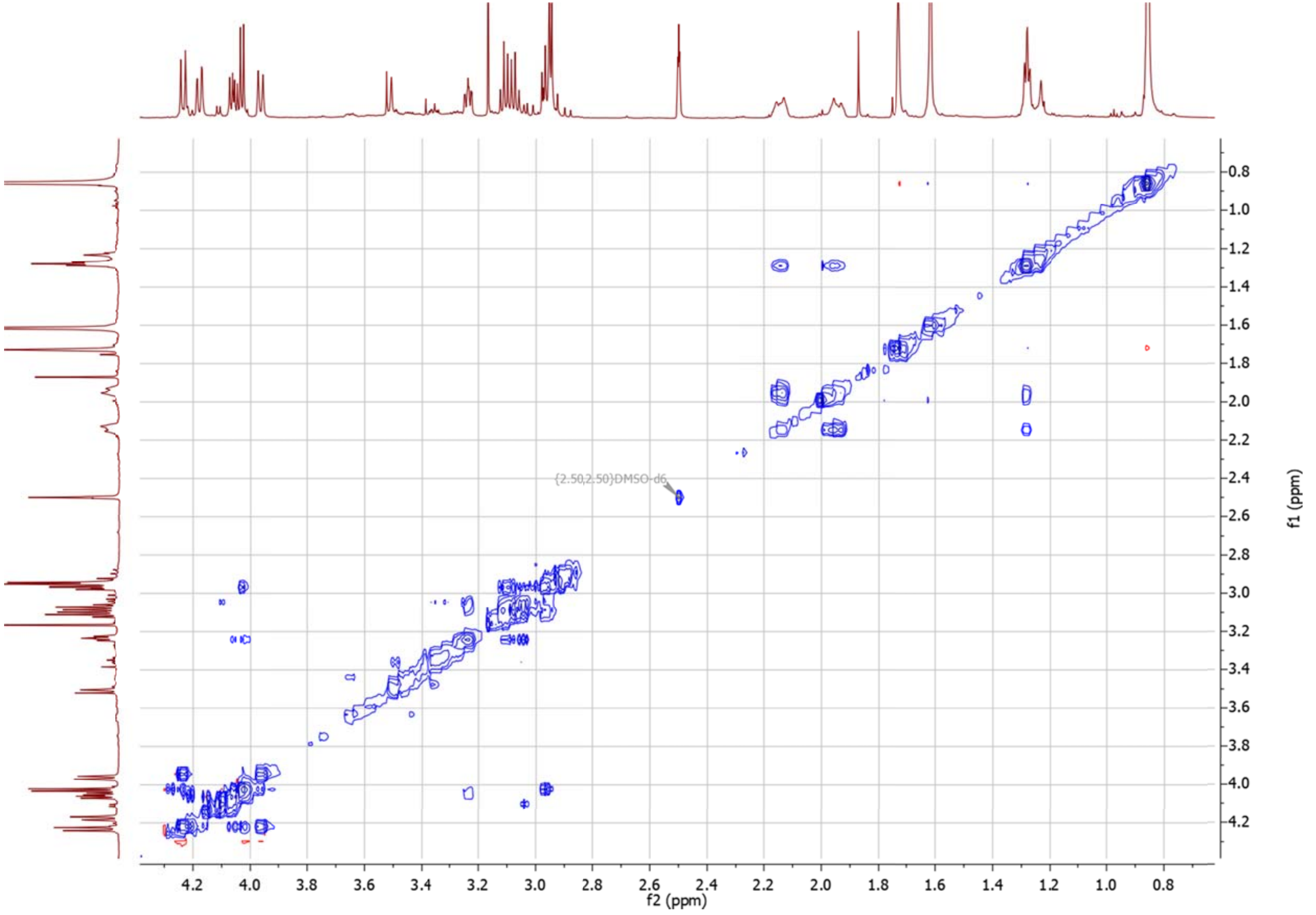

**Figure S2.37** $^{1}H$ -$^{1}H$ CLIP-COSY spectrum of **5** in DMSO-$d_6$.

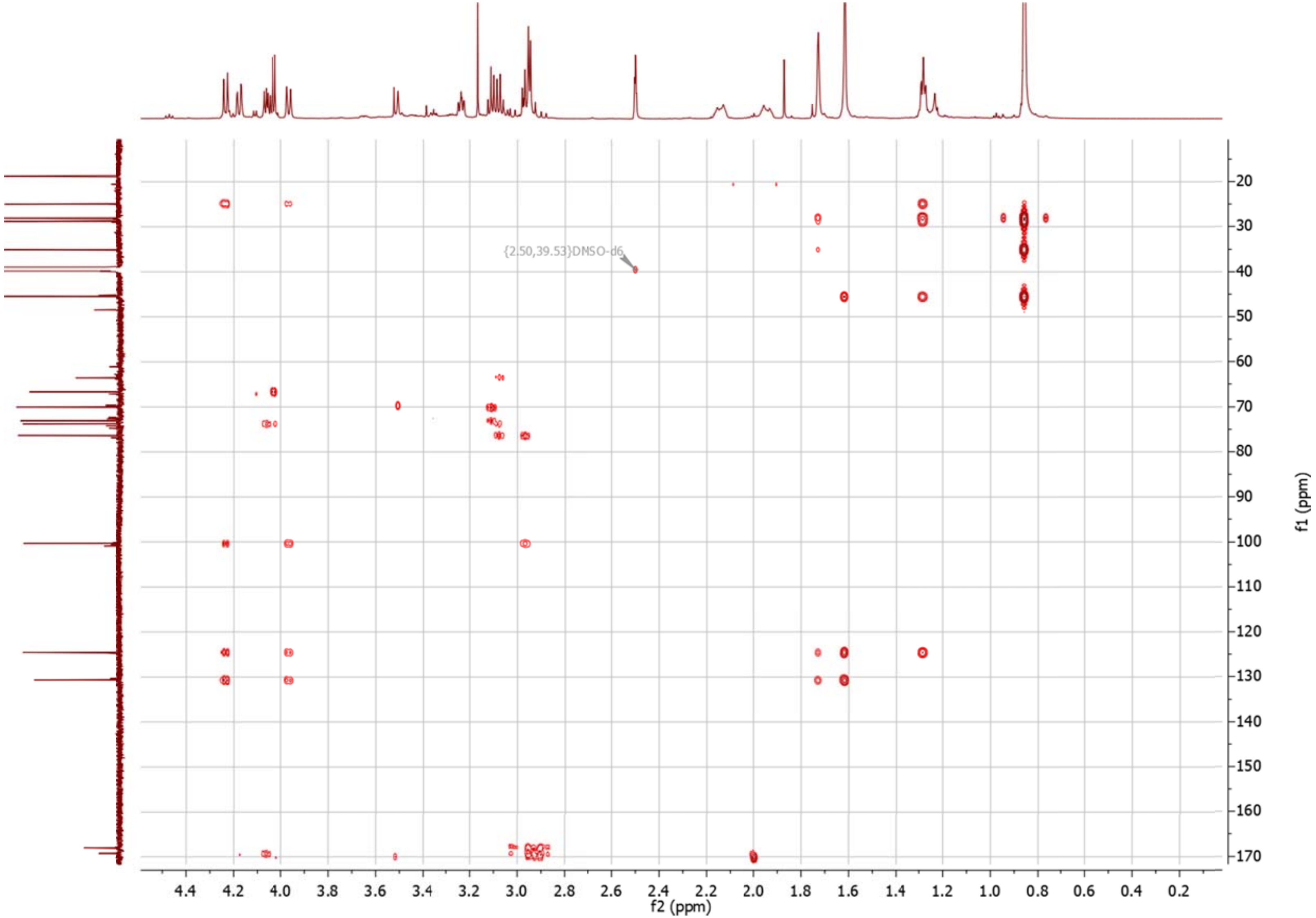

**Figure S2.38** $^{1}H$ -$^{13}C$ HMBC spectrum of **5** in DMSO-$d_6$.

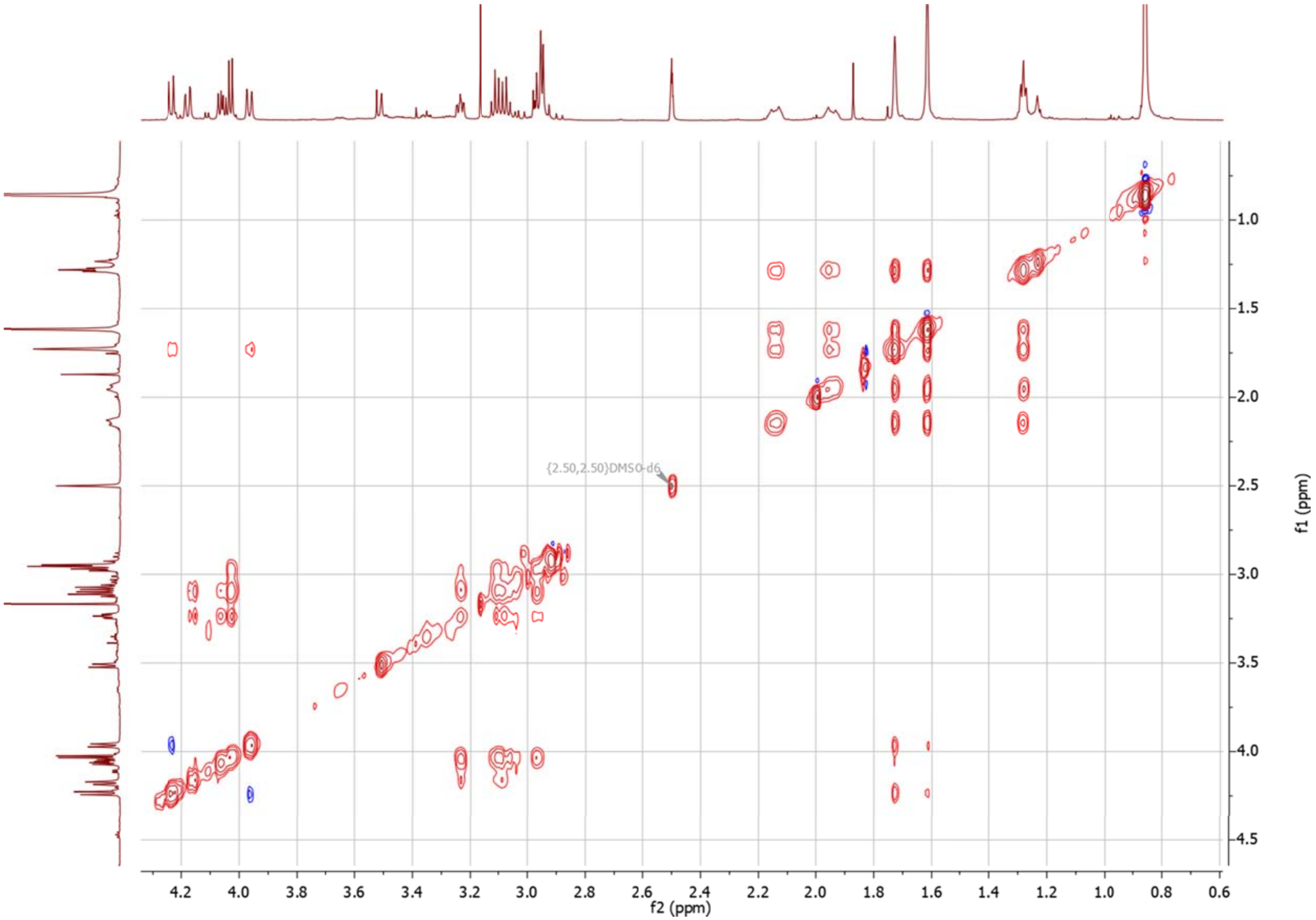

**Figure S2.39** [1]H -[1]H TOCSY spectrum of **5** in DMSO-$d_6$.

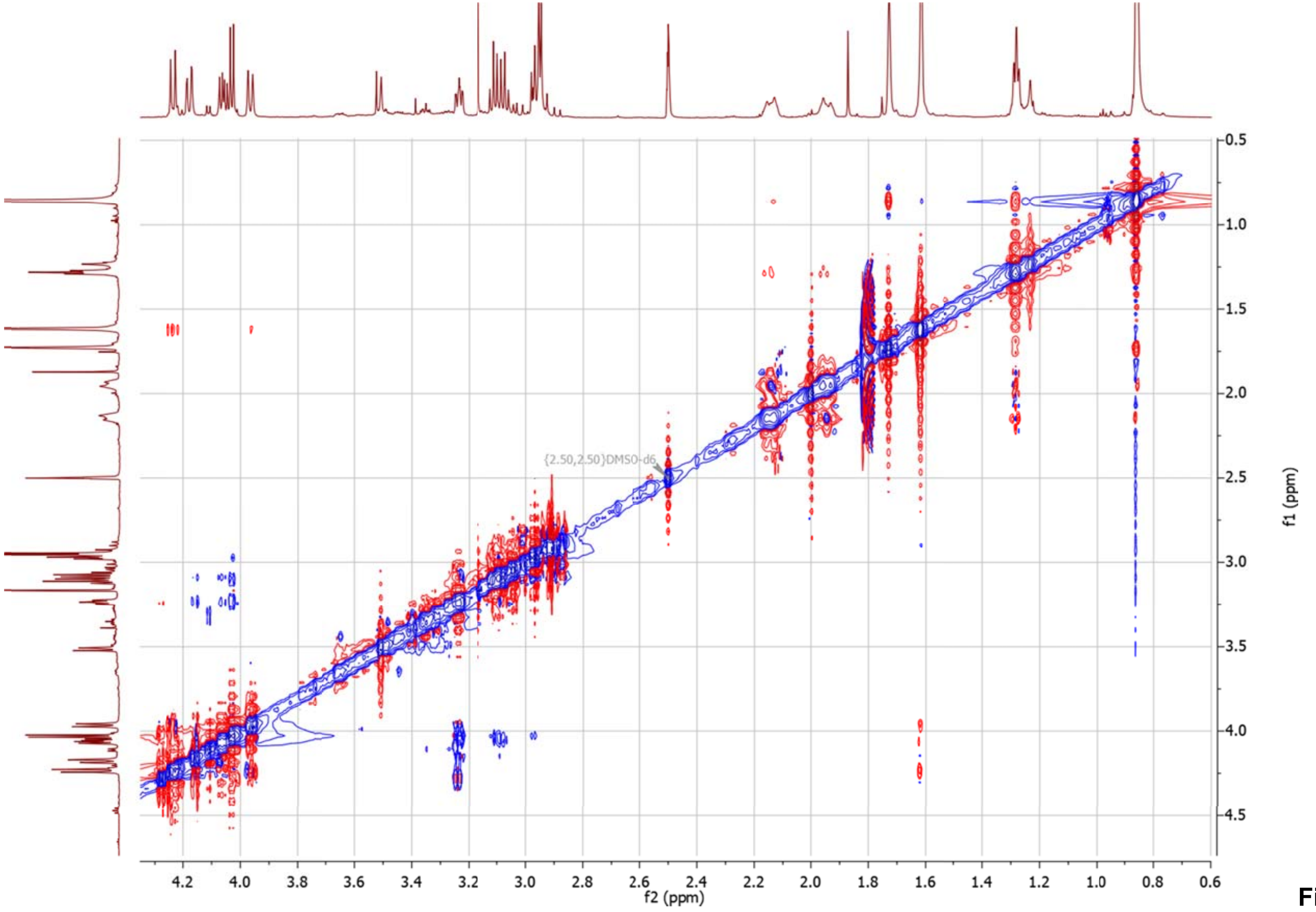

**Figure S2.40** $^{1}H$ -$^{1}H$ NOESY spectrum of **5** in DMSO-$d_6$.

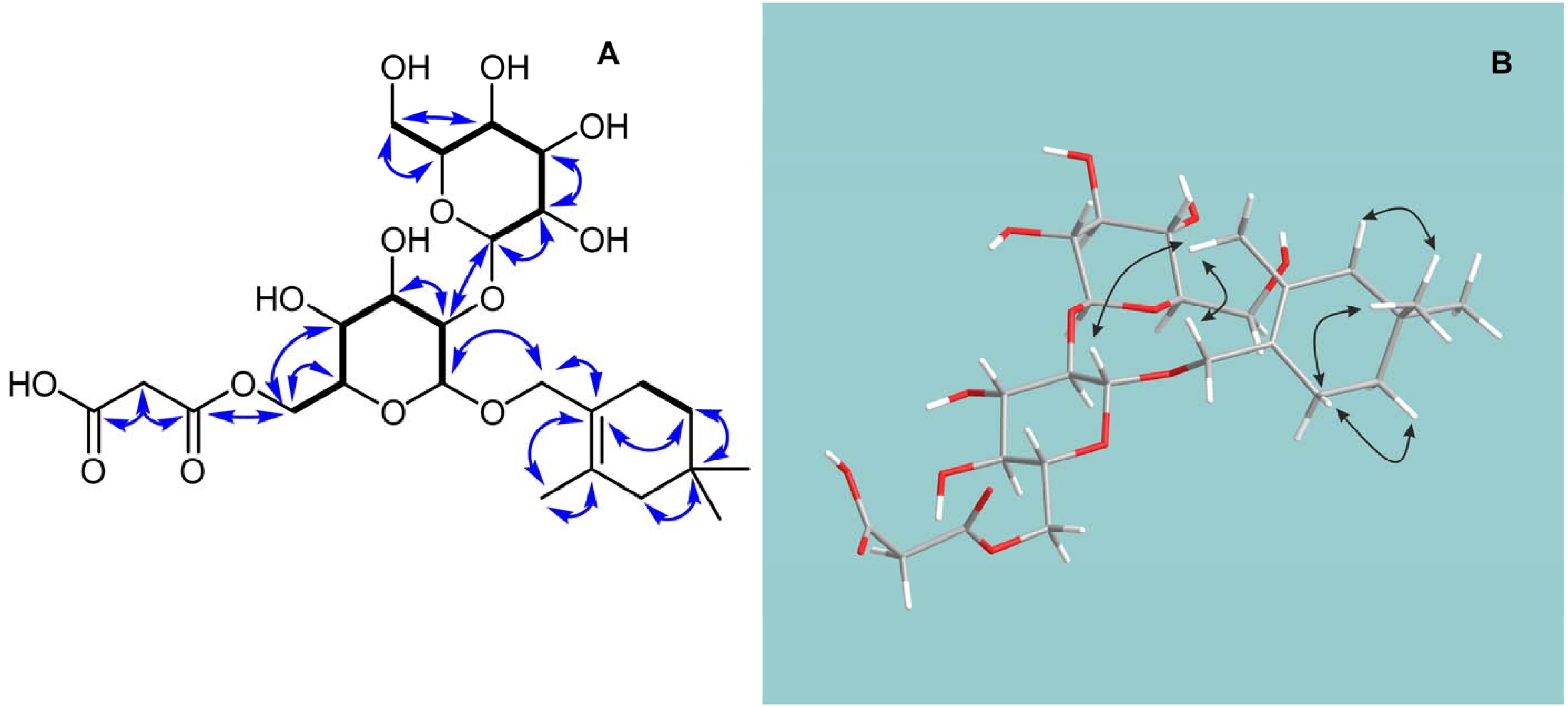

**Figure S2.41** COSY (bold lines) and key HMBC (arrows) correlations (A), and NOESY correlations (B) of **6** in DMSO-$d_6$.

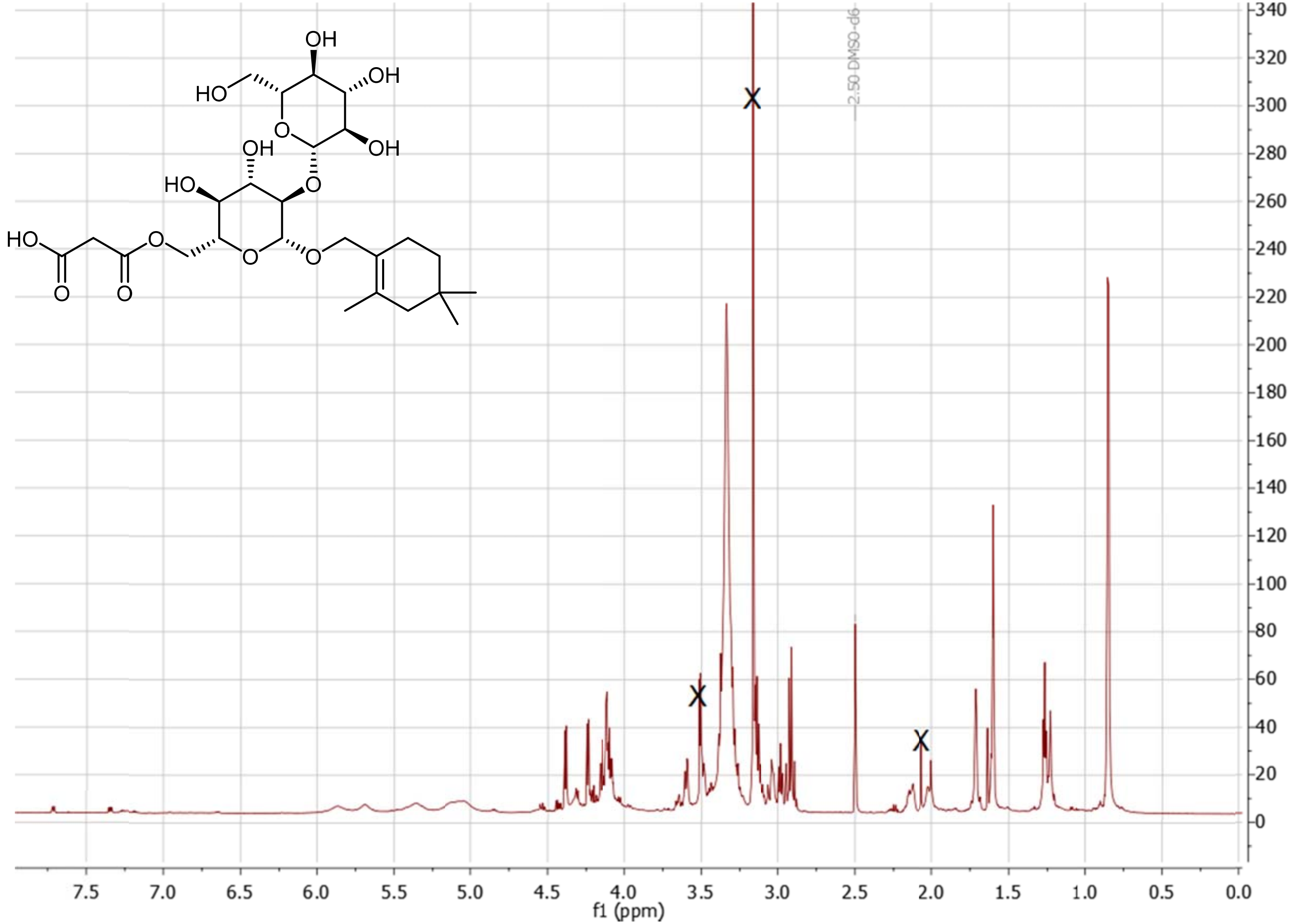

**Figure S2.42** $^1$H NMR spectrum of **6** in DMSO-$d_6$. The signals representing residual sample impurities are crossed out.

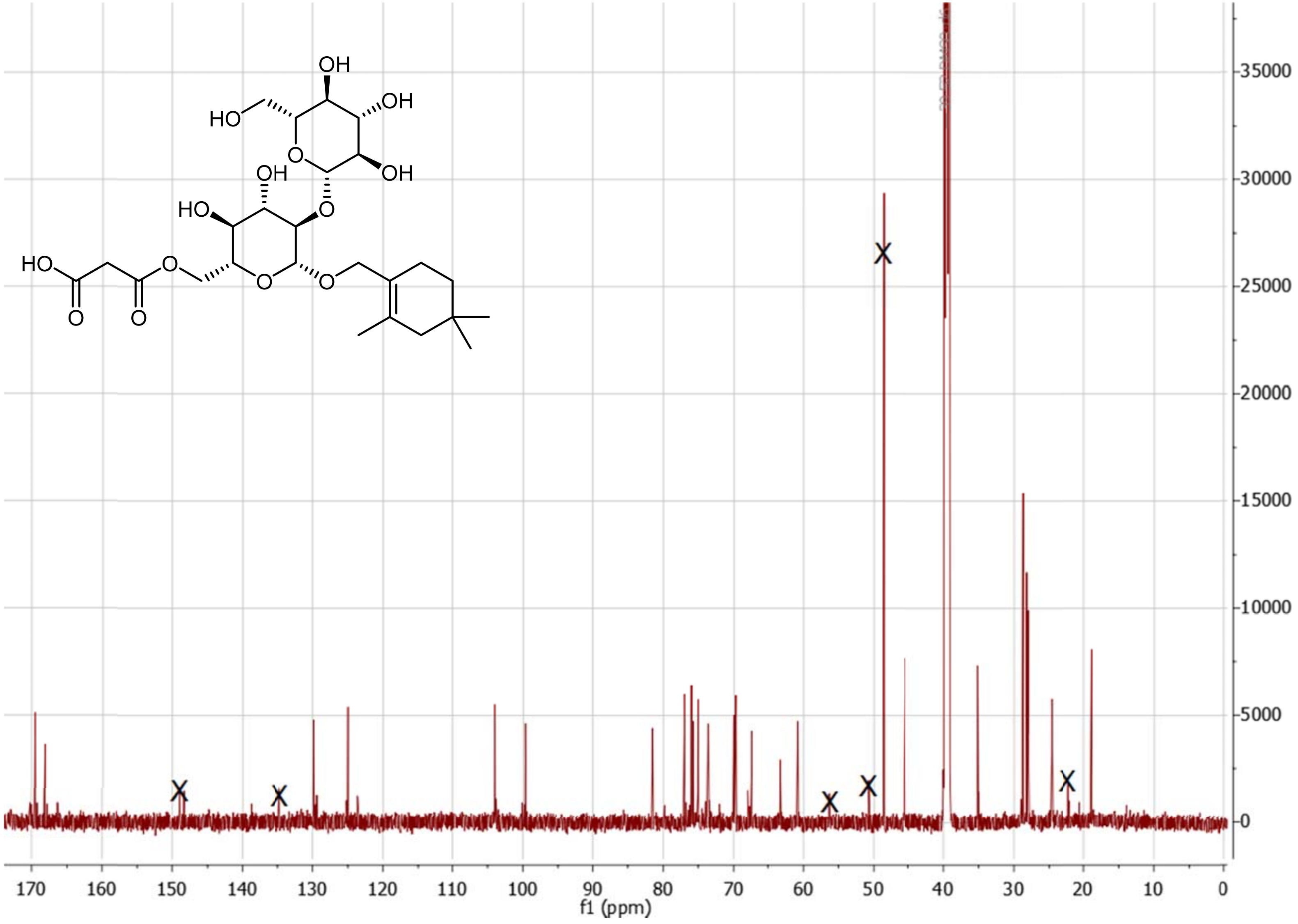

**Figure S2.43** $^{13}$C NMR spectrum of **6** in DMSO-$d_6$. The signals representing residual sample impurities are crossed out.

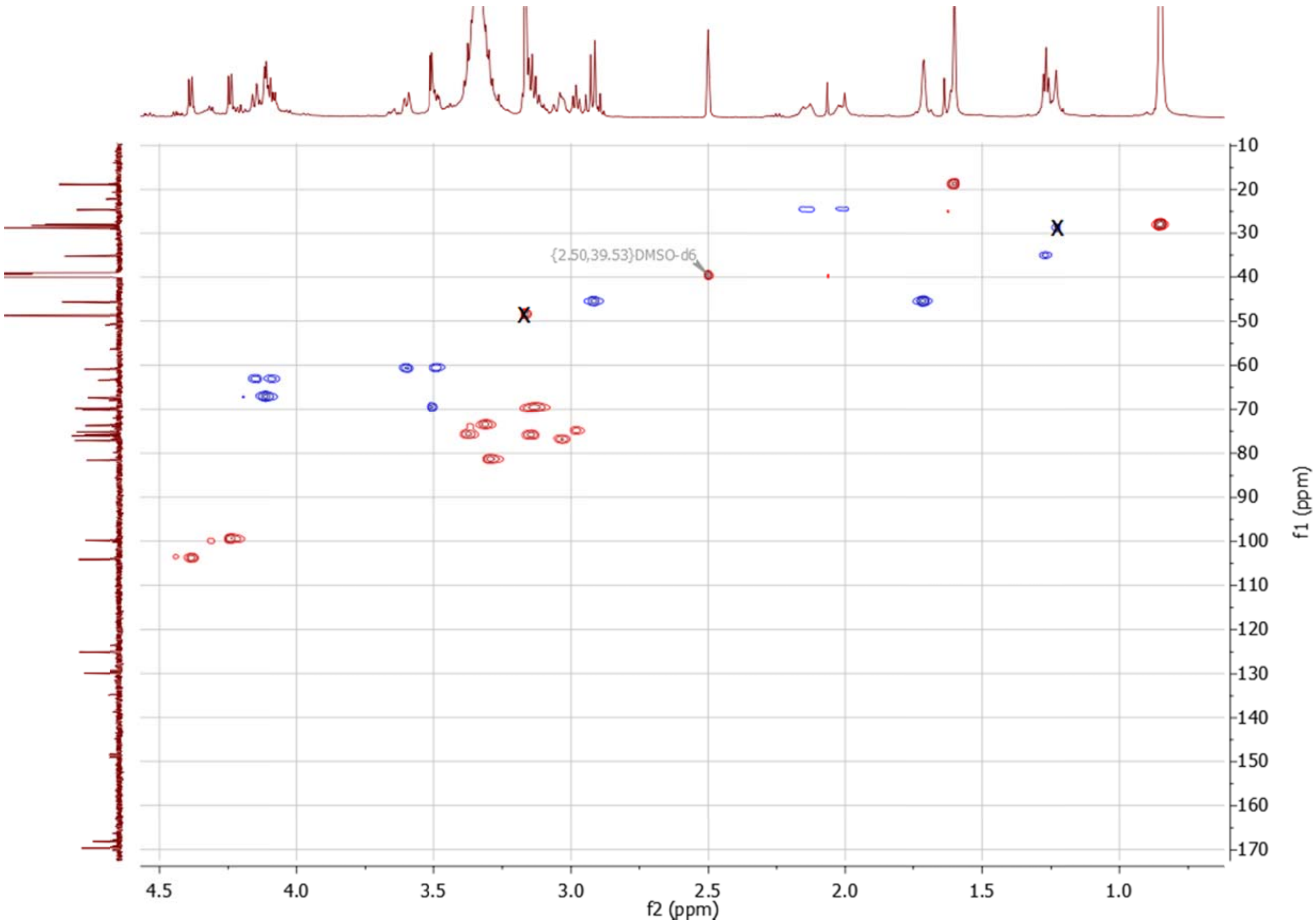

**Figure S2.44** $^{1}H$ -$^{13}C$ HSQC spectrum of **6** in DMSO-$d_6$. The signals representing residual sample impurities are crossed out.

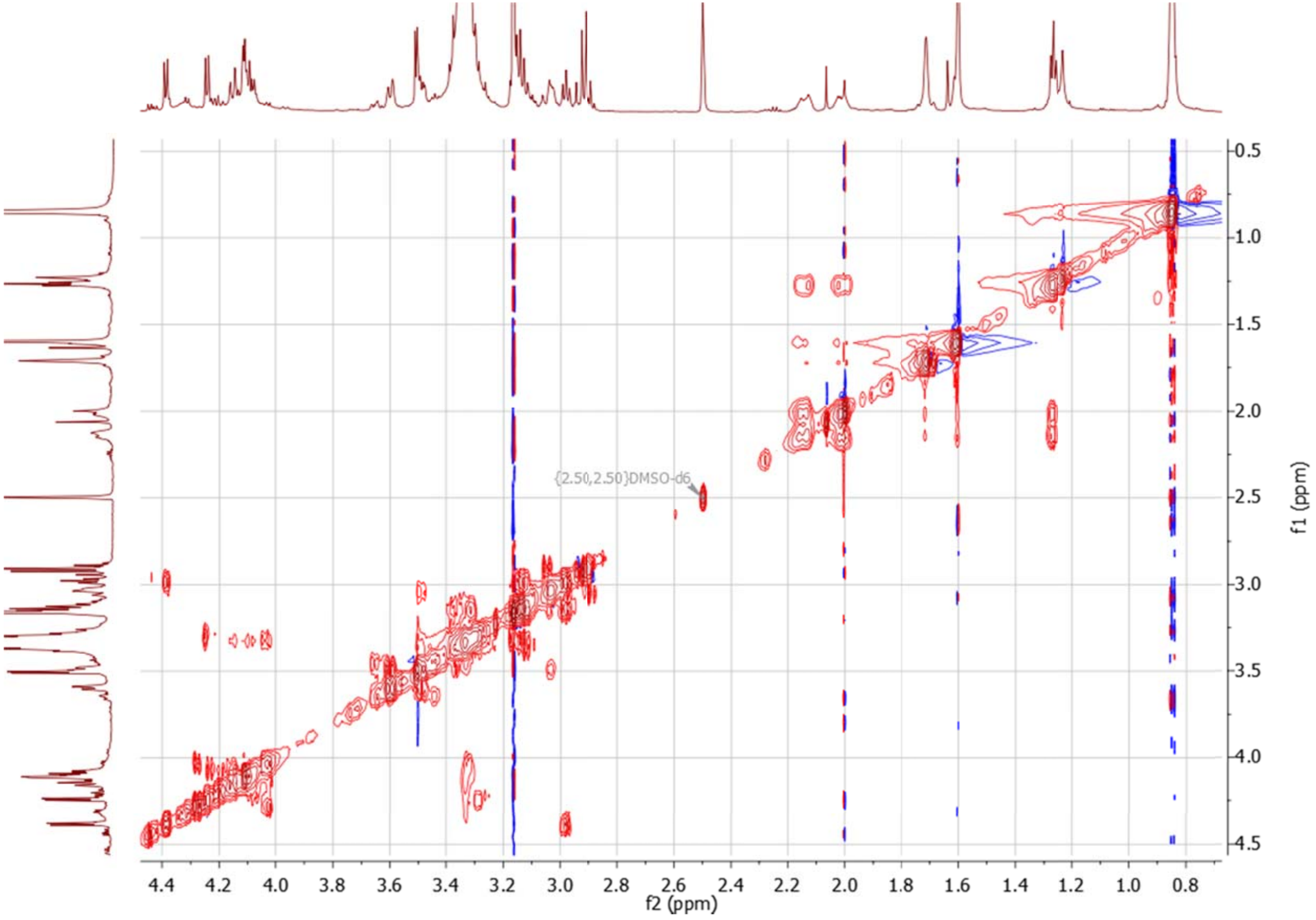

**Figure S2.45** $^{1}H$ -$^{1}H$ CLIP-COSY spectrum of **6** in DMSO-$d_6$.

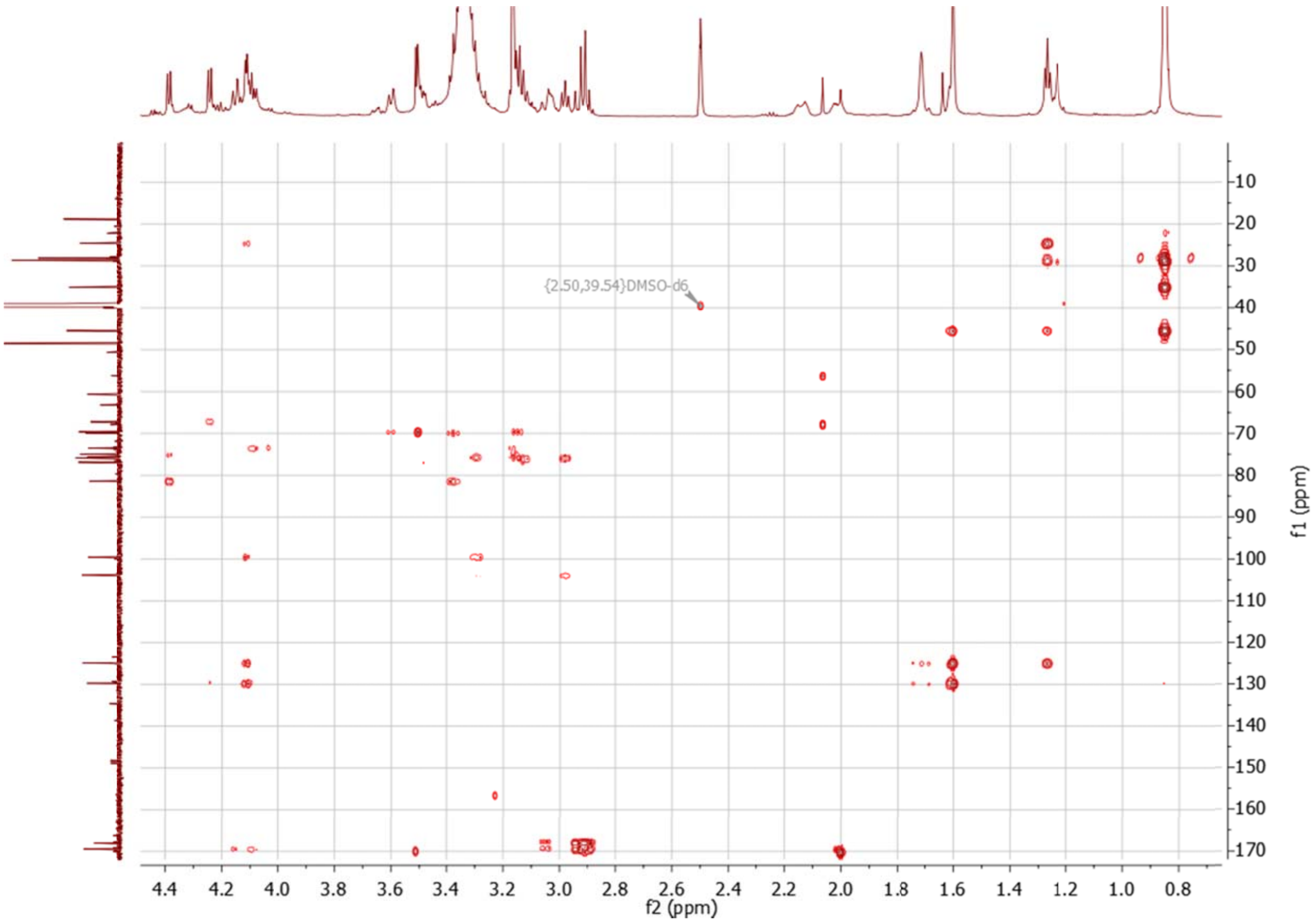

**Figure S2.46** $^{1}H$ -$^{13}C$ HMBC spectrum of **6** in DMSO-$d_6$.

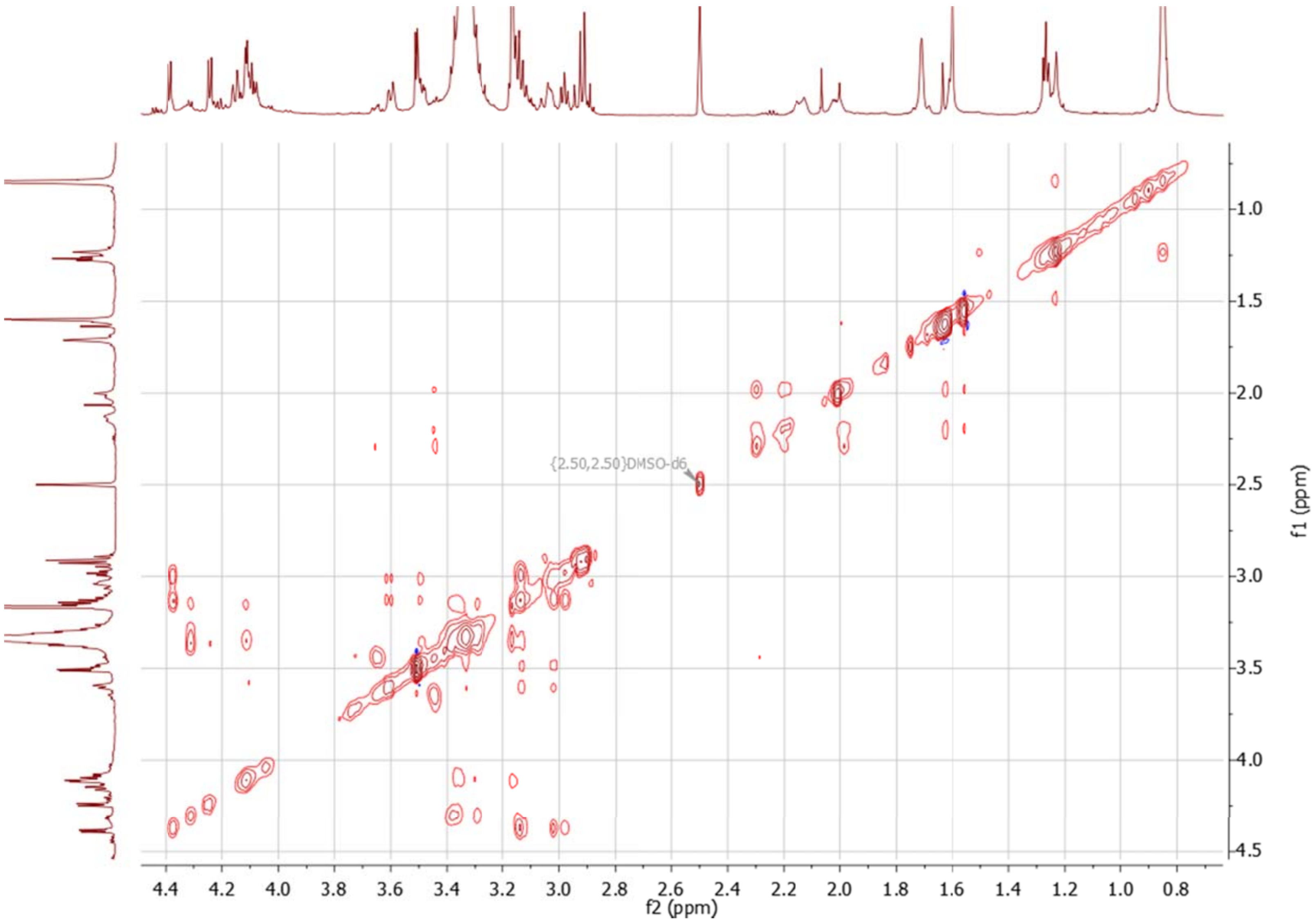

**Figure S2.47** $^{1}H$ -$^{1}H$ TOCSY spectrum of **6** in DMSO-$d_6$.

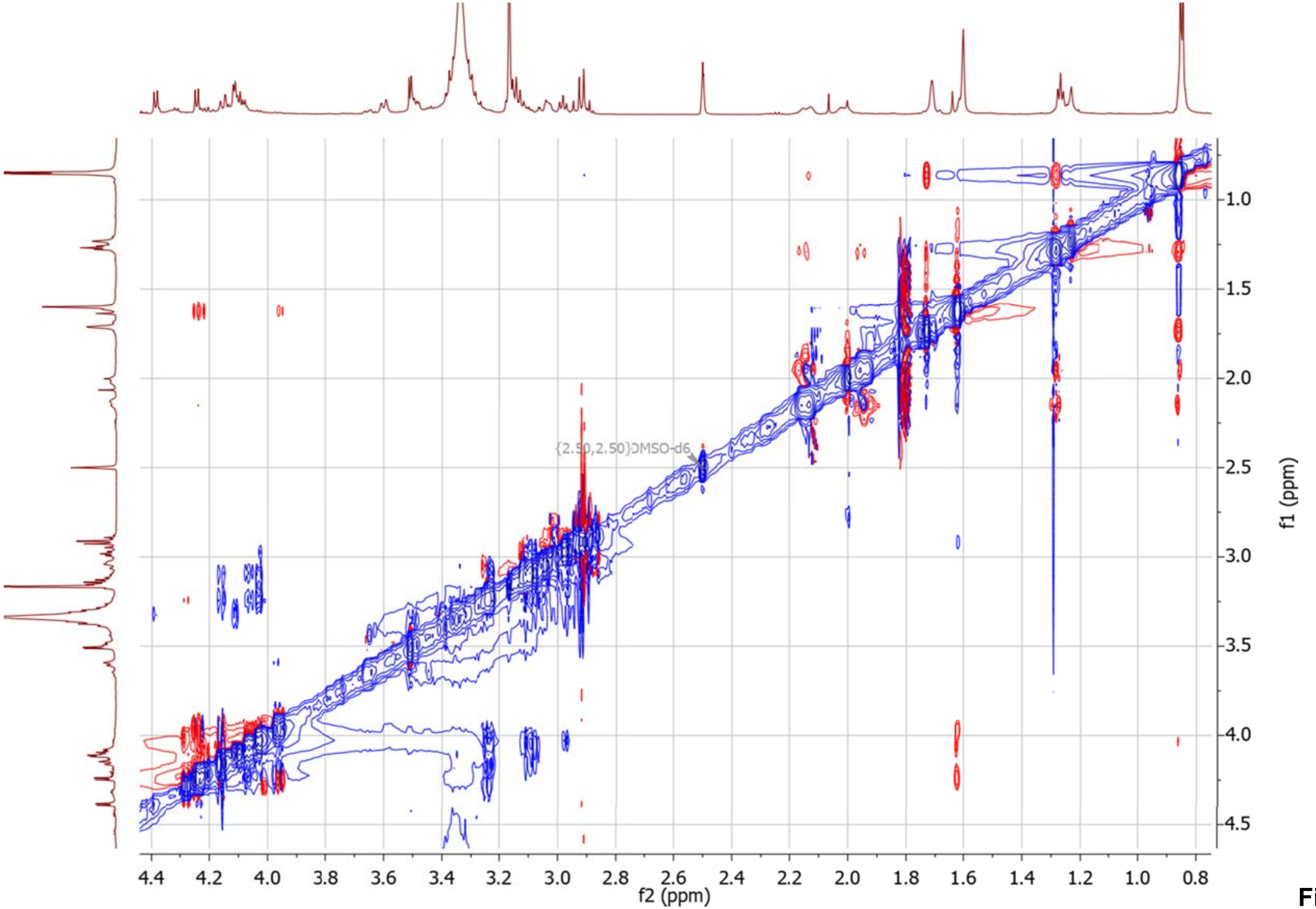

**Figure S2.48** $^1H$ -$^1H$ NOESY spectrum of **6** in DMSO-$d_6$.

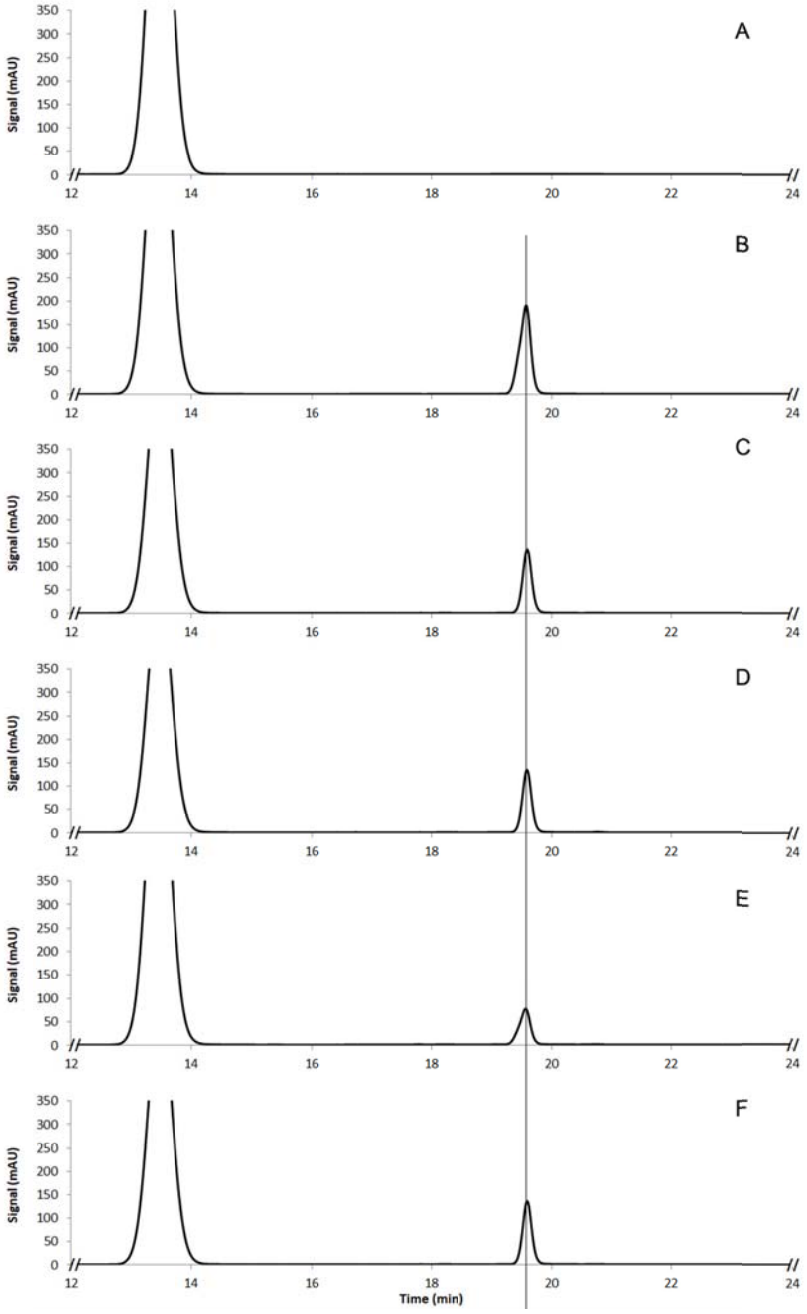

**Figure S2.49** Exemplary HPLC separation of the reaction mixture after MPP-derivatization. The following samples are represented: a) Control sample without the substrate; b) Reference sample of D-glucose; c) Sample containing the hydrolysis product of compound **5**; d) Sample **c** mixed with sample **b** in a 1:1 ratio; e) Sample containing the hydrolysis product of compound **6**; f) Sample **e** mixed with sample **b** in a 1:1 ratio.

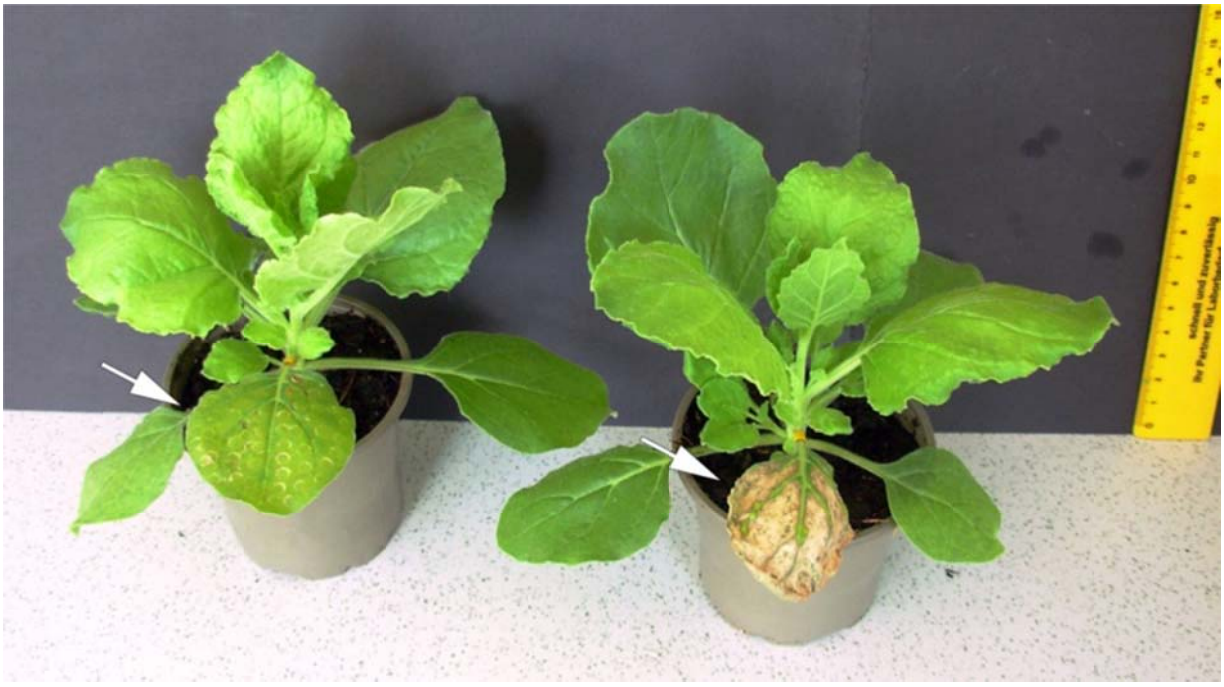

**Figure S2.50** *N. benthamiana* plants expressing StCLDS alone and in combination with tHMGR. The infiltrated leaves are indicated by the arrows.
